# Weak and tunable adhesion-clutch drives rapid cell migration and glioblastoma motility

**DOI:** 10.1101/2025.01.13.632873

**Authors:** Kentarou Baba, Ami Fukushi-Kumagaya, Megumi Morisaki, Ryosuke Takeuchi, Zhize Xiao, Yoshikazu Nagashima, Mizuki Sakai, Yasuna Higashiguchi, Hiroko Katsuno-Kambe, Asako Katsuma, Yoshihiro Ueda, Yuji Kamioka, Daisuke Kawauchi, Tatsuo Kinashi, Yonehiro Kanemura, Naoyuki Inagaki

## Abstract

To move forward, migrating cells must exert backward forces against the extracellular environment. Recent studies have highlighted the importance of integrin-independent forces for cell migration; but the molecular machinery that exerts forces remains unclear. Here, we show that the clutch-linker molecule shootin1 and the cell adhesion molecule L1 transmit the backward force of treadmilling actin filaments to the adhesive environment for rapid dendritic cell migration. Notably, shootin1 and L1 transmit weak traction forces, ∼100 times weaker than integrin-based forces, by constituting an integrin-independent slippery adhesion-clutch. This adhesion-clutch system is tunable in response to the chemoattractant CCL19 and the adhesive ligand laminin and mediates chemotaxis through its polarized activation within cells. Furthermore, its aberrant activity enhances glioblastoma cell motility. Our results show that the weak adhesion-clutch is well-suited for rapid cell migration, without forming strong adhesions that impede cell motility, and provides a potential target for inhibiting abnormal cell motility.

## INTRODUCTION

Cell migration is essential for various biological processes, including immune response, development and regeneration, and requires bidirectional interactions with the extracellular environment. Namely, migrating cells must exert backward forces onto the environment to propel themselves forward; conversely, their speed and direction are regulated in response to various extracellular cues. Dysregulation of these interactions leads to pathogenesis such as cancer cell invasion. Intracellular actin filaments (F-actins) polymerize at the front of migrating cells and disassemble proximally, thereby undergoing backward flow (treadmilling)^1,2^. To explain the force to drive cell migration, the adhesion-clutch model was proposed^3,4^. In this model, clutch-linker molecules (e.g., talin and vinculin) and cell adhesion molecules (e.g., integrins) transmit the movement of F-actin flow to the adhesive environment, thereby exerting backward traction forces that drive cell migration.

Indeed, integrins and the linker molecules, including talin and vinculin, mediate mechanical coupling between F-actin flow and adhesive substrates^5–7^, thereby generating large traction forces that can reach hundreds or kilo Pa^8,9^. However, KO of integrin, talin or vinculin does not always delay three-dimensional (3D) ameboid and two-dimensional (2D) mesenchymal migrations^6,7,10–13^. In contrast to the adhesion-clutch paradigm, integrin activation inhibits T cell motility^10^ and inhibition of the integrin-mediated force transmission by vinculin depletion promotes fibroblast migration^7^, highlighting a role of integrin-independent mechanisms^11,14,15^. Several alternative models have been proposed to explain the backward force exertion, including friction-based^15–19^ and molecular paddling-based^20^ mechanisms. In order to validate the force-exerting models, it is essential to identify the molecules that exert forces and to determine their mechanical dynamics. Thus, despite decades of intensive research, the mechanistic understanding of cell migration remains limited. In addition, it remains unclear how environmental chemical cues direct cell migration through force control^15,21,22^.

Leukocytes undergo rapid ameboid cell migration in tissues; their velocities are up to ∼100 times faster than those of mesenchymal and epithelial cells^11,23^. On the other hand, glioblastoma is the most common and most lethal primary brain tumor in adults with high invasiveness^24,25^. Shootin1a (*SHTN1*) is a neuronal clutch-linker molecule that transmit the force of F-actin flow to the environment for axonal extension^26–28^ and synaptic expansion^29^. We previously reported that shootin1b, a splicing variant of shootin1a, is expressed in the leukocytes, dendritic cells^30^. The present study shows that shootin1b and the cell adhesion molecule L1 form a tunable adhesion-clutch that transmits weak backward forces at levels of 10 Pa and controls the speed and direction of rapid dendritic cell migration under chemical cues. In addition, an aberrant expression of shootin1b in glioblastoma cells promotes their abnormal migration by mediating the weak adhesion-clutch. These results demonstrate that the weak adhesion-clutch enables rapid cell migration and guidance by chemoattractants, and represents a potential target for inhibiting glioblastoma invasion.

## RESULTS

### Weak backward forces propel rapid dendritic cell migration

To investigate the force that propels cell migration, we first monitored the forces generated by migrating dendritic cells using traction force microscopy^31^. Mouse bone marrow-derived dendritic cells in a mixture of collagen gel and Matrigel were plated on laminin-coated polyacrylamide gels with embedded 200-nm fluorescent beads (Figure 1A). Dendritic cells express L1^32^, and Matrigel contains its adhesion ligands laminins^33,34^ which are expressed in the dendritic cell migration pathways^35,36^. Consistent with the previous reports^16,37^, dendritic cells underwent rapid random migration in the presence of 20-200 ng/mL (2.1-21 nM) CCL19 under the semi-3D condition. Traction forces were monitored by force-induced deformation of the polyacrylamide gel, visualized by the bead movement (Video S1, Figure 1B). The cellular front of migrating dendritic cells exerted prominent backward forces on the gel (yellow arrows, Video S1). We also observed centripetal forces at the trailing region (yellow arrows). These data are consistent with the forces generated by migrating dendritic cells detected by micropost arrays in a 2D condition^38^.

**Figure 1.**
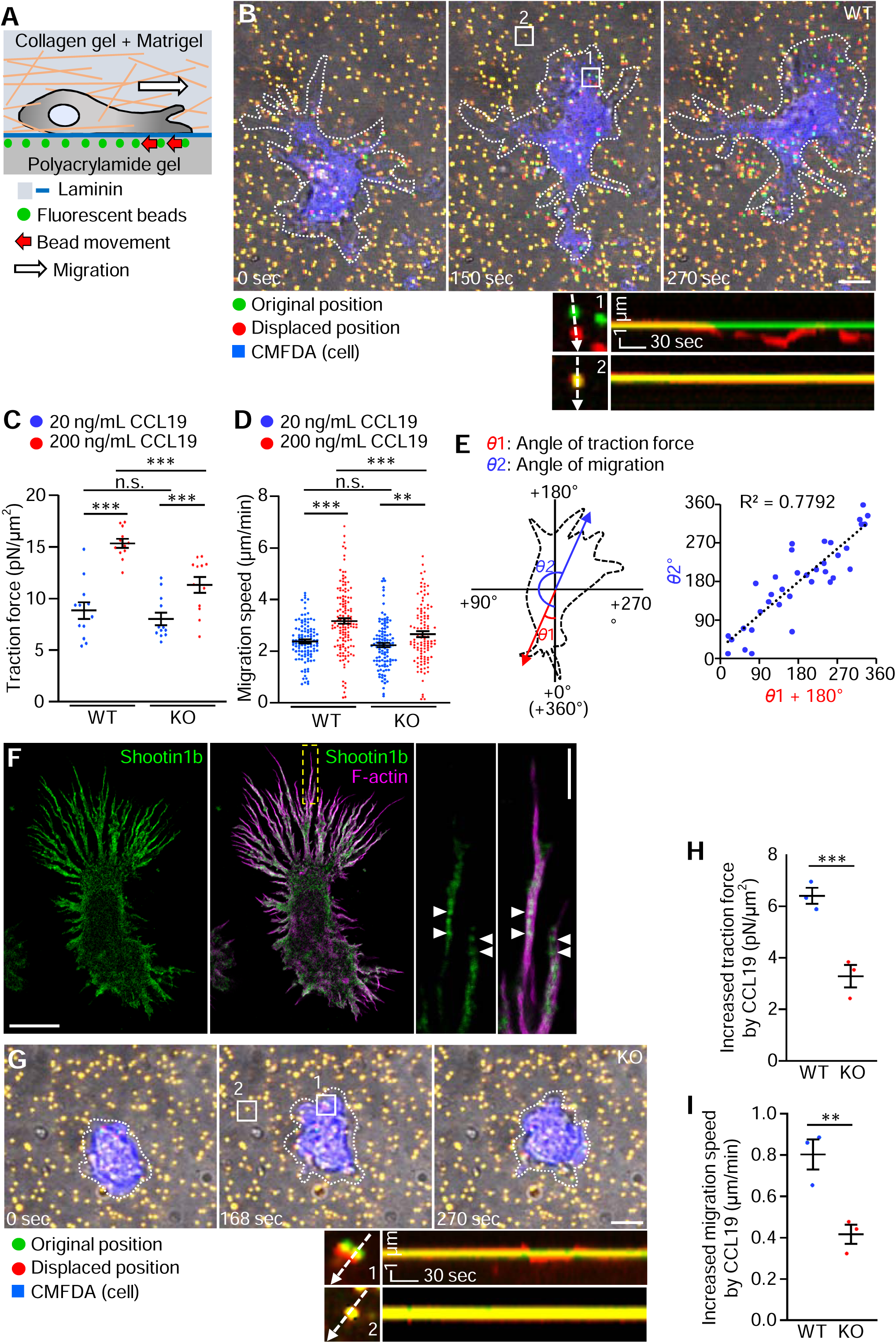
Shootin1b mediates generation of weak forces for dendritic cell migration. (A) Schema of traction force microscopy in a semi-3D condition. Dendritic cells were cultured on laminin-coated polyacrylamide gels embedded with 200-nm fluorescent beads in a mixture of collagen gel and Matrigel. Traction force under the cells was monitored by visualizing force-induced deformation of the gel, which is reflected by the bead movement (red arrows). (B) Overlayed differential interface contrast (DIC) and fluorescence images showing a dendritic cell migrating under the semi-3D condition in (A) in the presence of 200 ng/mL CCL19. See Video S1. The pictures show representative images from the time-lapse series taken every 3 sec for 270 sec. The original and displaced positions of the beads in the gel are indicated by green and red colors, respectively. The cells were visualized by CMFDA staining (blue color); dashed lines indicate the boundaries of the cells. The kymographs (panel below) along the axis of bead displacement (white dashed arrows) at indicated areas 1 and 2 show movement of beads recorded by every 3 sec. The bead in area 2 is a reference bead. Scale bar: 5 µm (in the inset, 1 µm). (C) Analyses of the magnitude of the traction force under WT and shootin1b KO dendritic cells stimulated by 20 ng/mL and 200 ng/mL CCL19. For multiple comparisons, one-way ANOVA with Turkey’s post hoc test was performed. WT + 20 ng/mL CCL19, n = 12 cells; WT + 200 ng/mL CCL19, n = 12 cells; KO + 20 ng/mL CCL19, n = 12 cells; KO + 200 ng/ mL CCL19, n = 12 cells. (D) Analyses of migration speed of WT and shootin1b KO dendritic cells cultured in a mixture of collagen gel and Matrigel in the presence of 20 ng/mL and 200 ng/mL CCL19. For multiple comparisons, one-way ANOVA with Turkey’s post hoc test was performed. WT + 20 ng/ mL CCL19, n = 111 cells; WT + 200 ng/ mL CCL19, n = 137 cells; KO + 20 ng/ mL CCL19, n = 107 cells; KO + 200 ng/ mL CCL19, n = 110 cells. See Figures S1, and Video S2. (E) Scheme showing the angles of traction force (*θ*1) and dendritic cell migration (*θ*2) (left panel). The angles were calculated from the data of sequential 30 images of migrating dendritic cells in the presence of 200 ng/mL CCL19 (B). Right panel shows the correlation analysis between the traction force angle and migration angle of dendritic cells. (F) Fluorescence images of a dendritic cell co-stained with anti-shootin1b antibody and phalloidin-Alexa 555 for F-actin. An enlarged view of the rectangular region is shown to the right. Arrowheads indicate shootin1b co-localization with F-actins in filopodia. The images were obtained by STED microscopy. Scale bar: 10 µm (in the inset, 2 µm). (G) Overlayed DIC and fluorescence images showing a shootin1b KO dendritic cell migrating under the semi-3D condition in (A) in the presence of 200 ng/mL CCL19. For detailed explanations, see (B). See Video S1. (H, I) Effects of shootin1b KO on the traction force (H) and migration speed (I) increased by 200 ng/mL CCL19. Two-tailed unpaired Student′s *t*-test was performed (n = 3 independent experiments). Data represent means ± SEM; **, p < 0.02; ***, p < 0.01; ns, not significant.

The force vectors detected by individual beads under a cell were averaged and expressed as a single vector composed of magnitude and angle (*θ*)^31^. Dendritic cells generated traction force of 8.8 ± 0.8 Pascal (Pa, pN/µm^2^) in the presence of 20 ng/mL CCL19 (Figure 1C). Increasing the CCL19 concentration to 200 ng/mL resulted in a significant increase in the force, reaching 15.4 ± 0.4 Pa (Figure 1C). The amplitude of the force generated by dendritic cells is very weak compared to the traction force produced by focal adhesions (FAs), which can reach hundreds or kilo Pa^8,9^. Importantly, increasing the CCL19 concentration from 20 ng/mL to 200 ng/mL also accelerated random dendritic cell migration under 3D conditions (collagen gel + Matrigel) from 2.4 ± 0.1 μm/min to 3.2 ± 0.1 μm/min (Figures S1A-C, 1D, Video S2). Furthermore, a significant positive correlation was observed between the angle of cell migration and the angle of traction force + 180° (Figure 1E). We therefore conclude that the observed weak backward force propels rapid dendritic cell migration.

### Shootin1b transmits forces for dendritic cell migration under CCL19 signaling

We previously reported that dendritic cells express shootin1b, a splicing variant of shootin1a (Figure S2A-B)^30^. Shootin1a is a neuronal clutch-linker molecule that mediates actin-substrate coupling for axonal extension, through its interaction with the F-actin binding protein cortactin and L1^27,28^. Stimulated emission depletion (STED) microscopy demonstrated that shootin1b colocalizes with F-actins, cortactin, and L1 at the leading edge (Figures 1F, S2C-D). To examine whether shootin1b is involved in the generation of the traction force for dendritic cell migration, we analyzed dendritic cells prepared from shootin1b knockout (KO) mice (Figure S2E-F). Shootin1b KO had no effect on the traction force produced by dendritic cells in the presence of 20 ng/mL CCL19 (Figure 1C). On the other hand, shootin1b KO resulted in 48.6% inhibition of the increase in traction force induced by 200 ng/mL CCL19 (Video S3, Figure 1C, G, H). Expression of a shootin1 dominant-negative mutant (shootin1-DN), which disrupts the interaction between endogenous shootin1 and L1^39^, resulted in a similar reduction in the traction force (Video S4, Figure S3A-C).

Consistently, although shootin1b KO did not affect the migration speed of dendritic cells in the presence of 20 ng/mL CCL19 (Figures S1E, 1D), it led to 48.1% inhibition of the migration speed increased by 200 ng/mL CCL19 (Video S5, Figures S1F, 1D, I). Together, these data indicate that shootin1b promotes forces for dendritic cell migration under CCL19 signaling thorough its interaction with L1. In addition, the parallel correlations between the reductions in traction forces and migration speeds support our conclusion that the weak force drives dendritic cell migration.

### Shootin1b and L1 form a weak and slippery adhesion-clutch enhanced by laminin

To analyze how shootin1b transmits weak forces on the substrate through L1, we monitored their dynamics at the leading edge of dendritic cells. HaloTag-actin, HaloTag-shootin1b or L1-HaloTag expressed in dendritic cells on the glass bottom dishes, coated with the nonspecific adhesive substrate poly-D-lysine (PDL) or subsequently coated with PDL and the extracellular matrix (ECL) protein laminin, were observed by speckle imaging using total internal reflection fluorescence (TIRF) microscopy^31^ in the presence of 200 ng/mL CCL19. F-actins underwent backward movement at the leading edge of dendritic cells (Video S6, Figure 2A) as reported^40^; shootin1b and L1 also moved retrogradely (Video S6, Figure 2B-C). The velocities of F-actins, shootin1b, and L1 were analyzed by tracing the speckles of HaloTag-actin, HaloTag-shootin1b and L1-HaloTag, respectively (yellow lines, Figure 2A-C). Notably, F-actins, shootin1b and L1 moved at similar rates of 1.7 ± 0.1 µm/min, 1.8 ± 0.1 µm/min and 1.8 ± 0.1 µm/min, respectively, on laminin (Figure 2D), suggesting that shootin1b and L1 move by stably interacting with F-actins (Figure 2E) in the presence of 200 ng/mL CCL19. On the other hand, F-actins, shootin1b and L1 showed a similar increase in velocity in the absence of laminin (Video S6, Figure 2D, F-H). Thus, shootin1b and L1 form a weak adhesion-clutch that continuously slips on the adhesive substrate, in which the L1-substrate interphase is enhanced by laminin (Figure 2E).

**Figure 2.**
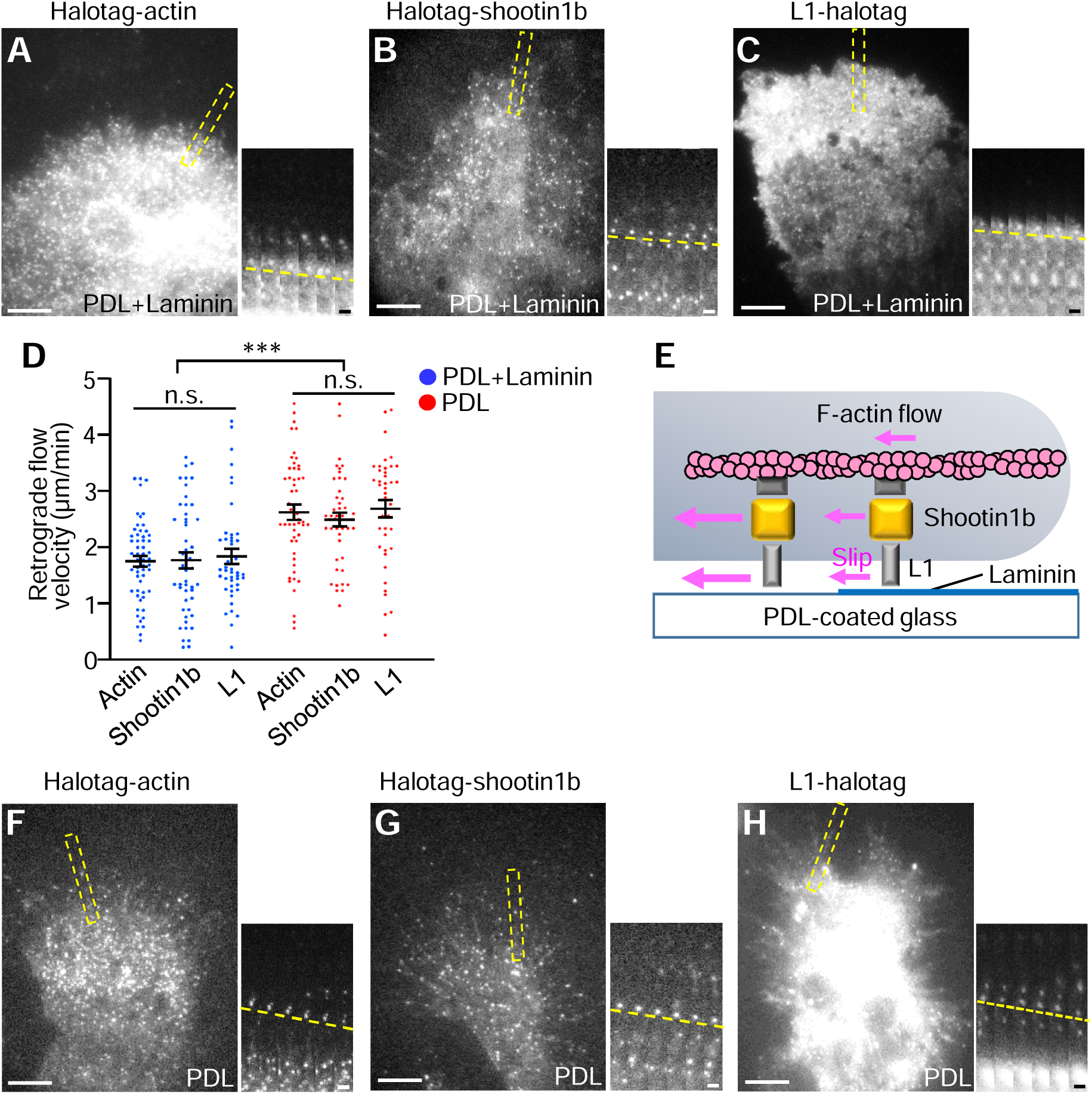
L1 forms a slippery adhesion-clutch enhanced by laminin. (A-C) Fluorescent speckle images of HaloTag-actin (A), Halotag-shootin1b (B) and L1-halotag (C) in dendritic cells cultured on dishes subsequently coated with PDL and laminin under agarose in the presence of 200 ng/mL CCL19. See Video S6. Time-lapse montages of the indicated rectangular regions at 2 sec intervals are shown to the right panel; dashed lines indicate the retrograde flow of speckles. Scale bar: 10 µm (in the inset, 2 µm). (D) Analyses of the retrograde flow speeds of actin, shootin1b and L1 in (A-C and F-H). Two-tailed unpaired Welch’s *t*-test for Halotag-actin between PDL+laminin- and PDL-coated dishes. PDL+laminin, n = 55 speckles from 12 cells; PDL, n = 50 speckles from 9 cells. Two-tailed Mann–Whitney *U*-test for Halotag-shootin1b between PDL+laminin- and PDL-coated dishs. PDL+laminin, n = 48 speckles from 8 cells; PDL, n = 44 speckles from 7 cells. Two-tailed Mann–Whitney *U*-test for L1-halotag between PDL+laminin- and PDL-coated dishs. PDL+laminin, n = 44 speckles from 7 cells; PDL, n = 40 speckles from 6 cells. For multiple comparisons of actin, shootin1b and L1 flow speeds on PDL+laminin-coated dish (blue dots) or PDL-coated dish (red dot), one-way ANOVA with Turkey’s post hoc test was performed. (E) Schema of the L1- and shootin1b-mediated slippery adhesion-clutch enhanced by laminin. (F-H) Fluorescent speckle images of HaloTag-actin (F), Halotag-shootin1b (G) and L1-halotag (H) in dendritic cells cultured on PDL-coated dishes under agarose in the presence of 200 ng/mL CCL19. See Video S6. Time-lapse montages of the indicated rectangular regions at 2 sec intervals are shown to the right panel; dashed lines indicate the retrograde flow of speckles. Scale bar: 10 µm (in the inset, 2 µm). Data represent means ± SEM; ***, p < 0.01; ns, not significant.

### Shootin1b-mediated adhesion-clutch is highly sensitive to CCL19

Stimulation of the CCL19 receptor CCR7 activates Cdc42 and Rac1 and their downstream kinase Pak1^41,42^, which are required for CCL19-induced dendritic cell chemotaxis^37,42^. Pak1 phosphorylates shootin1a at Ser101 and Ser249 in neurons^43^, which in turn enhances adhesion-clutch by promoting shootin1a-cortactin and shootin1a-L1 interactions^28,39^; shootin1b also contains Ser101 and Ser249^30^. To investigate how shootin1b promotes dendritic cell migration under CCL19 signaling, we examined shootin1b phosphorylation in dendritic cells. In the absence of CCL19, low levels of shootin1b phosphorylations at Ser101 and Ser249 were detected (Figure 3A). The phosphorylations increased markedly in a dose- and time-dependent manner, reaching saturation at 200 ng/mL CCL19 (Figure 3A-B) and 10 min (Figure S4A-B), respectively. They were inhibited by the Pak1 inhibitor NVS-PAK1-1 (Figure S4C-D), indicating that CCL19 triggers Pak1-mediated shootin1b phosphorylation in dendritic cells. Furthermore, immunoprecipitation analyses showed that the amounts of L1 and cortactin co-precipitated with shootin1b were strongly increased by increasing the concentration of CCL19 concentration from 0 ng/mL to 200 ng/mL (Figure 3C-D), indicating that the shootin1b-L1 and shootin1b-cortactin interactions are highly sensitive to CCL19 concentration.

**Figure 3.**
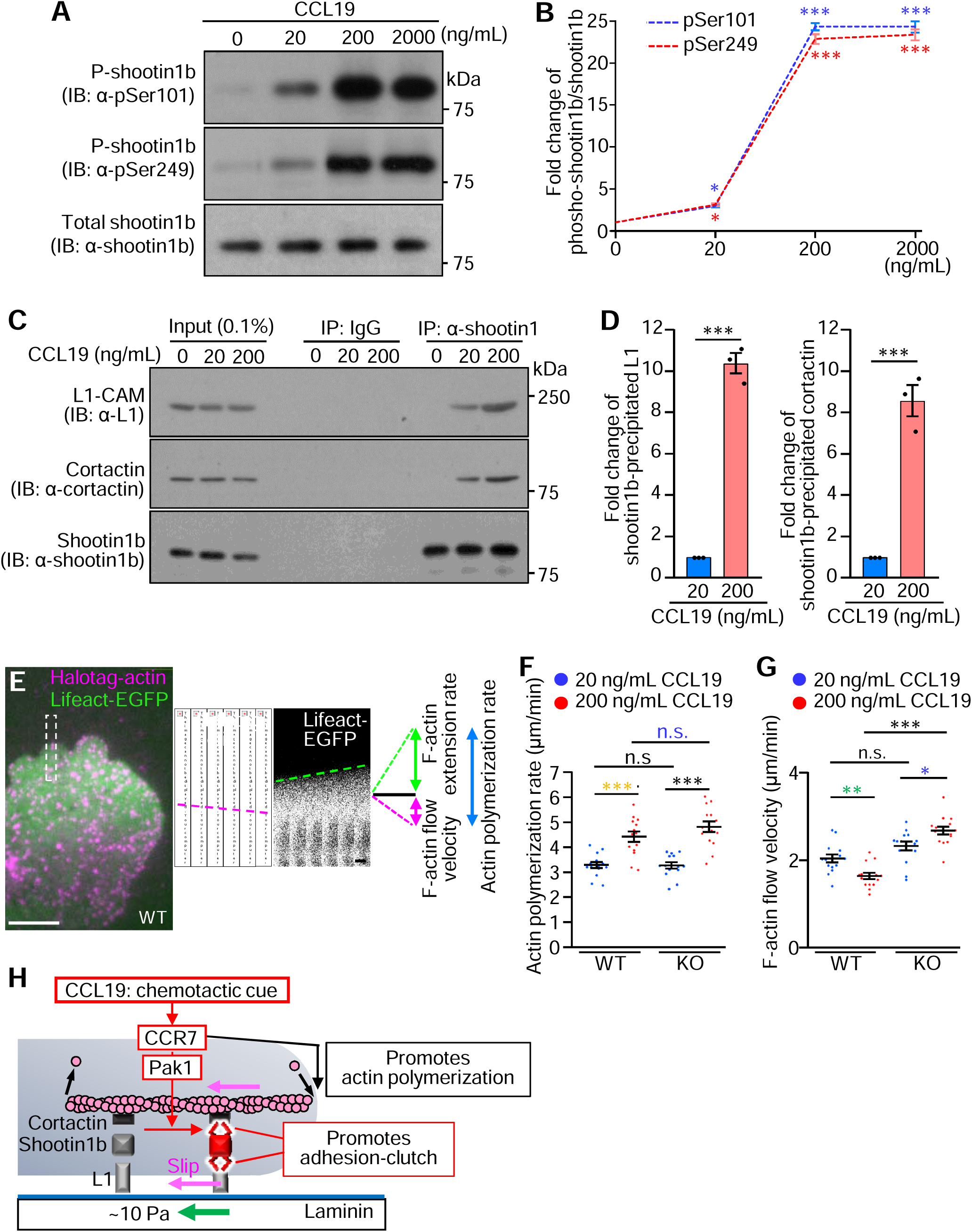
CCL19 promotes shootin1b-mediated adhesion-clutch. (A) Dendritic cells were treated with 0, 20, 200 or 2000 ng/mL CCL19 for 30 min. Cell lysates were then analyzed by immunoblot with anti-pSer101 shootin1, anti-pSer249 shootin1 and anti-shootin1b antibodies. (B) Fold change of the phosphorylated shootin1b levels at Ser101 and Ser249 normalized by total shootin1b in (A). One-way ANOVA with Turkey’s post hoc test was performed for multiple comparison of the data (n = 3 independent experiments). (C) Co-immunoprecipitation of L1 and cortactin with shootin1b in dendritic cells. After incubation of dendritic cells with 20 ng/mL, 200 ng/mL CCL19, or medium (0 ng/mL CCL19) for 30 min, cell lysates were prepared and incubated with anti-shootin1 antibody or isotype control IgG for immunoprecipitation. The immunoprecipitates and cell lysates (0.1 %) were immunoblotted with anti-shootin1b, anti-L1, and anti-cortactin antibodies. (D) Quantitative data for L1 and cortactin co-precipitated with shootin1b in (C). Two-tailed unpaired Welch’s *t*-test was performed (n = 3 independent experiments). (E) Overlayed fluorescent speckle image of HaloTag-actin and fluorescence image of Lifeact-EGFP at the leading edge of a dendritic cell on laminin-coated dishes under agarose in the presence of 200 ng/mL CCL19. See Video S7. Time-lapse montages of the indicated rectangular region at 2 sec intervals are shown to the right panel. Magenta and green dashed lines indicate HaloTag-actin retrograde flow and F-actin extension, respectively. The actin polymerization rate (blue double-headed arrow) was calculated as the sum of the F-actin retrograde flow and extension rates. Scale bar: 10 µm (in the inset, 2 µm). (F, G) Analyses of actin polymerization rate (F) and F-actin flow speed (G) in dendritic cells stimulated by 20 ng/mL or 200 ng/mL CCL19. One-way ANOVA with Turkey’s post hoc test was performed for multiple comparisons. WT + 20 ng/mL CCL19, n = 14 cells; WT + 200 ng/mL CCL19, n = 14 cells; KO + 20 ng/mL CCL19, n = 14 cells; KO + 200 ng/mL CCL19, n = 14 cells. (H) Schema of CCL19-induced dual enhancement of shootin1b-mediated adhesion-clutch and actin polymerization. Data represent means ± SEM; *, p < 0.05; **, p < 0.02; ***, p < 0.01; ns, not significant.

We further monitored the adhesion-clutch dynamics under CCL19 stimulation. Dendritic cells were transfected with HaloTag-actin and Lifeact-EGFP and cultured on laminin-coated dishes for observation (Video S7, Figure 3E). F-actin flow velocity was analyzed by tracing the speckles of HaloTag-actin (magenta double-headed arrow, Figure 3E), while actin polymerization rate was calculated as the sum (blue double-headed arrow) of F-actin protrusion rate monitored by Lifeact-EGFP (green double-headed arrow) and the F-actin flow velocity (magenta double-headed arrow)^31^. Actin polymerization rate increased with an increase in CCL19 concentration from 20 ng/mL to 200 ng/mL (yellow asterisks in Figure 3F), consistent with a previous report using T cells^16^.

Importantly, despite the increased actin polymerization, F-actin flow velocity decreased by 200 ng/mL CCL19 (green asterisks in Figure 3G). The decreased F-actin flow velocity and the increased traction force (Figure 1C) are the key indicators of increased adhesion-clutch^6,7,43^. In shootin1b KO dendritic cells, the increase in CCL19 concentration from 20 ng/mL to 200 ng/mL increased the F-actin flow velocity (blue asterisk in Figure 3G), indicating that CCL19 enhances shootin1b-mediated adhesion-clutch. On the other hand, shootin1b KO did not affect the increased actin polymerization induced by 200 ng/mL CCL19 (blue n.s., Figure 3F). Thus, CCL19 signaling leads to a dual enhancement of shootin1b-mediated adhesion-clutch and actin polymerization at the leading edge (Figure 3H).

### Shootin1b drives CCL19-induced chemotaxis through polarized activation

Dendritic cells migrate under CCL19 gradients *in vivo* from various tissues, including the skin, to lymph nodes through lymphatic vessels^44–46^. Next, we analyzed dendritic cell chemotaxis in 3D conditions (collagen gel + Matrigel) under CCL19 gradients. Dendritic cells underwent chemotaxis toward the CCL19 source (Video S8, Figure 4A) as reported^47^; on the other hand, shootin1b KO partially inhibited dendritic cell migration toward the CCL19 source (Video S8, Figure 4B). To further analyze the effects of shootin1b KO, we calculated the chemotaxis index (straight distance toward the CCL19 source/total distance) by tracing the migration trajectories (Figure 4C). The mean migration speed and chemotaxis index of control dendritic cells were 4.4 ± 0.2 µm/min and 0.5 ± 0.02, respectively (blue dots, Figure 4D-E). Deletion of shootin1b not only reduced the migration speed (red dots, Figure 4D), but also decreased the chemotaxis index (red dots, Figure 4E). Similarly, expression of shootin1-DN reduced the migration speed and chemotaxis index (Video S8, Figure S5). Together, these data indicate that the shootin1b- and L1-mediated adhesion-clutch drives CCL19-induced dendritic cell chemotaxis.

**Figure 4.**
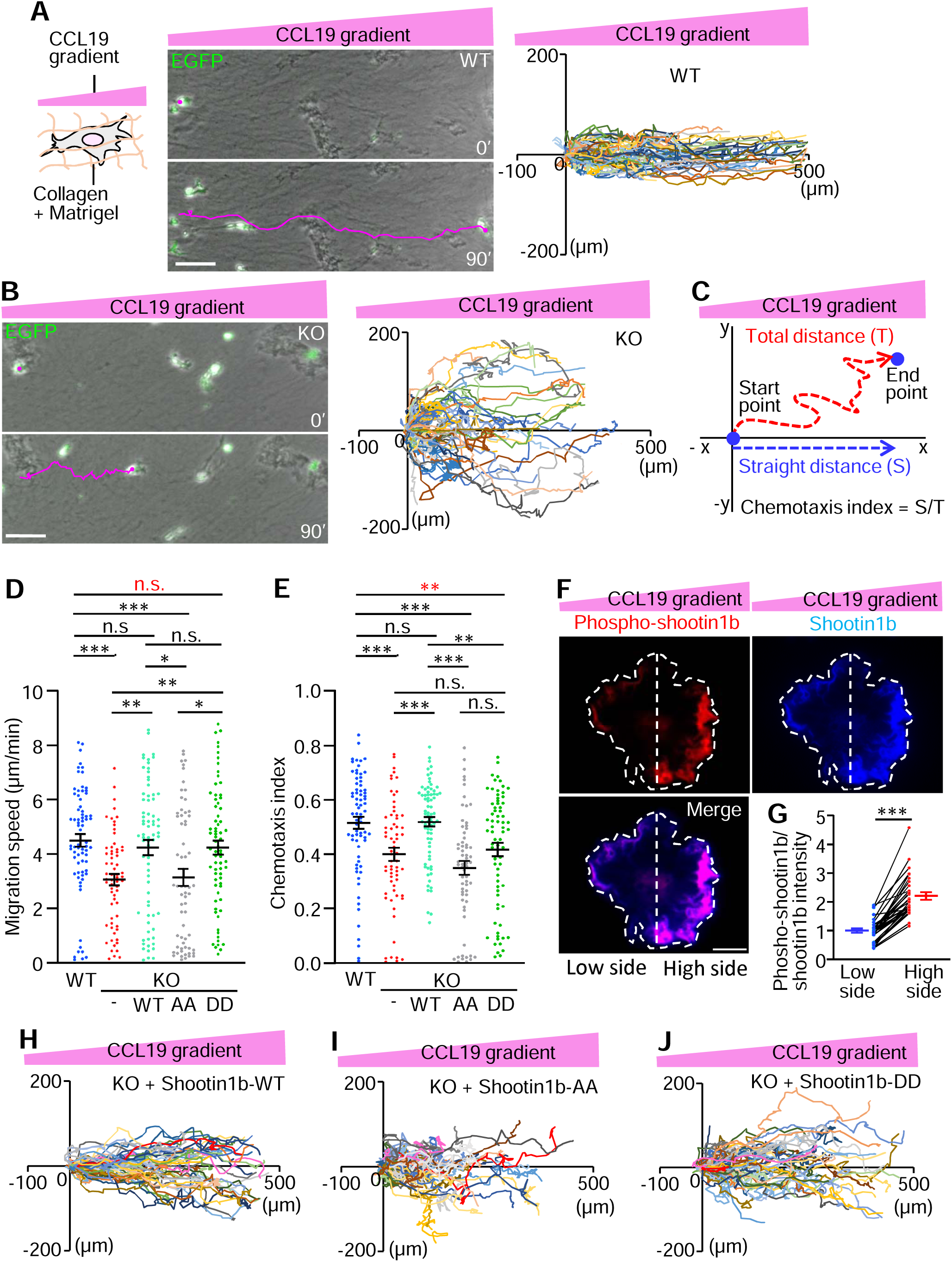
CCL19 gradient induces asymmetric shootin1b activation to drive chemotaxis. (A, B) A gradient of CCL19 was applied to dendritic cells cultured in a mixture of collagen gel and Matrigel (left panel, A). One hour after gradient application, time-lapse phase-contrast/fluorescence images of WT (A) and shootin1b KO (B) dendritic cells expressing EGFP were obtained. Nuclei were also visualized by Hoechst to accurately trace the trajectories of cell migrations (shown in right panels and Video S8). The pictures show representative images from the time-lapse series taken every 1 min for 90 min. The tracing lines (magenta) indicate dendritic cell migration for 90 min. Scale bar: 50 µm. The right panels depict trajectories of dendritic cell migrations. The initial cell positions are normalized at x = 0 µm and y = 0 µm. (C) Scheme of chemotaxis index. Chemotaxis index was calculated as the ratio of the straight distance toward the CCL19 source (*S*) to the total distance (*T*) by tracing the migration trajectories in (A, B). (D, E) Analyses of migration speed (D) and chemotaxis index (E) of WT and shootin1b KO dendritic cells expressing flag-GST (control flag-tagged protein), and KO cells expressing flag-shootin1b-WT (H), flag-shootin1b-AA (I) and flag-shootin1b-DD (J) under the CCL19 gradient. For multiple comparison, one-way ANOVA with Turkey’s post hoc test was performed. WT, n = 72 cells; KO, n = 65 cells; KO + flag-shootin1b-WT, n = 73 cells; KO + flag-shootin1b-DD, n = 72 cells; KO + flag-shootin1b-AA, n = 64 cells. See Video S8. (F) Dendritic cells were transfected with myc-shootin1b to visualize shootin1b. After the stimulation by CCL19 gradients for 30 min, they were fixed and immunolabeled with anti-myc and anti-pSer249 shootin1 antibodies. Fluorescence images show the detected phosphorylated shootin1b and shootin1b in a dendritic cell. White dashed lines indicate the boundary of a dendritic cell and the center line that separates the high side (CCL19 source side) and low side. Scale bar: 10 µm. (G) Quantitative data for shootin1b activation (phospho-shootin1b/shootin1b) in the CCL19 source side (high side) and low side of dendritic cells. Two-tailed Mann–Whitney *U*-test for phospho-shootin1b/total shootin1b between the high side and low side (n = 30 cells). (H, I, J) Migration trajectories of shootin1b KO cells expressing flag-shootin1b-WT (H), flag-shootin1b-AA (I) and flag-shootin1b-DD (J) stimulated by the CCL19 gradient. Data represent means ± SEM; *, p < 0.05; **, p < 0.02; ***, p < 0.01; ns, not significant.

We further examined shootin1b activation under a CCL19 gradient. A CCL19 gradient was applied to dendritic cells expressing myc-shootin1b for 30 min. They were then fixed and immunolabeled with an antibody that recognizes shootin1b phosphorylation at Ser249 (red, Figure 4F). Localization of shootin1b (phosphorylated and unphosphorylated shootin1b) was visualized with an anti-myc antibody (blue). As shown in Figure 4F, polarized localization of the phosphor-shootin1b was observed within dendritic cells. The relative level of shootin1b activation (phospho-shootin1b/shootin1b) was 120% higher on the CCL19 high side than on the low side (Figure 4G).

To examine a role of the polarized shootin1b phosphorylation (Figure 4F-G), we expressed wild-type shootin1b (shootin1b-WT), unphosphorylated shootin1b mutant (shootin1b-AA), in which Ser101 and Ser249 are replaced by alanine, and phosphomimic shootin1b mutant (shootin1b-DD), in which these residues are replaced by aspartate, in shootin1b KO dendritic cells. As expected, expression of shootin1b-WT, but not shootin1b-AA, rescued the reductions in both migration speed and chemotaxis index of shootin1b KO dendritic cells (Figure 4D-E, H-I). On the other hand, shootin1b-DD rescued the reduction in migration speed (red n.s., Figure 4D, J) but not the reduction in chemotaxis index (red asterisks, Figure 4E). Since shootin1b-DD is constitutively active and cannot be regulated under the CCL19 signaling, we conclude that the polarized shootin1b activation within dendritic cells drives the directional chemotaxis induced by CCL19 gradients.

### Shootin1b-and L1-mediated dendritic cell migration is laminin-sensitive and integrin-independent

As shootin1b- and L1-mediated adhesion-clutch is enhanced by laminin (Figure 2E), we next analyzed a role of laminins in chemotaxis. Removal of Matrigel, that contains laminins, from the 3D environment decreased both the migration speed and chemotaxis index of CCL19-induced dendritic cell chemotaxis (Figure 5A-C).

**Figure 5.**
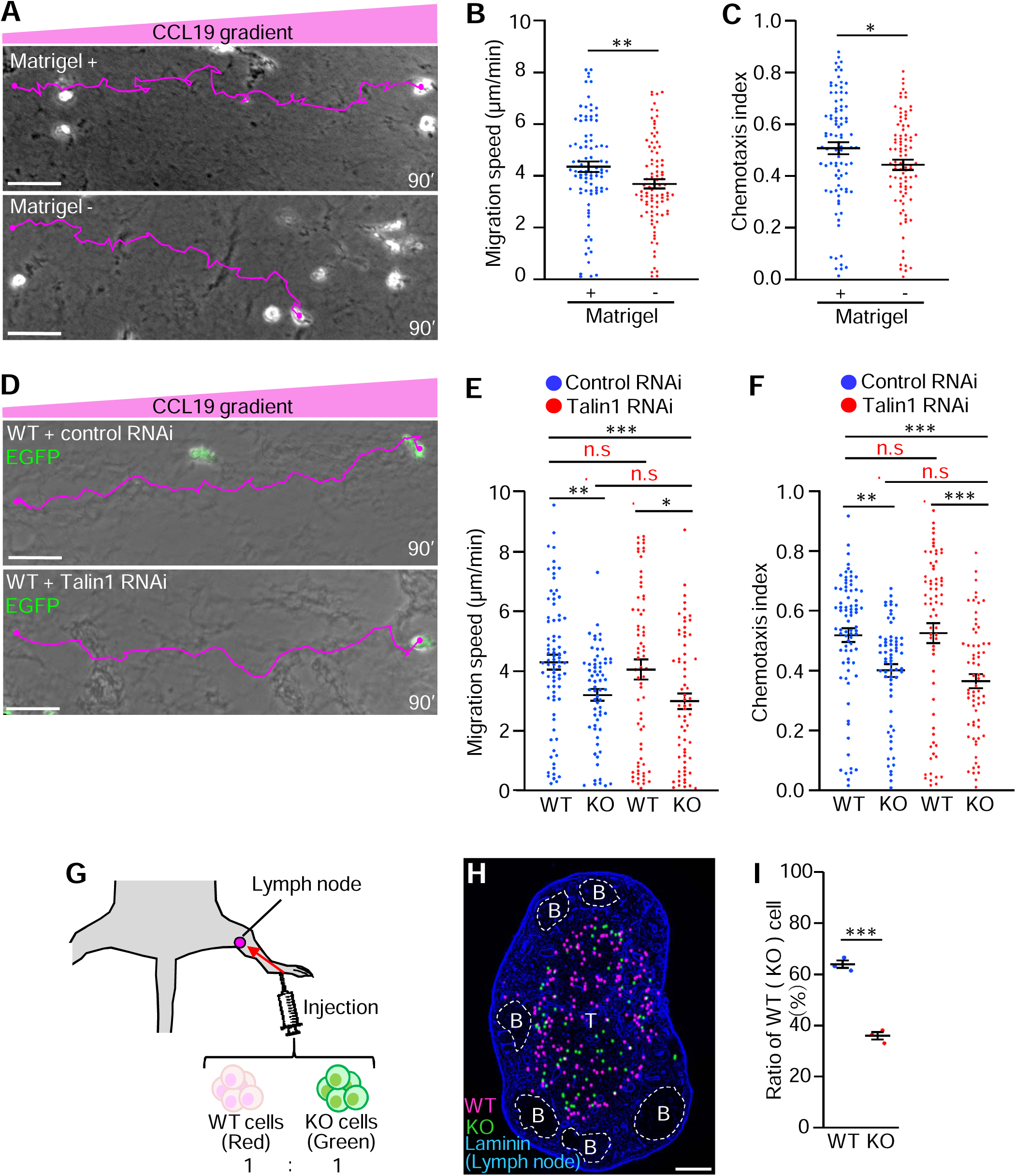
Matrigel-dependent dendritic cell chemotaxis and Shootin1b mediates talin-independent chemotaxis *in vitro* and chemotaxis *in vivo*. (A) Dendritic cells were cultured in a mixture of collagen gel and Matrigel, that contains laminin, (Matrigel +) or without Matrigel (Matrigel -). One hour after application of the CCL19 gradient, time-lapse phase-contrast/fluorescence images of dendritic cells were obtained. Nuclei were also visualized by Hoechst to accurately trace the trajectories of cell migrations. The pictures show representative images from the time-lapse series taken every 1 min for 90 min. Tracing line (magenta) indicate dendritic cell migration for 90 min. Scale bar: 50 µm. (B, C) Analyses of the migration speed (B) and chemotaxis index (C) of dendritic cells in (A). Two-tailed unpaired Student’s *t*-test was performed (Matrigel +, n = 89 cells; Matrigel –, n = 91 cells). (D) Dendritic cells expressing control RNAi or talin1 RNAi were cultured in a mixture of collagen gel and Matrigel. One hour after application of the CCL19 gradient, time-lapse phase-contrast/fluorescence images of dendritic cells were obtained. Nuclei were also visualized by Hoechst to trace the cell migrations (see also Video S9). Tracing line indicate dendritic cell migration for 90 min. Scale bar: 50 µm. (E, F) Analyses of migration speed (E) and chemotaxis index (F) of WT and shootin1b KO dendritic cells expressing control RNAi or talin1 RNAi vector in (D). One-way ANOVA with Turkey’s post hoc test was performed for multiple comparisons. WT + control RNAi, n = 79 cells; KO + control RNAi, n = 65 cells; WT + talin1 RNAi, n = 68 cells; KO + talin1 RNAi, n = 67 cells. (G) Scheme of the *in vivo* dendritic cell migration assay. WT and shootin1b KO dendritic cells were labeled by CMTPX (red) and CMFDA (green), respectively, and were mixed in a 1:1 ratio in suspension. The suspension was injected into the foot pad. Twenty-four hours after the injection, the popliteal lymph nodes were removed and the cells that arrived in the lymph nodes were analyzed (H). (H) Fluorescence images of WT (magenta) and shootin1b KO (green) cells migrated into a lymph node. The lymph node was visualized with anti-pan-laminin antibody. B, B-cell follicles; T, T-cell cortex. Scale bar: 200 µm. (I) Analyses of the ratio of the number of WT and shootin1b KO dendritic cells migrated into the lymph nodes. Two-tailed unpaired Student’s *t*-test was performed (n = 3 independent experiments and 3 lymph nodes). Data represent means ± SEM; *, p < 0.05; **, p < 0.02; ***, p < 0.01; ns, not significant.

Talin couples F-actins with the cell adhesion molecules integrins as a clutch-linker molecule^6,48^. However, shootin1b did not interact with β1- or β2-integrin (Figure S6A) which are expressed in dendritic cells^40^. *Talin* has two isoforms, talin1 and talin2; talin2 is not detected in dendritic cells^49^. Talin1 knockdown (Figure S6B) did not affect the migration speed or chemotaxis index of WT and shootin1b KO dendritic cells under CCL19 gradients (Video S9, Figure 5D-F), consistent with the previous report that integrin ablation does not affect dendritic cell migration *in vitro* and *in vivo*^11^. These data suggest that shootin1b and L1 mediate laminin-sensitive and integrin-independent dendritic cell migration.

### Shootin1b drives dendritic cell migration in tissues

Next, we examined whether shootin1b mediates the dendritic cell chemotaxis from the skin to lymph nodes in response to CCL19 gradients^44–46^. WT dendritic cells labelled with CMTPX and shootin1b KO dendritic cells labelled with CMFDA were mixed in a 1 : 1 ratio, and injected into the mouse footpad (Figure 5G). Twenty-four hours following the injection, the popliteal lymph nodes were excised, and the number of WT and shootin1b KO dendritic cells that had migrated into the lymph nodes was analyzed (Figure 5H). The ratio of shootin1b KO dendritic cells that migrated into lymph nodes was significantly lower than that of WT dendritic cells (Figure 5I), indicating that shootin1b KO inhibits dendritic cell migration into the lymph nodes. To further examine the migration of dendritic cells within the lymph nodes, CMFDA-labeled WT dendritic cells and CMTPX-labeled shootin1b KO dendritic cells were mixed in a 1 : 1 ratio, and placed on the lymph node slices. After 1h, dendritic cells migrated in the slices were observed by confocal microscopy (Figure S7A, Video S10). Shootin1b KO led to a significant decrease in migration speed of dendritic cells within the slices (Figure S7B). Collectively, these results indicate that shootin1b drives dendritic cell migration in tissues.

### Aberrant activity of shootin1b promotes glioblastoma cell motility

Finally, we investigated the motility of highly invasive glioblastoma cells. Although shootin1b expression was undetectable in normal human and mouse brain-derived astrocytes, immunoblot analysis detected a significant amount of shootin1b in glioblastoma cells derived from a human patient (KNBTG-8) (Figure 6A). In addition, L1 and cortactin, but do not shootin1a, was detected in these cells (Figure S8A). During their invasion within the brain tissue, glioblastoma cells extend long protrusions called tumor microtubes at the front^50,51^. We observed similar protrusions at the front of glioblastoma cells in culture (Figure 6B). Shootin1b colocalized with F-actins at the tip of the tumor microtubes (arrowheads, Figure 6B). Their migration speed in 3D Matrigel was 0.16 ± 0.01 µm/min and 3.7 times faster than that of normal human astrocytes (Figure 6C-D, Video S11).

**Figure 6.**
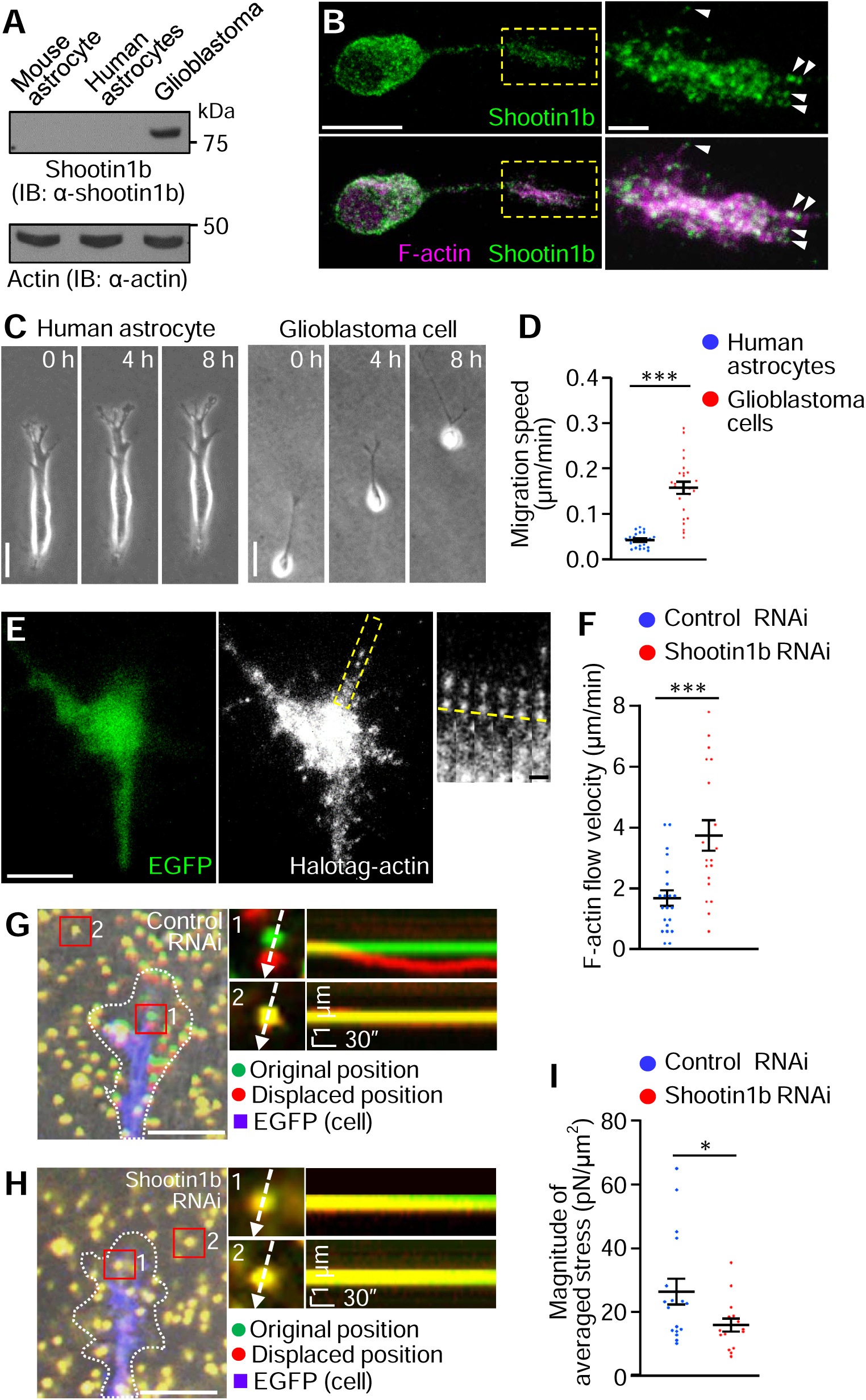
Shootin1b mediates weak adhesion-clutch at the leading edge of glioblastoma cells. (A) Immunoblot analysis of mouse astrocytes, human astrocytes, and glioblastoma cells derived from a human patient (KNBTG-8) with anti-shootin1b antibody. Actin was used as a loading control. (B) Fluorescence images of a human glioblastoma cell co-stained with anti-shootin1b antibody and phalloidin-Alexa 555 for F-actin. An enlarged view of the rectangular region is shown to the right. Arrowheads indicate shootin1b co-localization with F-actins at the tip of the tumor microtube^50,51^. The images were obtained by STED microscopy. Scale bar: 10 µm (in the inset, 2 µm). (C) Human astrocytes and human glioblastoma cells were cultured in Matrigel, and time-lapse phase-contrast images were obtained. The pictures show representative images from the time-lapse series taken every 10 min for 8 hours. Scale bar: 30 µm. See Video S11. (D) Analysis of the migration speeds of human astrocytes and human glioblastoma cells in (C). Two-tailed unpaired Welch’s *t*-test was performed (human astrocytes, n = 20; human glioblastoma cells, n = 24 cells). (E) Fluorescent speckle images of HaloTag-actin at the tip of the tumor microtube of a glioblastoma cell expressing control RNAi vector on a laminin-coated dish. The expression of control RNAi or shootin1b RNAi vector was detected by EGFP. See Video S12. Time-lapse montages of the indicated rectangular regions at 5 sec intervals are shown to the right. Yellow dashed lines indicate HaloTag-actin retrograde flow. Scale bar: 10 µm (in the inset, 2 µm). (F) Analyses of F-actin flow speed. Two-tailed unpaired Welch’s *t*-test was performed (control RNAi, 21 speckles from 19 cells; shootin1b RNAi, 19 speckles from 14 cells). (G, H) Overlayed DIC and fluorescence images showing the tip of the tumor microtube of glioblastoma cells expressing control RNAi (G) and shootin1b RNAi (H) cultured on laminin-coated polyacrylamide gels embedded with 200-nm fluorescent beads. See Video S13. The pictures show representative images from the time-lapse series taken every 3 sec for 180 sec. The original and displaced positions of the beads in polyacrylamide gel are indicated by green and red colors, respectively. The tip of the tumor microtube were visualized by EGFP (blue color); dashed lines indicate the boundaries of the cells. The kymographs (right panels) along the axis of bead displacement (white dashed arrows) at indicated areas 1 and 2 show the movement of beads recorded by every 3 sec. The beads in area 2 are reference bead. Scale bar: 5 µm (in the inset, 1 µm). (I) Analyses of the magnitude of the traction force under the tip of the tumor microtube of glioblastoma cells expressing control RNAi or shootin1b RNAi vector. Two-tailed unpaired Welch’s *t*-test was performed (control RNAi, n = 17 cells; shootin1b RNAi, n = 15 cells). Data represent means ± SEM; *, p < 0.05; ***, p < 0.01.

In control glioblastoma cells, F-actins moved retrogradely at 1.7 ± 0.3 µm/min at the tip of the tumor microtubes (Figure 6E-F, Video S12), resulting in the generation of a traction force of 26.4 ± 4.1 Pa there (Figure 6G, I, Video S13). Shootin1b knockdown (Figure S8B) accelerated the F-actin flow (Figure 6F, Video S12) and led to 39% reduction in the traction force (Figure 6H-I, Video S13), indicating that shootin1b mediates a weak adhesion-clutch at the tip of the tumor microtubes. Furthermore, shootin1b knockdown and shootin1-DN expression inhibited the migration speed of glioblastoma cells by 43% and 40%, respectively (Figure 7A-F, Video S11).

**Figure 7.**
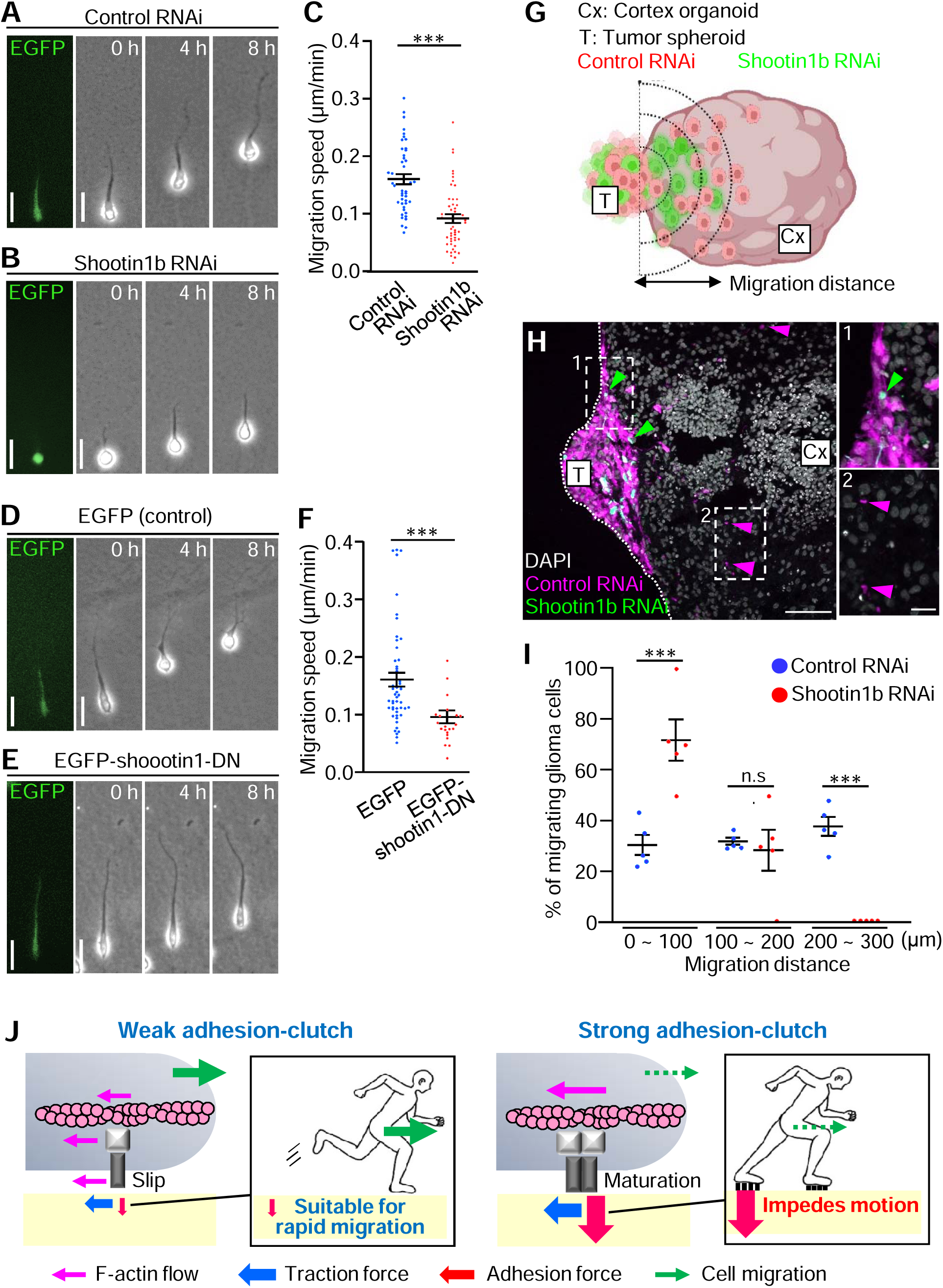
Aberrant activity of shootin1b promotes glioblastoma cell motility. (A, B, D, E) Human glioblastoma cells were cultured in Matrigel, and time-lapse phase-contrast images of the cells expressing control RNAi (A) or shootin1b RNAi (B) vector and overexpressing EGFP (control) (D) or EGFP-shootin1-DN (E) were obtained. The expression of control RNAi or shootin1b RNAi vector was detected by EGFP (left panel). The pictures show representative images from the time-lapse series taken every 10 min for 8 hours. See Video S11. Scale bar: 30 µm. (C, F) Analyses of migration speed of human glioblastoma cells expressing control RNAi or shootin1b RNAi vector (C) and overexpressing EGFP or EGFP-shootin1-DN (F). Two-tailed Mann-Whitney *U*-test was performed for the migration speed of cells expressing control RNAi or shootin1b RNAi (C). Two-tailed Mann-Whitney *U*-test was performed for the migration speed of cells overexpressing EGFP or EGFP-shootin1-DN (F) (control RNAi, n = 44 cells; shootin1b RNAi, n = 48 cells; EGFP, n = 51 cells; EGFP-shootin1-DN, n = 22 cells). (G) Scheme of the glioblastoma invasion assay. Glioblastoma cells were infected with lentiviruses carrying shScramble-mCherry (control RNAi, red) or shShootin1b-EGFP (shootin1b RNAi, green). Tumor spheroids (T) were formed by mixing control RNAi cells (red) and shootin1b RNAi cells (green) in a 1:1 ratio, and co-cultured with the brain cortex organoids (Cx) derived from human iPS cells. The migration distance of the control RNAi cells and shootin1b RNAi cells invaded into the organoids (H) was then analyzed. (H) Fluorescence images of control RNAi cells (magenta) and shootin1b RNAi cells (green) invaded into a brain cortex organoid (Cx) from a tumor spheroid (T). Enlarged images in the rectangles are shown to the right. Scale bar: 50 µm (in the inset, 20 µm). (I) Migration distance of the control RNAi cells and shootin1b RNAi cells invaded into the organoids The number of cells was expressed as the percentage of cells in the migration distance range of 0∼100 μm, 100∼200 μm and 200∼300 μm. Two-tailed unpaired Student’s *t*-test for the ratio at the range of 0∼100 μm. Two-tailed unpaired Welch’s *t*-test for the for the ratio at the range of 100∼200 μm. Two-tailed unpaired Welch’s *t*-test for the for the ratio at the range of 200∼300 μm. n = 5 brain cortex organoids. (J) Weak and strong adhesion-clutch model: adhesion-clutch can mediate both cell motility and immobilization depending on the force amplitude. Weak adhesion-clutch is well-suited for rapid cell migration, without forming strong adhesions that impede cell motility. As transmission of large forces requires strong adhesion, strong adhesion-clutch can immobilize cells against mechanical stress. Data represent means ± SEM; ***, p < 0.01; ns, not significant.

To further analyze a possible role of shootin1b in glioblastoma invasion in tissue, we prepared brain cortex organoids from human induced pluripotent stem (iPS) cells. Equal numbers of the control cells, labeled with mCherry, and shootin1b-knockdown cells (Figure S8C), labeled with EGFP, were combined; the resulting tumor spheroids were co-cultured with the organoid for 72 h to facilitate fusion between them. They were then cultured for further 5 days to allow the glioblastoma cells to invade into the organoids, and their migration distance in the organoid was analyzed (Figure 7G-H). As shown in Figure 7I, shootin1b knockdown significantly inhibited the glioblastoma invasion into the brain cortex organoid. Thus, we conclude that the aberrant activity of the actin-substrate coupling involving shootin1b promotes the glioblastoma cell motility.

## DISCUSSION

### Weak adhesion-clutch for rapid cell migration

The classical adhesion-clutch paradigm alone cannot explain accumulating experimental data on cell migration. As examples, although disruption of the integrin-based adhesion-clutch inhibits migration of melanoma cells and mast cells^52,53^, it does not delay rapid leukocyte migration^10,11,13^ and paradoxically facilitates fibroblast motility^7^. In addition, integrin activation inhibits T cell motility^10^.

This study identified an integrin-independent adhesion-clutch machinery involving shootin1b and L1 that drives rapid dendritic cell migration. Notably, shootin1b and L1 slipped backwards against the adhesive substrate without forming stable adhesions. This slippery actin-substrate coupling transmitted weak backward forces at levels of 10 Pa. The integrin-based adhesion-clutch also undergoes similar retrograde slippage during the initial stages of FA stabilization^5,54^. However, in mesenchymal cells, it generates large traction force that reaches to kPa levels during tension-dependent mechanosensing and FA maturation^8,9,55–58^. As transmission of large forces requires strong adhesion, which impedes cell migration^14,18,59^, the strong forces transmitted by mature FAs would be rather suitable for immobilizing cells against mechanical stress. In fact, fibroblasts do not undergo rapid migration under physiological conditions, and increasing substrate stiffness decreases their random motility^60^ while increasing traction force^9^.

By integrating these previous and present data, we propose that the adhesion-clutch can mediate both cell motility and immobilization depending on the force amplitude; the weak adhesion-clutch is well-suited for rapid cell migration, without forming strong adhesions (Figure 7J). In this regard, the integrin-mediated adhesion-clutch can propel cell migration in a low adhesion context. We consider that the weak adhesion-clutch is mechanistically consistent with the friction-based model^15–19^, and may cooperate with the adhesion-independent mechanisms^11,49,61,62^ to propel efficient cell migration.

### Driving cell migration in response to environmental chemicals

Our data show that the present adhesion-dependent machinery enables cell migration guided by the adhesive ligands in the environment. The ECM protein laminin on the adhesive substrate enhanced the L1-substrate coupling (Figure 2D-E), thereby accelerating dendritic cell migration in an haptokinetic manner (Figure 5A-C). Previous studies have shown that neuronal shootin1a interacts not only with L1 but also with other immunoglobulin-superfamily cell adhesion molecules, cadherin and DCC^29,63^, raising the possibility that the adhesion-clutch system involving shootin1b responds to multiple adhesive ligands in the environment. Furthermore, shootin1b drives chemotaxis through its polarized activation within cells under CCL19 signaling and Pak1-mediated shootin1b phosphorylation (Figure 4F-J). Pak1 is regulated by Rac1 and Cdc42, which are shared key signaling molecules activated by multiple chemoattractants^64–66^. The present adhesion-clutch machinery could be tuned in response to a variety of environmental chemicals, thereby regulating the speed and direction of cell migration and motility.

### Shootin1b promotes glioblastoma cell motility

The present study extends the versatility of the shootin1b- and L1-mediated adhesion-clutch by demonstrating that its aberrant activity enhances glioblastoma cell motility. Glioblastoma is highly invasive, infiltrating and damaging surrounding brain tissue^24,67^. They are rapidly integrated into neural networks^25^, and actively invade along white matter tracts and blood vessels^68,69^. L1 on glioblastoma cells can directly interact with L1 expressed on neurons and oligodendrocytes that form neural networks^70^ and with laminins localized in the basement membrane of blood vessels^71^.

In addition, glioblastoma cells express various neurotransmitter receptors including glutamate receptors^25,72^, and recent studies demonstrated that neuron-glioblastoma interactions through neurotransmitters promotes glioblastoma invasion under receptor activation and calcium signaling^51,73^. Interestingly, glutamate released from the presynapse activates the shootin1a-mediated adhesion-clutch under calcium signaling and shootin1a phosphorylation, thereby promoting the expansion of post-synaptic dendritic spines^29^. Collectively, these data raise the possibility that the adhesion-clutch involving shootin1b and L1 could potentially mediate haptotactic and neuron-driven glioblastoma invasion in the brain.

Despite the availability of clinical treatments, including surgery, radiotherapy and chemotherapy, glioblastoma patients have a poor clinical prognosis^74,75^. Consistent with the previous report that shootin1b is undetectable in the adult mouse brain^30^, we were unable to detect it in normal human and mouse brain-derived astrocytes. Importantly, shootin1b knockdown and shootin1-DN expression decreased the glioblastoma motility, providing a potential target for specific inhibition of glioblastoma invasion in the adult brain.

In conclusion, we present a weak adhesion-clutch suitable for driving physiologically controlled and pathologically enhanced rapid cell migration.

### Limitations of the study

The weak adhesion-clutch paradigm is limited in that it is based on the data obtained from the shootin1b- and L1-mediated adhesion-clutch. Any adhesion-clutch mediated by other molecules could propel cell migration in a low adhesion context. In this regard, shootin1b depletion did not completely inhibit traction forces produced by dendritic cells and glioblastoma cells which were paralleled by their partial inhibition of migration. Thus, future investigation of other molecular machinery and other cell types, along with the measurements of force and migration velocity, will be important for obtaining a more generalized view of the weak adhesion-clutch in cell migration.

This work provides a potential target shootin1b for glioblastoma treatment. However, it is important to note that the cell migration assays were performed in Matrigel and human brain cortex organoids, where the normal neural networks are not established. In addition, glioblastoma is heterogeneous^72^ and we analyzed the effect of shootin1b knockdown using cells derived from a single human patient. Future work to analyze shootin1b-reduced glioblastoma invasion in brain tissue, using cells expressing different levels of shootin1b from multiple patients, will contribute to a more complete understanding of glioblastoma invasion and help guide future therapeutic strategies.

## RESOURCE AVAILABILITY

### Lead Contact

Further information and requests for resources and reagents should be directed to and will be fulfilled by the Lead Contact, Naoyuki Inagaki.

### Materials availability

All unique materials generated in this study are available from the Lead Contact without restriction.

### Data and code availability

Source data of immunoblot and statistical analyses in the figures are available at Mendeley Data (https://data.mendeley.com/datasets/vvhs642dvz/1). Any additional information required to reanalyze the data reported in this paper are available from the lead contact upon request.

## ACKNOWLEDGMENTS

We thank Dr. Peter Friedl (Radboud University Medical Centre), Dr. Michael Sixt (Institute of Science and Technology Austria), Dr. Ewa Paluch (University of Cambridge), Dr. Tim Lämmermann (Max Planck Inst of Immunobiology and Epigenetics) and Dr. Takunori Minegishi (Nara Institute of Science and Technology) for valuable discussions; Mieko Ueda and Kazumi Maekawa for technical support; and Satoko Shimamura for kind encouragement. This research was supported in part by AMED under Grant Number JP17gm0810011 (N.I.), JSPS KAKENHI (JP19H03223, N.I.) JSPS Grants-in-Aid for Early-Career Scientists (JP19K16127, K.B.), and the Osaka Medical Research Foundation for Incurable Diseases (K.B.).

## AUTHOR CONTRIBUTIONS

K.B., D.K., T. K., Y. K., and N.I. designed the experiments. K.B., A. F.-K., M. M., R. T., Z. X, Y. N., M. S., Y. H., H. K.-K., A. K., Y. U., and Y. K. performed the experiments and data analysis. K.B, D.K., Y.K., and N.I wrote the manuscript. N.I. supervised the project. All authors discussed the results and commented on the manuscript.

## STAR METHODS

Detailed methods are provided in the online version of this paper and include the following:

- KEY RESOURCE TABLE
- **EXPERIMENTAL MODEL AND STUDY PARTICIPANT DETAILS**

◦ Ethical considerations
◦ Generation of shootin1 knockout mice
◦ Bone marrow-derived dendritic cell culture and induction of dendritic cell maturation
◦ Transfection of dendritic cells
◦ Glioblastoma cell culture and transfection
◦ HEK293T cell and human astrocyte culture and transfection
◦ Mouse primary astrocyte cell culture
◦ Generation of brain cortex organoid derived from human iPS cells
- METHOD DETAILS

◦ DNA construction and RNAi experiment
◦ RNAi experiment mediated by lentivirus carrying short hairpin RNA
◦ Invasion assay using brain cortex organoid
◦ Flow cytometry analysis
◦ Immunocytochemistry
◦ Immunoprecipitation and Immunoblot
◦ Chemotaxis assay and random migration assay
◦ Glioblastoma cell and astrocyte 3D migration assay
◦ Fluorescent speckle imaging
◦ Traction force microscopy
◦ Analysis of the correlation between the directions of force and cell migration
◦ Analysis of dendritic cell migration in lymph node slice
◦ *In vivo* dendritic cell migration assay
◦ Generation of CCL19 gradients for immunocytochemistry
- QUANTIFICATIONS AND STATISTICAL ANALYSIS

## SUPPLEMENTAL INFORMATION

Supplemental information can be found online at…….

## STAR METHODS

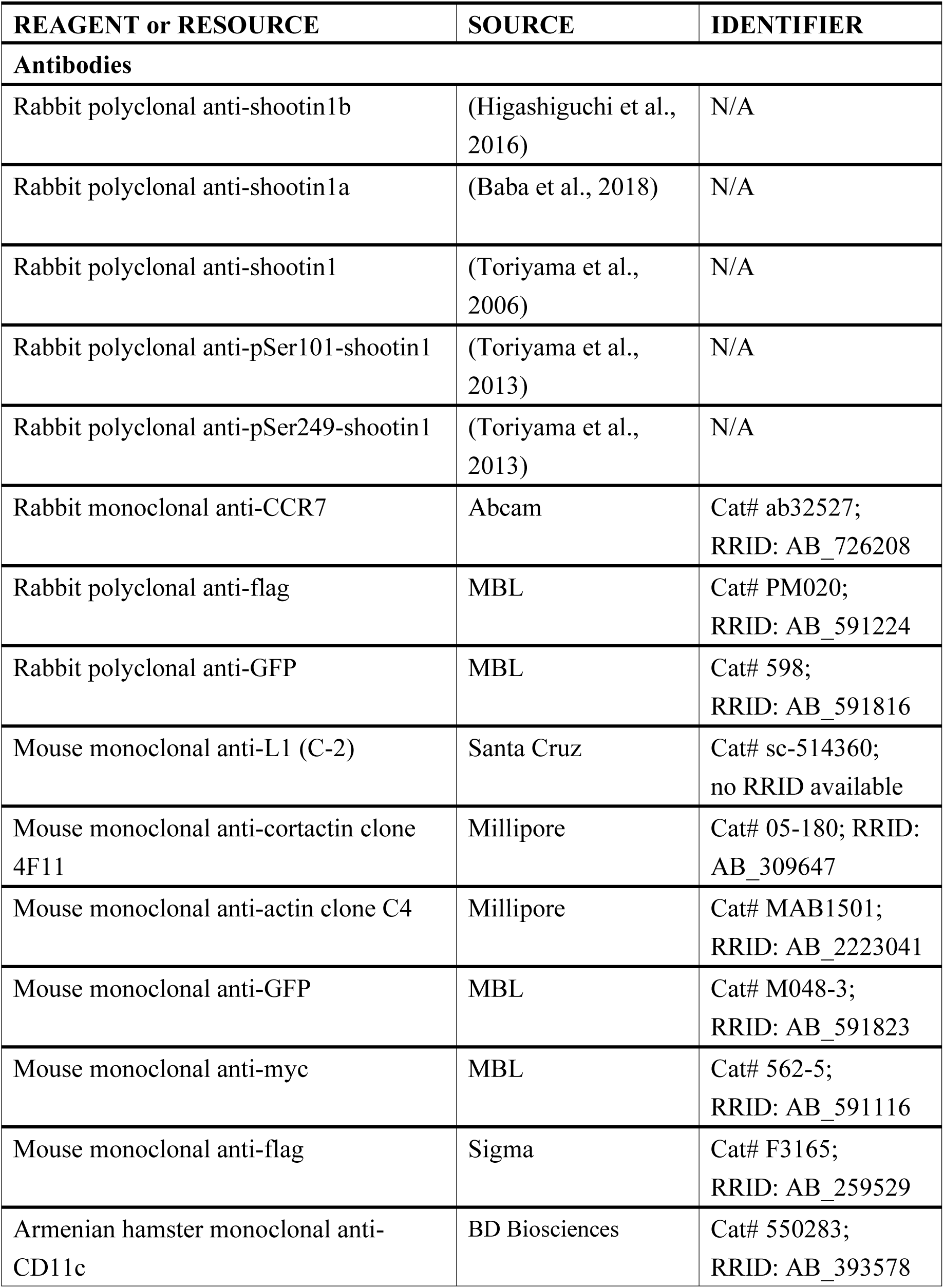
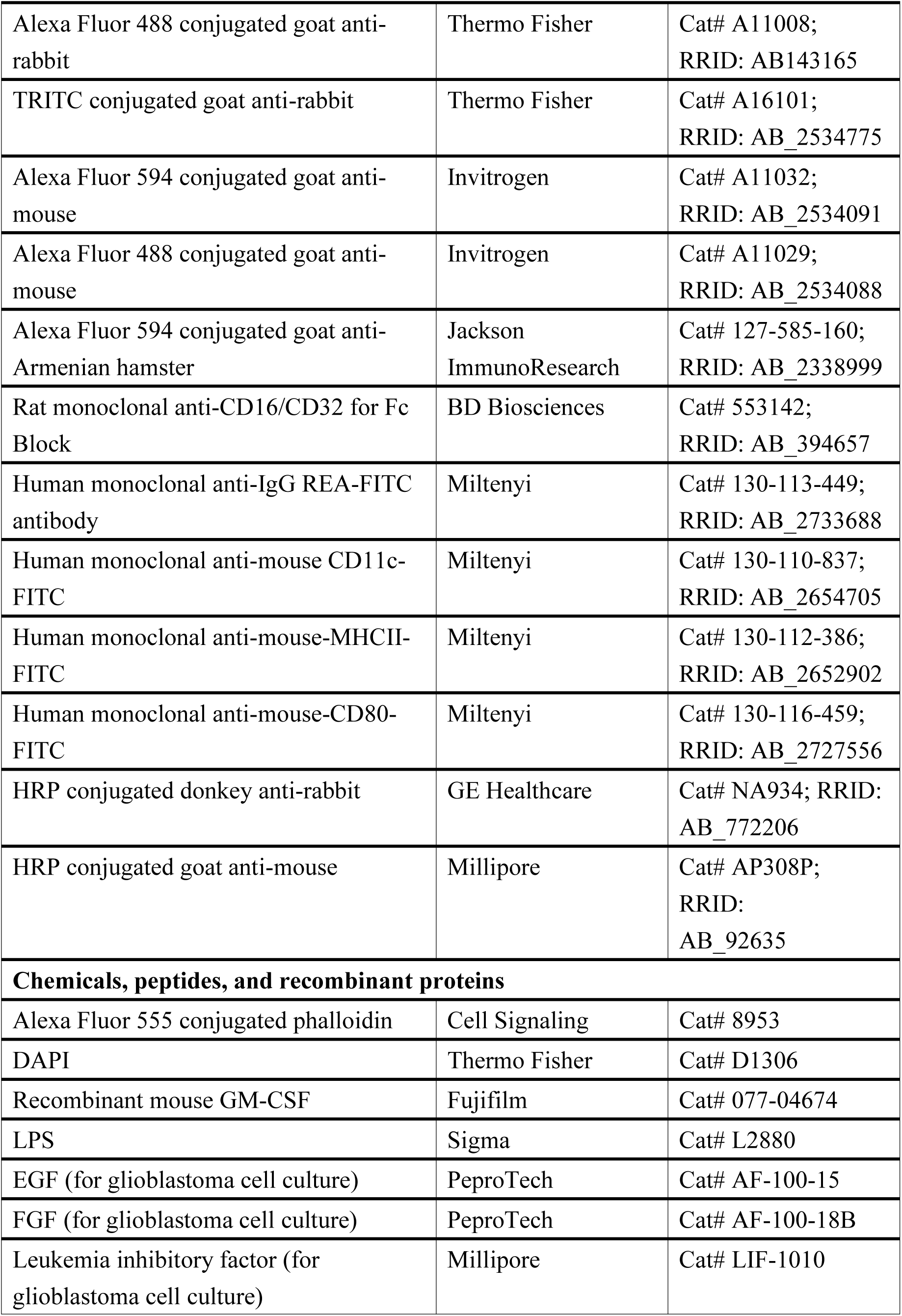
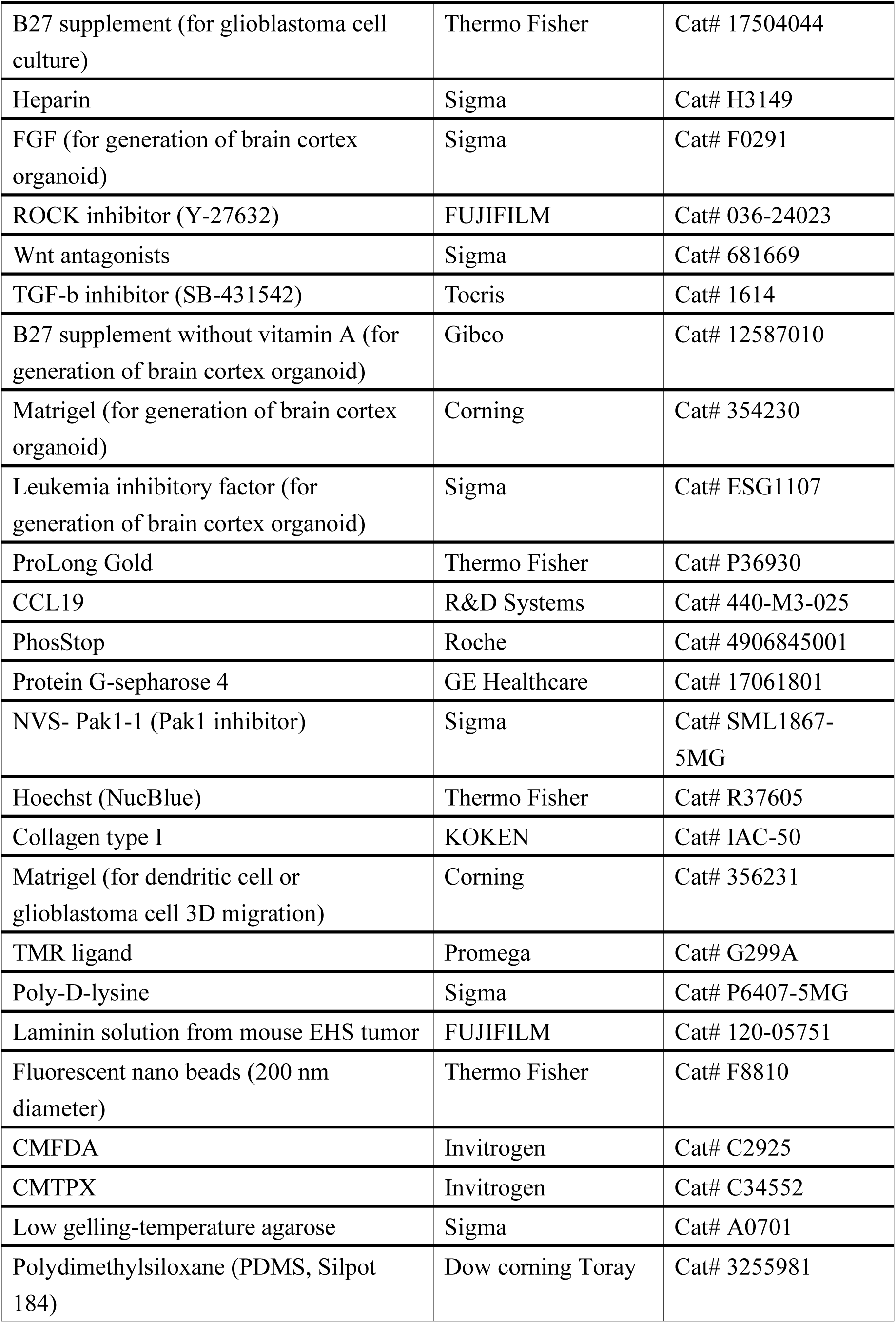
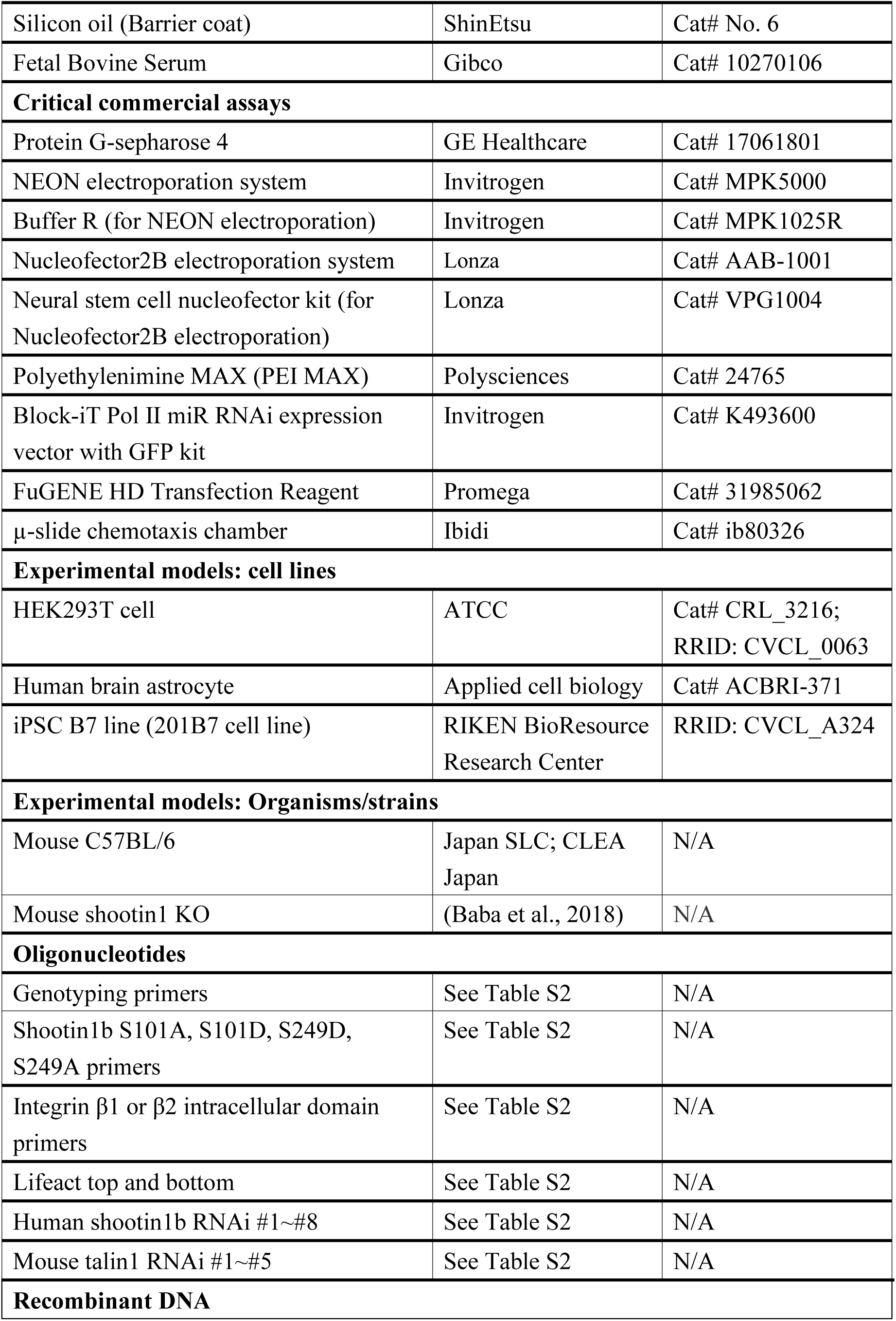
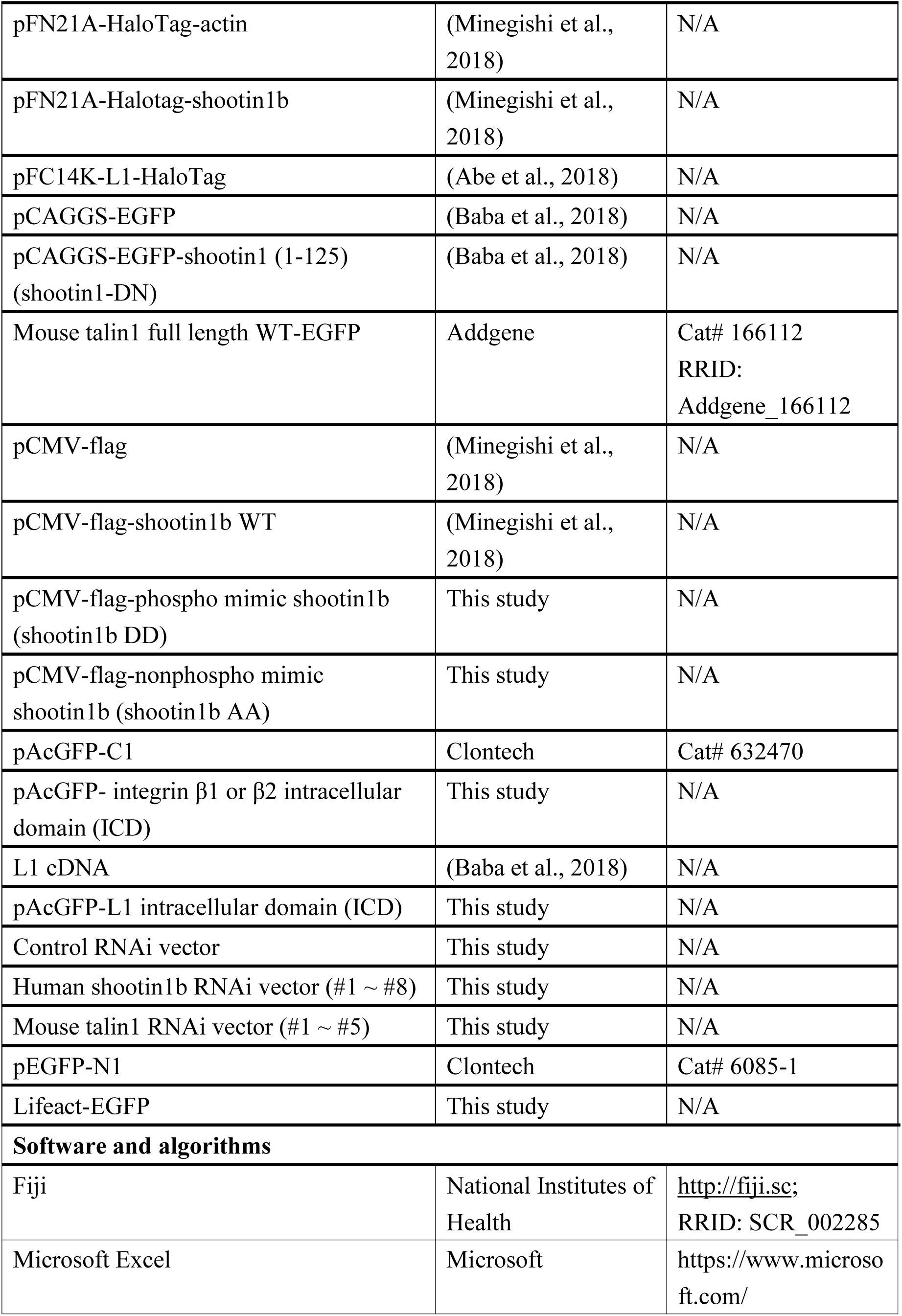
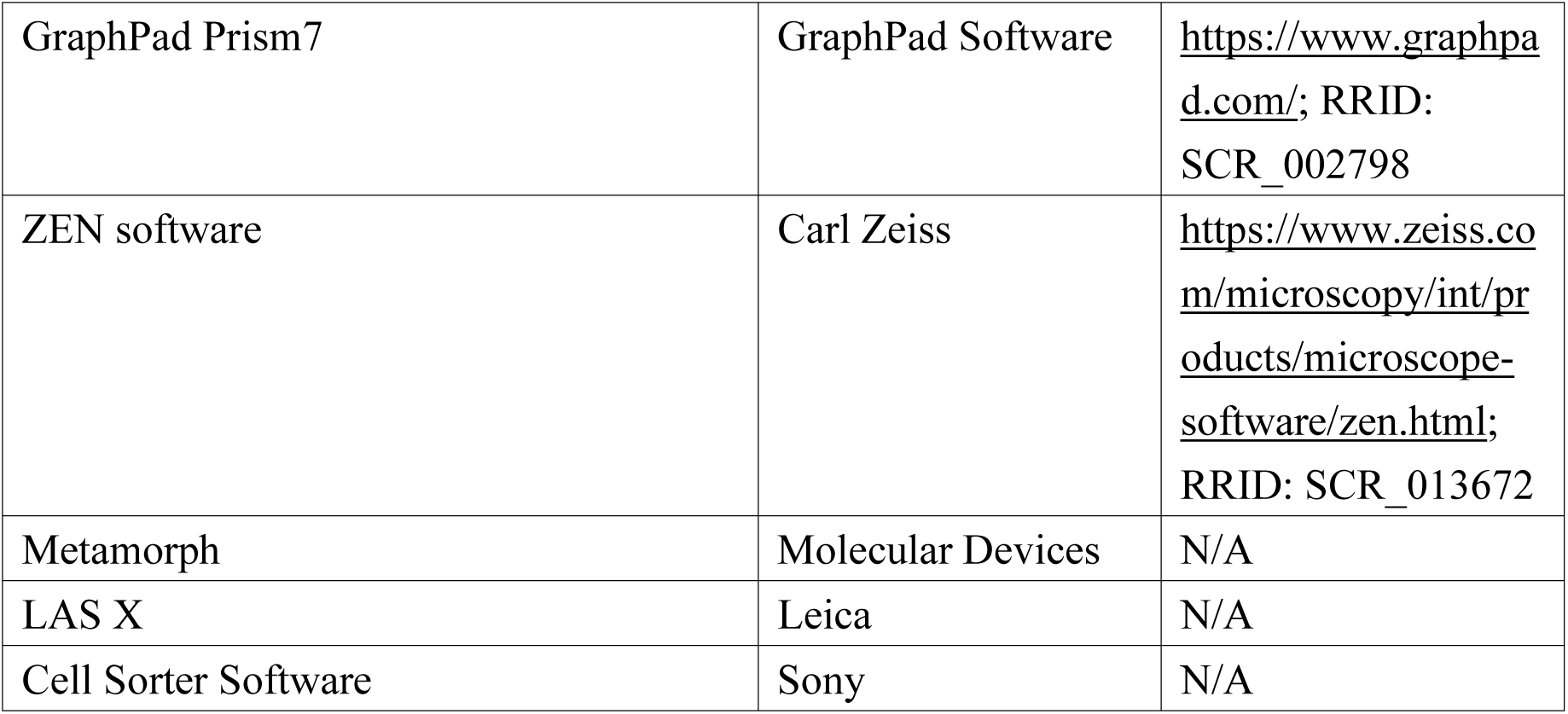
KEY RESOURCES TABLE.

## EXPERIMENTAL MODEL AND STUDY PARTICIPANT DETAILS

### Ethical considerations

This study was carried out in accordance with the principle of the Helsinki Declaration. The use of glioblastoma tissues was approved by the Institutional Review Boards of Nara Institute of Science and Technology (approved number: No. 2018G32), Osaka National Hospital (approved number: No.713) and all collaborating institutions. Surgically removed brain tumor tissues were collected at Osaka National Hospital after obtaining written informed consent. iPSC experiments were conducted with prior approval from NCNP ethical Committee.

### Generation of shootin1 KO mice

All relevant aspects of the experimental procedures were approved by the Institutional Animal Care and Use Committee of Nara Institute of Science and Technology. The generation of shootin1 KO mice is described elsewhere^39^. Chimeric mice were crossed with C57BL/6 mice for at least seven generations before analysis. Male and female shootin1 heterozygous mice were mated to obtain shootin1 KO mice; the offspring genotypes were checked by PCR with the following primers: Genotyping Fw1 (5’-CAGACTGCTACCCACTACCCCCTAC-3’), Genotyping Rv1 (5’-CCTAGAGCTGGACAGCGGATCTGAG-3’), Genotyping Fw2 (5’-CCCAGAAAGCGAAGGAACAAAGCTG-3’), and Genotyping Rv2 (5’-ACCTTGCTCCTTCAAGCTGGTGATG-3’)^29^. Six-to-eight-week-old mice were used for dendritic cell culture^76,77^.

### Bone marrow-derived dendritic cell culture and induction of dendritic cell maturation

Bone marrow-derived dendritic cells were prepared as described^73,74^ with modifications. Tibias and femurs were removed from the surrounding muscle tissue by sterile dissection scissors, and then immersed in 70% ethanol for 5 min. Bones were washed with RPMI-1640 medium (Fujifilm, catalog number: 189-02145) under sterile conditions. Both ends of the bone were cut off with scissors. RPMI-1640 medium was sucked up using the 25-gauge needle (TERUMO, catalog number: NN-2525R) equipped with 1 mL syringe (TERUMO, catalog number: SS-01T). The needle was inserted into the bone cavity to rinse the bone marrow out of the cavity into a sterile culture dish containing RPMI-1640 medium. The suspension of bone marrow-derived cells was filtered by 70 µm strainer (Falcon, catalog number: 352350) and centrifuged at 190 G for 5 min. The supernatant was discarded, and the cell pellet was resuspended in red blood cell lysis buffer (17 mM Tris, 160 mM NH_4_Cl, pH 7.2) for 8 min. Following the centrifugation, the supernatant was discarded and the pellet was washed by PBS. The cell suspension in PBS was centrifuged at 190 G for 5 min. The supernatant was discarded, and the pellet of bone marrow-derived cells was resuspended in the complete RPMI-1640 medium supplemented with 10 % fetal bovine serum (FBS, heat-inactivated, filtered) (Gibco: catalog number: 10270-106), 100 U/mL Penicillin-100 µg/mL streptomycin (Nacalai, catalog number: 26253-84), 2 mM GlutaMAX (Gibco, catalog number: 35050061) and 20 ng/mL recombinant mouse GM-CSF (Fujifilm, catalog number: 077-04674).

Resuspended cells were plated in sterile culture dishes at a density of 1×10^6^ cells/mL and cultured at 37 ℃ in an incubator containing 5 % CO_2_. On day 3, the equal volume of the fresh complete medium containing GM-CSF was added to the cultured medium. On day 7, LPS (Sigma, Escherichia coli o55:B5, catalog number: L2880) was applied to the cultured cells at a final concentration of 200 ng/mL to induce dendritic cell maturation. After 18 ∼ 24 hours of incubation, non-adherent mature dendritic cells were collected by gently pipetting the medium, and then centrifuged at 190 G for 5 min. The supernatant was discarded, and the pellet was resuspended in the fresh complete medium containing GM-CSF and transferred to new culture dishes. Non-adherent mature dendritic cells were collected from culture dishes and used for following experiments. We also confirmed that shootin1b KO does not affect the expression level of the CCL19 receptor, CCR7 (Figure S2E-F), and those of the dendritic cell markers, CD11c, MHC II and CD80, on dendritic cells (Figure S9).

### Transfection of dendritic cells

Dendritic cells were transfected with DNA vectors using the NEON electroporation system (Invitrogen, catalog number: MPK5000). On culture day 7, non-adherent cells were washed by PBS and resuspended in 10 µL resuspension buffer R (Invitrogen: catalog number: MPK1025R) at a density of 2.0×10^7^ cells/mL. The cells were subsequently mixed with DNA vectors and electroporated by the electroporation system. The transfected cells were plated in the complete RPMI-1640 medium. After more than 6h of culture, LPS was applied to the cultured cells at a final concentration of 200 ng/mL to induce dendritic cell maturation.

### Glioblastoma cell culture and transfection

Human glioblastoma-derived cells (KNBTG-8 cells) were isolated from the tumor specimen of a 74-year-old female patient located in the left occipital lobe using the cell sphere method as previously described^78,79^. This tumor was pathologically diagnosed as glioblastoma, IDH-wildtype CNS WHO grade 4, with *TERT* promoter mutation, *EGFR* gene amplification, *PTEN* deletion, and *CDKN2A/B* deletion. Briefly, brain tumor tissue specimens were mechanically dissociated, digested with 0.05% trypsin-EDTA (Thermo Fisher Scientific, Waltham, MA, USA), and then, cultured as floating culture of cell spheres in the completed DMEM/F12 medium (Sigma, catalog number: H4034), Antibiotic-Antimyotic (1:100 dilution, Thermo, catalog number: 15240062), 20 ng/mL EGF (PeproTech, catalog number: AF-100-15), 20 ng/mL FGF (PeproTech, catalog number: AF-100-18B), 10 ng/mL leukemia inhibitory factor (Millipore, LIF-1010), 2% B27 supplement (Thermo, catalog number: 17504044), and 5 ng/mL heparin (Sigma, catalog number: H3149), as described^78,79^. Half of the medium was changed once per week. The spheres were dissociated into single cells every 2 weeks by incubation with Accutase (Nacalai, catalog number: 12679-54) at 37 °C for 10 min. The cells were then resuspended in 50% fresh medium plus 50% conditioned medium at a density of 1 × 10^5^ cells/mL and cultured in culture flasks.

Glioblastoma cells were transfected with DNA vectors using the Nucleofector2B electroporation system (Lonza, catalog number: AAB-1001). After dissociation of the cell sphere by Accutase, the cells were washed with PBS and resuspended in 100 µL nucleofector solution (Lonza, catalog number: VPG1004) at a density of 1.0×10^7^ cells/mL. They were subsequently mixed with DNA vector and electroporated by Nucleofector2B machine.

### HEK293T cell and human astrocyte culture and transfection

HEK293T cells (ATCC, catalog number: CRL3216) and primary human brain astrocytes (Applied cell biology, catalog number: ACBRI 371)^80^ were cultured in DMEM (Sigma, catalog number: D6429) supplemented with 10% FBS, 100 U/mL Penicillin-100 µg/mL streptomycin, 0.01 % Gentamycin (Gibco, catalog number: 15750060) as described previously^39^. HEK 293T cells were transfected with plasmid DNA using PEI MAX (Polysciences, catalog number: 24765) following the manufacture’s protocol.

### Mouse primary astrocyte cell culture

Primary cultured astrocytes were prepared from the cerebral cortices postnatal day 1 mice as described^81^. They were cultured in Dulbeco’s modified Eagle’s medium supplemented with 10% FBS, 100 U/mL Penicillin-100 µg/mL streptomycin, and 0.01 % Gentamycin. After two weeks, the cells were grown to a confluent monolayer and used for the immunoblot analysis.

### Generation of brain cortex organoid derived from human iPS cells

iPSC line B7 (201B7 cell line) were obtained from RIKEN BioResource Research Center and cultured on 0.1% gelain-coated plates with irradiated mouse embryonic feeders (MEF) in Primate ESC Medium (Repro, catalog number: RCHEMD001) supplemented with 10 ng/mL of human basic Fibroblast Growth Factor (FGF, Sigma, catalog number: F0291). They were maintained at 5% CO_2_ at 37℃ with daily medium change and were passaged every 6 days at 1:3–1:6 using cell dissociation (CTK) solution (Repro, catalog number: RCHETO002)^82^.

Cortical organoid differentiation was performed as described^83,84^ with several modifications. Briefly, iPSCs were dissociated to single cells and plated into low-attachment V-bottom 96-well plates (Greiner Bio-One, catalog number: 651161) to form aggregates in Glasgow’s Minimal Essential Medium (GMEM, Gibco, catalog number: 11710035) supplemented with 20% KSR (knockout serum replacement, Gibco, catalog number: 10828010), 1×MEM-NEAA solution (modified Eagle’s medium (MEM) non-essential amino acids solution (Gibco, catalog number: 11140050), 100 µg/mL of Primocin (InvivoGen, catalog number: 14860-94), 0.1 mM 2-Mercaptoethanol (Gibco, catalog number: 21985023), 1 mM Sodium Pyruvate (Gibco, catalog number: 11360070), 20 µM ROCK inhibitor (FUJIFILM, catalog number: 036-24023), 3 µM Wnt antagonists (Sigma, catalog number: 681669) and 5 µM TGF-b inhibitor (Tocris, catalog number: 1614). Half of the medium was changed every 2–3 days. ROCK inhibitor was removed after 6 days. Aggregates were then transferred to a hyperoxygenated incubator at 5% CO_2_ and 40% O_2_ and maintained in DMEM/F12 supplemented with N2 supplement (Gibco catalog number: 17502048), GlutaMAX (Gibco, catalog number: 35050061), chemically defined lipid concentrate (CDLC, Gibco, catalog number: 11905031), and 0.4% methylcellulose (Sigma, catalog number: M7140). The culture medium was changed every 2–3 days thereafter.

On day 35, the organoids were cut in half using Micro Scissors, and the medium was changed to N2B27 medium containing DMEM/F12 supplemented with N2 supplement, GlutaMAX, CDLC, 0.4% methylcellulose, B27 without vitamin A (Gibco, catalog number: 12587010), 1% growth factor-reduced Matrigel (Corning, catalog number: 354230), and 5 µg/mL of heparin (Sigma, catalog number:H3149). On day 56, organoids were cut in half and transferred to oxygen permeable dishes containing N2B27 medium. Organoids were subsequently cut in half every 2 weeks and routinely sustained for up to 120 days. For STAT3 activation to differentiate and obtain mature cortical organoids, recombinant mouse LIF (Sigma, catalog number: ESG1107) was added at 2,000 U/mL from day 35 onward.

## METHOD DETAILS

### DNA construction and RNAi experiment

Preparations of vectors to express pCMV-flag-shootin1b, pFN21A-Halotag-shootin1b, pFN21A-Halotag-actin, pFC14K-L1-HaloTag, pCAGG-myc-shootin1(1-125), pCAGGS-EGFP-shootin1(1-125) have been described have been described previously^39,85^. pEGFP_WT-talin1(1-2541) (mouse talin1 full length WT-EGFP) was a gift from Vesa Hytönen (Addgene plasmid # 166112 ; http://n2t.net/addgene:166112 ; RRID: Addgene 166112)^86^.

To generate pEGFP-N1-LifeAct vector, the annealed LifeAct (forward: 5’-TCGAGATGGGTGTCGCAGATTTGATCAAGAAATTCGAAAGCATCTCAAAGGAAGAAGGG-3’; reverse: 5’-GATCCCTTCTTCCTTTGAGATGCTTTCGAATTTCTTGATCAAATCTGCGACAC CCATC-3’) was fused to N-terminal of EGFP-N1 vectors (Clontech, catalog number: 6085-1).

Unphosphorylated (S101A/S249A: AA) and phosphomimic (S101D/S249D: DD) mutants of rat shootin1b^30^ were generated with the QuickChange Ⅱ site -directed mutagenesis kit (Stratagene) to replace each serine (S) with alanine (A) or aspartate (D) using the following primers : S101A Fw (5’-AAAAGAATCGCCATGCTATACATG-3’), S101A Rv (5’-CATGTATAGCATGGCGATTCTTTT-3’), S101D Fw (5’-AAAAGAATCGACATGCTATACATG-3’), S101D Rv (5’-CATGTATAGCATGTCGATTCTTTT-3’), S249A Fw (5’-AAGAGACAAGCCCACCTTCTGCTG-3’), S249A Rv (5’-CAGCAGAAGGTGGGCTTGTCTCTT-3’), S249D Fw (5’-AAGAGACAAGACCACCTTCTGCTG-3’), S249D Rv (5’-CAGCAGAAGGTGGTCTTGTCTCTT-3’). cDNA of shootin1b DD or AA mutants was sub-cloned into pCMV-flag vector^85^.

To generate AcGFP-integrin β1 and β2 intracellular domains (ICDs), integrin β1- or β2-ICD was amplified mouse brain cDNA (for integrin β1 cDNA) or mouse dendritic cell cDNA (for integrin β2 cDNA) by PCR using the following primers : Integrin β1-ICD Fw (5’-AAAGGATCCAAACTTTTAATGATAATTCAT-3’), Integrin β1-ICD Rv (5’-AAAGTCGACTCATTTTCCCTCATACTTCG-3’), Integrin β2-ICD Fw (5’-AAAGGATCCAAGGCCCTGACCCACCT-3’), Integrin β2-ICD Rv (5’-AAAGTCGACCTAGCTTTCAGCAAACTTGGG-3’). cDNA of integrin β1-, β2-ICD and L1-ICD^39^ were subcloned into AcGFP vector (Clontech, catalog number: 632470). For mouse talin1 and human shootin1b RNAi experiments, we used a Block-iT Pol Ⅱ miR RNAi expression vector kit (Invitrogen, catalog number: K493600) and Block-iT RNAi designer. The targeting sequences for mouse talin1 are following: mouse talin1 #1 (5’-CATGATGAGTATTCACTGGTT-3’), #2 (5’-TTAAATCAGGCCGCCACAGAA-3’), #3 (5’-TCACATCAAACACCGAGTACA-3’), #4 (5’-TCCTGGTAGCTTGCAAGGTCA-3’) or #5 (5’-GGGCCTCAGATAACCTGGTAA-3’). The targeting sequences for human shootin1b are following: human shootin1b #1 (5’-AAGCAATAGGCGAATATGAAG-3’), #2 (5’-AAGACTTGTCGAGAAAGTGCT-3’), #3 (5’-AAGCTGGGACCAGATGTAATA-3’),#4 (5’-GTCCATGTTAGCTGTAGAAGA-3’), #5 (5’-GCTGGAGAAGGACCTTCGAAA-3’) or #6 (5’-AAGTTCCAAGGTTACGTTTCA-3’). The oligonucleotides containing target sequences for mouse talin1 (#1 ∼ #5) and human shootin1b (#1∼#6) were inserted into the RNAi vector to express microRNAs against the targeting sequences. To confirm the reduction of mouse talin1 and human shootin1b expression, immunoblot was performed using lysates of HEK293T cells transfected with mouse talin1-EGFP (RRID: Addgene 166112) and each of mouse talin1 RNAi vector (#1 ∼ #5) (Figure S6B) or lysates of HEK293T cells transfected with each of human shootin1b RNAi vector (#1 ∼ #6) (Figure S8B).

### RNAi experiment mediated by infection of lentivirus carrying short hairpin RNA

The lentiviral vectors carrying short hairpin RNA (shRNA) against human shootin1b with EGFP and mCherry (pLKO.1-EGFP and pLKO.1-mCherry) were generated by replacing the gene encoding PuroR in pLKO.1 puro (Addgene Plasmid #8453) with EGFP and mCherry, respectively. The inserted oligonucleotides containing target sequences were designed according to the online cloning protocol of pLKO.1 (https://www.addgene.org/protocols/plko/). The target sequences used in this study are as follows: (5’-CCTAAGGTTAAGTCGCCCTCG-3’) for a scramble RNA as a negative control, #7 (5’-TGGTCATAGAGGAAGTTAATT-3’) and #8 (5’-AAGACTTGTCGAGAAAGTGCT-3’) for human shootin1b. The oligos were inserted into pLKO.1-mCherry and pLKO.1-EGFP vectors, yielding to pLKO.1-shScramble-mCherry and pLKO.1-shShootin1b-EGFP to produce lentivirus carrying shRNAs against Scramble (control) and human shootin1b (#7) were used for RNAi experiments in Figure 7G-H.

For lentiviral production, 6 x 10^6^ HEK293FT cells were plated on a 10-cm culture dish for one day prior to transfection in complete DMEM medium (Sigma catalog number: D5796) supplemented with 10% FBS, (catalog number: F7524), 292 μg/mL of L-Glutamine, 100 U/mL of penicillin, and 100 μg/mL of streptomycin (Gibco 10378016). 10 μg of pLKO.1 lentiviral vector, 3 μg of psPAX2 (Addgene Plasmid #12260), 1.5 μg of pMD2-VSVG (Addgene Plasmid #12259) and 36 μl of FuGENE HD Transfection Reagent (Promega, catalog number: E2312) were mixed in 350 μl of Opti-MEM (Gibco, catalog number: 31985062) and stored at room temperature for > 15 min before the transfection, and then this mixture was added to the medium of HEK293FT cells. After 16 h incubation, the culture medium was replaced with 10 ml of a fresh complete DMEM medium. After 6 hours, the culture medium was further replaced with 4 ml of the optimal culture medium for glioblastoma cells. The 4 mL of viral supernatant was collected every 24 hours followed by the addition of 4 mL of the fresh optimal glioblastoma cell culture medium. The collected supernatant (total 8 mL) was filtered by 0.45 μm filter (Millipore, catalog number: SLHPR33RS).

For lentivirus infection to glioblastoma cells, the viral supernatant was added to the culture medium. Twenty four hours after the infection, the virus-containing medium was replaced with a fresh medium. Expression of mCherry and EGFP (pLKO.1-shScramble-mCherry and pLKO.1-shShootin1b-EGFP) was confirmed within 48 h after infection under an epifluorescent microscopy. Successful reduction of shootin1b expression level was validated by immunoblotting using the lysates of the infected glioblastoma cells (shootin1b RNAi #7, #8) (Figure S8C).

### Invasion assay using brain cortex organoid

Tumor spheroids were formed by mixing glioblastoma cells expressing shScramble-mCherry or shShootin1b-EGFP in a 1:1 ratio (1.5 × 10^4^ cells), followed by 72 h incubation in a V-bottom 96-well plate (Greiner, catalog number: 651101) at 37 ℃, 5% CO_2_. Then, the tumor spheroids were co-cultured with the brain cortex organoid for an additional 72 h at 37 ℃, 5% CO_2_ to allow fusion between the spheroids and organoids in N2B27 organoid culture medium. mCherry- or EGFP-labelled glioblastoma cells invaded from the spheroids to the organoids. Five days after fusion between the spheroids and organoids, they were fixed by 4% Paraformaldehyde in PBS for 40 minutes on ice, and then washed twice in PBS. The fixed organoids were then cryo-preserved in 30% sucrose in PBS overnight at 4 °C until completely submersed and embedded in OCT-compound blocks. Subsequently, the organoids were cryo-sectioned at 10μm thickness, and collected onto Superfrost Plus slides (Fisher Scientific, catalog number: 1255015). Nuclei were stained with DAPI for 5 min before mounting slides with Prolong Antifade Mountant (Invitrogen, catalog number: P36970).

Fluorescence images were obtained by an Olympus FV3000 microscope equipped with ×20 and ×40 dry objectives. All images were compiled in ImageJ and Adobe Photoshop, with image adjustments applied to the entire image and restricted to brightness, contrast, and levels. Images shown in figures were obtained and processed in parallel using identical imaging settings. To quantify the distribution of infected tumor cells, the invasion distance from the spheroid to the organoid was first measured in the slice of tumor-spheroid-fused organoid. Then, the ratio of the number of shScramble cells or shShootin1b cells invaded in the organoid at each range of 0∼100 μm, 100∼200 μm, or 200∼300 μm to the number of total cells (shScramble cells or shShootin1b cells) was calculated.

### Flow cytometry analysis

Non-adherent mature dendritic cells were collected and washed with PBS. The cells were then resuspended in HBSS without Ca^2+^ buffer (Gibco, catalog number: 14175-095) containing 2 mM EDTA and 1% BSA at a density of 1×10^6^ cells/mL. For Fc receptor blocking, anti-mouse CD16/CD32 antibody (BD Biosciences, catalog number: 553142) was added to the cell suspension at 1:500 dilution at 4℃ to prevent non-specific antibody binding. After 30 min, the cells were washed with PBS and resuspended in the buffer at 4℃. To analyze the cell surface markers of mature dendritic, the cells were stained for 30 min at 4℃ by the following antibodies; anti-mouse CD11c-FITC (1:500 dilution) (Miltenyi, catalog number: 130-110-837), anti-mouse-MHCII-FITC (1:500 dilution) (Miltenyi, catalog number: 130-112-386), and anti-mouse-CD80-FITC (1:500 dilution) (Miltenyi, catalog number: 130-116-459) antibodies. The anti-IgG REA-FITC antibody (1:500 dilution) (Miltenyi, catalog number: 130-113-449) was used as an isotype IgG control antibody for negative staining. The stained cells were washed with PBS and resuspended in the HBSS without Ca^2+^ buffer at a density of 1×10^6^ cells/mL. The cell suspension was filtrated by 35 µm cell strainer cap with a 5 mL tube (Falcon, catalog number: 352235) to remove the aggregated cells, and then loaded into the MA900 flow cytometer (Sony, catalog number: P1FFW1000040-1) for the analysis of the dendritic cell surface markers. The percentage of dendritic cell surface marker positive cells were calculated using Cell Sorter Software (Sony). The range of negative cells was determined using the isotype IgG control antibody (see magenta frame in Figure S9A). More than 90 % of the cells were positive for the mature dendritic cell markers in WT and shootin1b KO cells (see green frames in Figure S9B-D).

### Immunocytochemistry

Dendritic cells and glioblastoma cells were fixed with 3.7 % formaldehyde in PBS for 10 min at room temperature and for 5 min on ice, followed by treatment with 0.05% Triton X-100 in PBS for 15 min on ice and 10 % FBS in PBS for 1 hour at room temperature. They were then incubated overnight at 4°C with primary antibodies diluted in PBS containing 10% FBS. The following primary antibodies were used: rabbit anti-shootin1b (1:2000 dilution), rabbit anti-shootin1a (1:2000 dilution), rabbit anti-pSer249 (1:1000 dilution), mouse anti-cortactin (1:500 dilution) (Millipore, catalog number: 05-180), mouse anti-L1 (1:500 dilution) (Santa Cruz, catalog number: sc-514360), mouse anti-myc (1:1000) (MBL, catalog number: 562-5), and Armenian hamster anti-CD11c (1:400 dilution) (BD Biosciences, catalog number: 550283) antibodies. The cells were washed with PBS, and then incubated with secondary antibodies diluted in PBS for 1 hour at room temperature. The following secondary antibodies were used: Alexa Fluor 488 conjugated goat anti-rabbit (1:1000 dilution) (Thermo Fisher, catalog number: A11008), TRITC conjugated goat anti-rabbit (1:1000 dilution) (Thermo Fisher, catalog number: A16101), Alexa Fluor 594 conjugated goat anti-mouse (1:1000 dilution) (Invitrogen, catalog number: A11032), Alexa Fluor 488 conjugated goat anti-mouse (1:1000 dilution) (Invitrogen, catalog number: A11029) and Alexa Fluor 594 conjugated goat anti-Armenian hamster (1:1000 dilution) (Jackson ImmunoResearch, catalog number: 127-585-160) antibodies. For F-actin staining, cells were stained with Alexa Fluor 555 conjugated phalloidin (1:100 dilution) (Cell Signaling, catalog number: 8953) for 30 min at room temperature. Immunostained cells were mounted with 50% (v/v) glycerol in PBS for a fluorescence microscope or ProLong Gold (Thermo Fisher, catalog number: P36930) for a Leica Stellaris 8 STED microscope.

Fluorescence images were acquired using a fluorescence microscope (BZ-X710, KEYENCE) equipped with a UPlansApo 60 × oil, 1.35 NA objective (Olympus), a Plan-Apochromat 100 × oil, 1.45 NA objective (KEYENCE) and imaging software (BZ-X Analyzer software) (BZH4A, KEYENCE) or a Leica Stellaris 8 STED microscope equipped with an HC PL APO 93×/1.30-motCORR glycerol immersion objective lens and imaging software (LAS X) (Leica Microsystems). ProLong Gold was used as mounting agent for STED image acquisition. Dendritic cell images were acquired with x/y pixel size of 41 or 68 nm for the whole cell (left panels in Figures 1F and S2C-D) and 15 or 24 nm for the leading edge (right panels in Figures 1F and S2C-D). Glioblastoma cell images (Figure 6B) were acquired with x/y pixel size of 41 nm for the whole cell and 12 nm for the tip of the tumor microtube. Stimulated emission depletion was accomplished with a 660 nm STED laser. Excitation was provided by a white light laser at the desired wavelength for each sample.

### Immunoprecipitation and Immunoblot

Immunoprecipitation and immunoblot were performed as described^26,39^. For immunoprecipitation with dendritic cells after stimulation with CCL19 (0, 20, 200 ng/mL) (R&D Systems, catalog number: 440-M3-025) for 30 min (Figure 3C) and HEK293T cells expressing AcGFP-tagged proteins and flag-shootin1b (Figure S6A), cell lysates were collected by NP40-Triton lysis buffer (0.5% NP-40, 0.5% Triton X-100, 20 mM HEPES pH 7.5, 3 mM MgCl2, 100 mM NaCl, 1 mM EGTA, 1 mM DTT, 1 mM PMSF, 0.01 mM leupeptin, 1× PhosStop). The supernatants of the lysates were incubated with anti-shootin1 antibody (Figure 3C) or anti-flag antibody (Sigma, catalog number: F3165) (Figure S6A) overnight at 4℃, and then the immunocomplexes were precipitated with protein G-sepharose 4 (GE Healthcare, catalog number: 17061801). After washing the beads with wash buffer (0.1% Tween 20, 20 mM HEPES pH 7.5, 3 mM MgCl2, 100 mM NaCl, 1 mM EGTA, 1 mM DTT), the immunocomplexes were detected by immunoblot using rabbit anti-shootin1b (1:5000 dilution), mouse anti-cortactin (1:1000 dilution), mouse anti-L1 (1:1000 dilution), rabbit anti-GFP (1:1000 dilution) (MBL, catalog number: 598) and rabbit anti-flag antibodies (1:1000 dilution) (MBL, catalog number: PM020).

To determine shootin1b phosphorylation, dendritic cells were incubated with 200 ng/mL CCL19 (R&D Systems, catalog number: 440-M3-025) at each time point (0, 10, 30, 60 min) (Figure S4A) or each CCL19 concentration (0, 20, 200, 2000 ng/mL) for 30 min (Figure 3A). To analyze Pak1-mediated shootin1b phosphorylation (Figure S4C), mature dendritic cells were incubated with 0.1 % BSA (control), 200 ng/mL CCL19 or 200 ng/mL CCL19 + 250 nM NVS-Pak1-1 (Pak1 inhibitor) (Sigma, catalog number: SML1867-5MG). Dendritic cell lysates were collected using RIPA buffer (50 mM Tris-HCl pH8.0, 150 mM NaCl, 1 mM EDTA, 1% Triton, 0.1% SDS, 0.1% sodium deoxycholate, 1 mM DTT, 1 mM PMSF, 0.01 mM leupeptin, 1× PhosStop). Immunoblot was performed using rabbit anti-pSer101-shootin1 (1:1000 dilution), rabbit anti-pSer249-shootin1 (1:2000 dilution) and rabbit shootin1b (1:5000 dilution) antibodies as described^39^. For the mouse talin1 and human shootin1b RNAi experiments (Figures S6B, S8B), HEK293T cells were transfected with talin1-EGFP + talin1 RNAi vector or shootin1b RNAi vector using PEI MAX. After 48 hours, lysates of the transfected cell were collected by RIPA buffer. Immunoblot was performed using mouse anti-GFP (1:1000 dilution) (MBL, catalog number: M048-3), mouse anti-actin (1:5000 dilution) (Millipore, catalog number: MAB1501) and rabbit anti-shootin1b (1:5000 dilution) antibodies. To analyze the expression of shootin1b, shootin1a, cortactin, L1 and CCR7 in mature dendritic cells, glioblastoma cells and human astrocytes, the cell lysates were collected by RIPA buffer. Immunoblot was performed using rabbit anti-shootin1b (1:5000), rabbit anti-shootin1a (1:5000), mouse anti-cortactin (1:1000 dilution), mouse anti-L1 (1:1000 dilution) and rabbit anti-CCR7 (1:4000) antibodies.

### Chemotaxis assay and random migration assay

To accurately track migrating dendritic cells, the nuclei of cells were stained with Hoechst (1:2000 dilution) (Thermo Fisher, catalog number R37605) for 30 min in the serum-free complete RPMI-1640 medium. For the chemotaxis assay, dendritic cells (1×10^5^ cells) were embedded in the 100 µL mixture of 1.5 mg/mL collagen gel (bovine collagen type I) (KOKEN, catalog number: IAC-50) + 10 % Matrigel (Corning, catalog number: 356231) mixed to give the compositions listed in Table S1. Matrigel was used to allow laminin-mediated migration in collagen gel^47^. The cell mixture was placed in a µ-slide chemotaxis chamber (Ibidi, catalog number: ib80326) according to the manufacturer’s instructions and incubated at 37 ℃, 5 % CO_2_ for 30 min to allow polymerization of collagen fibers and gelation of Matrigel. 600 ng/mL CCL19^11^ (R&D Systems, catalog number: 440-M3-025) in 10% FBS/RPMI-1640 was added to the CCL19 source side of the reservoir, while 10 % FBS/RPMI-1640 medium (chemoattractant-free medium) was added to the opposite side.

For the random migration assay under the bath application of CCL19, dendritic cells (5×10^4^ cells) were embedded in the 100 µL mixture of 1.5 mg/mL collagen gel + 10 % Matrigel + CCL19 (20 or 200 ng/mL) mixed to give the compositions listed in Table S1. The cell-mixture was placed on the glass bottom dish (Matsunami, catalog number: D11130H). After 30 min incubation to allow polymerization of collagen fibers and gelation of Matrigel, the surface of the mixture was covered with the complete RPMI1640 medium including CCL19 (20 or 200 ng/mL). Time-lapse images were acquired every 1 min for 180 min at 37 ℃ using a fluorescence microscope (IX81, Olympus) equipped with an EM-CCD (Ixon DU888, Andor), a Plan Fluor × 20, 0.45 NA objective (Olympus) and MetaMorph software. Migrating cells were tracked using the manual tracking (Fiji) to calculate migration speed and the chemotaxis index (straight distance toward the CCL19 source/total distance) (Figure 4C).

### Glioblastoma cell and astrocyte 3D migration assay

Glioblastoma cells or human astrocytes were embedded in 50 % Matrigel diluted with the complete DMEM/F12 medium at a density of 5×10^5^ cells/mL. The Matrigel-cell mixture was placed on the glass bottom dish (Matsunami, catalog number: D11130H), and then this mixture was incubated for 30 min at 37 ℃, 5 % CO_2_ to allow gelation of Matrigel. The surface of the mixture was covered with complete L15 medium (Gibco, catalog number: 21083027) supplemented with Antibiotic-Antimyotic (1:100 dilution), 20 ng/mL EGF, 20 ng/mL FGF, 10 ng/mL leukemia inhibitory factor, 2% B27 supplement and 5 ng/mL heparin + conditioned medium in a 1:1 ratio. After 24h of culture, time-lapse images were acquired every 10 min for 480 min at 37 ℃ using a fluorescence microscope (IX81, Olympus) equipped with an EM-CCD (Ixon DU888, Andor), a Plan Fluor × 20, 0.45 NA objective (Olympus), and MetaMorph software.

### Fluorescent speckle imaging

Fluorescent speckle imaging was performed as previously described^31^ with modifications. Dendritic cells or glioblastoma cells were treated with TMR ligand (Promega, catalog number: G299A) at 1:2000 (final concentration 50 nM) in the serum-free complete RPMI-1640 medium (for dendritic cells) or the serum-free complete DMEM/F12 medium (for glioblastoma cells) for 30 min at 37 ℃, 5 % CO_2_ to visualize HaloTag-actin, HaloTag-shootin1b or Halotag-L1. The ligand was then washed with PBS, and cells were incubated in the complete RPMI-1640 medium or the complete DMEM/F12 medium at 37 ℃, 5 % CO_2_.

For fluorescent speckle imaging in dendritic cells, a 1 % agarose block was prepared by mixing the following in a 1:2:1 ratio : (i) 56 ℃ pre-warmed 2×HBSS (Sigma, catalog number: H-1387), (ii) 56 ℃ pre-warmed RPMI-1640 medium supplemented with 20 % FBS, (iii) 4 % agarose (Sigma, catalog number: 16500) in warmed water. Recombinant mouse CCL19 (20 ng/mL or 200 ng/mL) (R&D Systems, catalog number: 440-M3-025) was added to liquid agarose when liquid agarose was heated to 37 ℃. The liquid agarose was then poured into a glass bottom dish coated with PDL (Sigma, catalog number: P6407) and laminin (Fujifilm, catalog number: 120-05751) and allowed to solidify for 1 hour at room temperature. After the TMR treatment and PBS washing, dendritic cells were resuspended in the complete RPMI1640 medium containing CCL19 (20 or 200 ng/mL) at a density of 1×10^6^ cells/mL. The cell suspension of the cells was injected under the agarose block on the laminin-coated glass bottom dish. For fluorescent speckle imaging in glioblastoma cells, 1×10^5^ glioblastoma cells were placed on the glass bottom dish coated subsequently by PDL and laminin. After the TMR treatment and PBS washing, the medium was changed to 50% conditioned medium + 50% the complete L15 medium.

Fluorescent speckles were observed at the leading edge of dendritic cells and at the tip of the tumor microtube of glioblastoma cells at 37 ℃ using a TIRF microscope (IX81; Olympus) equipped with an EM-CCD (Ixon3, Andor), a complementary metal oxide semiconductor (CMOS) camera (ORCA Flash 4.0LT, HAMAMATSU), a UAPON ×100 1.49 NA (Olympus) and MetaMorph software^87^ for dendritic cells, or using a fluorescence microscope (AxioObserver Z1, Carl Zeiss) equipped with a complementary metal oxide semiconductor (CMOS, ORCA Flash4.0 V2, Hamamatsu), a Plan-Apochromat 100x, 1.40 NA (Carl Zeiss), and imaging software (ZEN2012, Carl Zeiss) for glioblastoma cells^31^. Fluorescence time-lapse images were acquired every 2 sec for dendritic cells and 5 sec for glioblastoma cells. We analyzed the fluorescent speckles that could be traced for at least 5 frames during imaging. F-actin flow velocity was calculated by tracing the speckle of HaloTag-actin. Actin polymerization rate was calculated as the sum of the F-actin retrograde flow and extension rates as reported^31^.

### Traction force microscopy

Traction force microscopy was performed as previously described^31^ with modifications. Polyacrylamide gels with embedded fluorescent nano beads (200 nm diameter) (Thermo, catalog number: F8810) were prepared as described^31^. To measure forces under dendritic cells, they were stained for 30 min by 1 µM CMFDA (Invitrogen, catalog number: C2925) in serum-free complete RPMI-1640 medium, or transfected with EGFP or EGFP-shootin1-DN (dominant negative shootin1)^39^ to visualize the entire cell body. CMFDA- or EGFP-labeled dendritic cells (1×10^5^ cells) were placed on the PDL- and laminin-coated polyacrylamide gels for 30 min at 37 ℃, 5 % CO_2_ to allow the adhesion to the gels. Cells were covered by the mixture of 1.5 mg/mL collagen gel + 10% Matrigel + CCL19 (20 or 200 ng/mL). The mixture was incubated at 37 ℃, 5 % CO_2_ for 30 min to allow polymerization of collagen fibers and gelation of Matrigel. To measure forces under glioblastoma cells, they were transfected with human shootin1b RNAi vector to co-express microRNA against human shootin1b RNA target sequence and EGFP to visualize the entire cell body. EGFP-labeled glioblastoma cells (1×10^5^ cells) were cultured overnight on the PDL- and laminin-coated polyacrylamide gels.

Time-lapse imaging of fluorescent nano beads and cells was performed at 37 ℃ using confocal microscope (LSM 710, Carl Zeiss) equipped with a C-Apochromat 63×/1.2 W Corr objective. Time-lapse images were acquired every 3 sec. The leading edge of a dendritic cell or the tumor microtube of a glioblastoma cell was determined by CMFDA or EGFP fluorescence images and DIC images. Traction forces under the cells were monitored by visualizing the force-induced deformation of the elastic substrate, which is reflected by the movement of the beads from their original positions^31^. The force vectors detected by beads under individual dendritic cells or under the tip of the tumor microtube of glioblastoma cells were then averaged, and were expressed as a vector composed of magnitude and angle (*θ*).

### Analysis of the correlation between the directions of force and cell migration

To analyze a possible correlation between the direction of traction force and the direction of dendritic cell migration, the 90 time-lapse images taken every 3 sec for 270 sec (Figure 1B) were divided into 3 groups: 1-90 sec (30 images), 91-180 sec (30 images), and 181-270 sec (30 images). The angle of the traction force was calculated for each image as described above, and then the average angle of the 30 images during the 90-sec observation in each group (*θ*1) was determined. The centroid of migrating cells in each image was calculated from the manually traced cell contour using Image J (Fiji), then the direction of cell migration during 90-sec observation in each group (*θ*2) was determined by the locations of the centroids in the 1st and 30th images. Thus, the 36 data in Figure 1E correspond to data from 12 cells (3 data from each cell).

### Analysis of dendritic cell migration in lymph node slice

Live imaging of dendritic cells in lymph node slices was performed as described^88,89^ with modifications. Isolated lymph nodes (inguinal, axial) were embedded in 4 % low gelling-temperature agarose (Sigma, catalog number: A0701) diluted in PBS. Agar-embedded lymph node tissue was sliced at 320 µm thickness using a vibratome (Leica) in a bath of ice-cold PBS. Lymph node slices were placed in a 35 mm culture dish containing the IMDM medium supplemented with 25mM HEPES, 10 % FBS, 100 U/mL Penicillin-100 µg/mL streptomycin, and then incubated at 37 ℃, 5 % CO_2_. WT and shootin1b KO dendritic cells were labeled with 1 µM CMFDA and 1 µM CMTPX, respectively, in serum-free complete RPMI-1640 medium for 30 min at 37 ℃, 5 % CO_2_. After washing with PBS, WT and shootin1b KO dendritic cells were resuspended in a 1:1 ratio (WT : KO) of 1×10^6^ cells/mL in the complete RPMI1640 medium. 100 µL of the mixed cell suspension was placed on the surface of a lymph node slice. The lymph node slice with cells was incubated at for 60 min at 37 ℃, 5 % CO_2_ to allow dendritic cell migration into the lymph node slice. After washing with PBS to remove the cells that did not migrate into the tissue, the lymph node slice was placed in glass bottom dish containing the complete RPMI1640 medium.

Dendritic cells in the lymph node slice were observed using a Leica SP8 Falcon microscope equipped with an HC PL APO 20×/0.75 dry objective lens and imaging software (LAS X) (Leica Microsystems). Fluorescence images were collected at more than 10 ∼ 25 µm below the surface of the lymph node slice. Fluorescence time-lapse images of x-y section with 10 µm z-spacing in total depth of 40 ∼ 50 µm were acquired at 37 ℃ every 30 sec for 60 min. In our perfusion system, a glass bottom dish containing the lymph node slice was placed in a custom chamber, and the oxygenated fresh RPMI1640 medium was delivered into the dish from one side using a peristaltic pump (ATTA, catalog number: SJ-1211) at a rate of 0.5 mL/min. The ascending medium was aspirated using another peristaltic pump connected to a waste collection flask. Thus, the medium in the glass bottom dish was continuously perfused with fresh RPMI1640 medium oxygenated with 95% O_2_・5% CO_2_ gas at 37℃ during time-lapse imaging.

### *In vivo* dendritic cell migration assay

WT and shootin1b KO dendritic cells were labeled with 1 µM CMTPX and 1 µM CMFDA, respectively, in serum-free RPMI1640 medium for 60 min. After washing with PBS, 3×10^6^ dendritic cells in a 1:1 ratio (WT:KO) were suspended in PBS and injected subcutaneously into the hind footpads of six-week-old C57BL/6 mice. Popliteal lymph nodes were removed after 24 hours and fixed by immersion in freshly prepared 4% paraformaldehyde at 4℃ for 60 min and frozen in OCT compound (SFJ, catalog number: 4583). 12-µm cryosections cut by a cryostat (Leica) were preincubated with 10 % normal goat serum (Vector, catalog number: S-1000) in phosphate buffer (PB) containing 0.3 % Triton-X 100 for 2 hours. The sections were then incubated with anti-pan-laminin antibody (from rabbit, 1:500 dilution) (Sigma, catalog number: L9393) diluted in PB containing 0.3 % Triton-X 100 at 4℃ overnight. Alexa 647 conjugated donkey anti-rabbit antibody (Abcam, catalog number: ab150075) was used as secondary antibody at a 1000-fold dilution overnight at 4℃. The sections were mounted with 50% (v/v) glycerol in PB. Fluorescence images were acquired using a confocal microscope (LSM710) equipped with a Plan-Apochromat ×10, 0.45 NA objective lens (Carl Zeiss). The lymph node was visualized with anti-pan-laminin antibody (Figure 5H). The ratio of WT or shootin1b KO dendritic cells migrated into the lymph node was calculated as the number of WT or KO cells divided by the total number of dendritic cells (WT + KO cells) in the T cell cortex of the lymph node. The average value of the ratio was calculated from three different layers of 12 µm cryosections in each experiment.

### Generation of CCL19 gradients for immunocytochemistry

A microfluidic device to generate CCL19 gradients in culture medium was fabricated as described^90^. Briefly, the device was fabricated with polydimethylsiloxane (PDMS; Silpot 184, Dow Corning Toray, catalog number: 3255981) and attached to a glass coverslip. The device consists of an open rectangular cell culture area and two microchannels on the long sides of the culture area. The micro-molds of the channel pattern were lithographically fabricated on a photoresist (SU-8 3025, MicroChem, USA) spin-coated on a 70-μm thick silicon wafer. PDMS sheets were obtained from this mold, which had been treated with silicone oil (Barrier coat No. 6, ShinEtsu, Japan) to facilitate their removal. The PDMS sheet was then bonded to a glass coverslip using plasma irradiation (Sakigake, catalog number: YHS-R). The glass coverslip was coated with PDL and laminin, and then dendritic cells were cultured on the 2-D cell culture area of the device for 1h to allow adhesion to the laminin-coated glass. To generate CCL19 gradient in the cell culture area, flow of complete RPMI1640 medium (7.5 µm/min) with or without 600 ng/mL CCL19^11^ were applied to the microchannels on either side of the cell culture area for 30 min using syringe pump.

## QUANTIFICATION AND STATISTICAL ANALYSIS

All statistical analysis were performed using Microsoft Excel (Microsoft Office LTSC professional Plus 2021 ver.) and Graphpad prism 7 (GraphPad Software). For samples with more than 7 data points, the D′Agostino–Pearson normality test was used to determine whether the data followed a normal distribution. In cases where the number of data points was between 3 and 7, the Shapiro‒Wilk test was used for the normality test. We also tested the equality of variation with the F-test for two independent groups that followed normal distributions. Significance tests were performed as follows: (1) two-tailed unpaired Student′s *t*-test to compare normally distributed data with equal variance from two independent groups; (2) two-tailed unpaired Welch′s *t*-test to compare normally distributed data with unequal variance from two independent groups; (3) two-tailed Mann–Whitney *U*-test to compare nonnormally distributed data from two independent groups. (4) For multiple comparisons, we used one-way ANOVA with Tukey’s post hoc test. For each experiment, the corresponding statistical information and number of samples are indicated in the figure legends. For detailed statistical results including the test statistics and p values, see the statistical source data associated with each figure. All data are shown as the mean ± SEM. Statistical significance was defined as ***p < 0.01; **p < 0.02; *p < 0.05; ns, not significant. All experiments were performed at least three times and reliably reproduced. Investigators were blinded to the experimental groups for each analysis, except biochemical analysis.

## REFERENCES

1. Wang, Y.L. (1985). Exchange of actin subunits at the leading edge of living fibroblasts: possible role of treadmilling. J. Cell Biol. 101, 597–602.

2. Theriot, J.A., and Mitchison, T.J. (1991). Actin microfilament dynamics in locomoting cells. Nature 352, 126–131.

3. Mitchison, T., and Kirschner, M. (1988). Cytoskeletal dynamics and nerve growth. Neuron 1, 761–772.

4. Case, L.B., and Waterman, C.M. (2015). Integration of actin dynamics and cell adhesion by a three-dimensional, mechanosensitive molecular clutch. Nat. Cell Biol. 17, 955–963.

5. Hu, K., Ji, L., Applegate, K.T., Danuser, G., and Waterman-Storer, C.M. (2007). Differential transmission of actin motion within focal adhesions. Science 315, 111–115.

6. Zhang, X., Jiang, G., Cai, Y., Monkley, S.J., Critchley, D.R., and Sheetz, M.P. (2008). Talin depletion reveals independence of initial cell spreading from integrin activation and traction. Nat. Cell Biol. 10, 1062–1068.

7. Thievessen, I., Thompson, P.M., Berlemont, S., Plevock, K.M., Plotnikov, S.V., Zemljic-Harpf, A., Ross, R.S., Davidson, M.W., Danuser, G., Campbell, S.L., and Waterman, C.M. (2013). Vinculin-actin interaction couples actin retrograde flow to focal adhesions, but is dispensable for focal adhesion growth. J. Cell Biol. 202, 163–177.

8. Moore, S.W., Roca-Cusachs, P., and Sheetz, M.P. (2010). Stretchy proteins on stretchy substrates: the important elements of integrin-mediated rigidity sensing. Dev Cell 19, 194–206.

9. Elosegui-Artola, A., Oria, R., Chen, Y., Kosmalska, A., Perez-Gonzalez, C., Castro, N., Zhu, C., Trepat, X., and Roca-Cusachs, P. (2016). Mechanical regulation of a molecular clutch defines force transmission and transduction in response to matrix rigidity. Nat. Cell Biol. 18, 540–548.

10. Friedl, P., Entschladen, F., Conrad, C., Niggemann, B., and Zanker, K.S. (1998). CD4+ T lymphocytes migrating in three-dimensional collagen lattices lack focal adhesions and utilize beta1 integrin-independent strategies for polarization, interaction with collagen fibers and locomotion. Eur. J. Immunol. 28, 2331–2343.

11. Lammermann, T., Bader, B.L., Monkley, S.J., Worbs, T., Wedlich-Soldner, R., Hirsch, K., Keller, M., Forster, R., Critchley, D.R., Fassler, R., and Sixt, M. (2008). Rapid leukocyte migration by integrin-independent flowing and squeezing. Nature 453, 51–55.

12. Paluch, E.K., Aspalter, I.M., and Sixt, M. (2016). Focal Adhesion-Independent Cell Migration. Annu. Rev. Cell Dev. Biol. 32, 469–490.

13. Paterson, N., and Lammermann, T. (2022). Macrophage network dynamics depend on haptokinesis for optimal local surveillance. Elife 11, e75354.

14. Schmidt, S., and Friedl, P. (2010). Interstitial cell migration: integrin-dependent and alternative adhesion mechanisms. Cell Tissue Res. 339, 83–92.

15. Bergert, M., Erzberger, A., Desai, R.A., Aspalter, I.M., Oates, A.C., Charras, G., Salbreux, G., and Paluch, E.K. (2015). Force transmission during adhesion-independent migration. Nat. Cell Biol. 17, 524–529.

16. Hons, M., Kopf, A., Hauschild, R., Leithner, A., Gaertner, F., Abe, J., Renkawitz, J., Stein, J.V., and Sixt, M. (2018). Chemokines and integrins independently tune actin flow and substrate friction during intranodal migration of T cells. Nat. Immunol. 19, 606–616.

17. Ruprecht, V., Wieser, S., Callan-Jones, A., Smutny, M., Morita, H., Sako, K., Barone, V., Ritsch-Marte, M., Sixt, M., Voituriez, R., and Heisenberg, C.P. (2015). Cortical contractility triggers a stochastic switch to fast amoeboid cell motility. Cell 160, 673–685.

18. Liu, Y.J., Le Berre, M., Lautenschlaeger, F., Maiuri, P., Callan-Jones, A., Heuze, M., Takaki, T., Voituriez, R., and Piel, M. (2015). Confinement and low adhesion induce fast amoeboid migration of slow mesenchymal cells. Cell 160, 659–672.

19. Hawkins, R.J., Poincloux, R., Benichou, O., Piel, M., Chavrier, P., and Voituriez, R. (2011). Spontaneous contractility-mediated cortical flow generates cell migration in three-dimensional environments. Biophys. J. 101, 1041–1045.

20. Aoun, L., Farutin, A., Garcia-Seyda, N., Negre, P., Rizvi, M.S., Tlili, S., Song, S., Luo, X., Biarnes-Pelicot, M., Galland, R., et al. (2020). Amoeboid swimming is propelled by molecular paddling in lymphocytes. Biophys. J. 119, 1157–1177.

21. Yamada, K.M., and Sixt, M. (2019). Mechanisms of 3D cell migration. Nat. Rev. Mol. Cell Biol. 20, 738–752.

22. Lammermann, T., and Sixt, M. (2009). Mechanical modes of ’amoeboid’ cell migration. Curr. Opin. Cell Biol. 21, 636–644.

23. Friedl, P. (2004). Prespecification and plasticity: shifting mechanisms of cell migration. Curr. Opin. Cell Biol. 16, 14–23.

24. Cuddapah, V.A., Robel, S., Watkins, S., and Sontheimer, H. (2014). A neurocentric perspective on glioma invasion. Nat. Rev. Neurosci. 15, 455–465.

25. Tetzlaff, S.K., Reyhan, E., Layer, N., Bengtson, C.P., Heuer, A., Schroers, J., Faymonville, A.J., Langeroudi, A.P., Drewa, N., Keifert, E., et al. (2025). Characterizing and targeting glioblastoma neuron-tumor networks with retrograde tracing. Cell 188, 1–22.

26. Toriyama, M., Shimada, T., Kim, K.B., Mitsuba, M., Nomura, E., Katsuta, K., Sakumura, Y., Roepstorff, P., and Inagaki, N. (2006). Shootin1: A protein involved in the organization of an asymmetric signal for neuronal polarization. J. Cell Biol. 175, 147–157.

27. Shimada, T., Toriyama, M., Uemura, K., Kamiguchi, H., Sugiura, T., Watanabe, N., and Inagaki, N. (2008). Shootin1 interacts with actin retrograde flow and L1-CAM to promote axon outgrowth. J. Cell Biol. 181, 817–829.

28. Kubo, Y., Baba, K., Toriyama, M., Minegishi, T., Sugiura, T., Kozawa, S., Ikeda, K., and Inagaki, N. (2015). Shootin1-cortactin interaction mediates signal-force transduction for axon outgrowth. J. Cell Biol. 210, 663–676.

29. Kastian, R.F., Minegishi, T., Baba, K., Saneyoshi, T., Katsuno-Kambe, H., Saranpal, S., Hayashi, Y., and Inagaki, N. (2021). Shootin1a-mediated actin-adhesion coupling generates force to trigger structural plasticity of dendritic spines. Cell Rep. 35, 109130.

30. Higashiguchi, Y., Katsuta, K., Minegishi, T., Yonemura, S., Urasaki, A., and Inagaki, N. (2016). Identification of a shootin1 isoform expressed in peripheral tissues. Cell Tissue Res. 366, 75–87.

31. Minegishi, T., Kastian, R., and Inagaki, N. (2021). Single speckle imaging using a standard epi-fluorescent microscope and traction force microscopy at nerve growth cones. J. Vis. Exp., e63227.

32. Maddaluno, L., Verbrugge, S.E., Martinoli, C., Matteoli, G., Chiavelli, A., Zeng, Y., Williams, E.D., Rescigno, M., and Cavallaro, U. (2009). The adhesion molecule L1 regulates transendothelial migration and trafficking of dendritic cells. J. Exp. Med. 206, 623–635.

33. Hall, H., Carbonetto, S., and Schachner, M. (1997). L1/HNK-1 carbohydrate- and beta 1 integrin-dependent neural cell adhesion to laminin-1. J. Neurochem. 68, 544–553.

34. Colombo, F., and Meldolesi, J. (2015). L1-CAM and N-CAM: From adhesion proteins to pharmacological targets. Trends Pharmacol. Sci. 36, 769–781.

35. Sixt, M., Kanazawa, N., Selg, M., Samson, T., Roos, G., Reinhardt, D.P., Pabst, R., Lutz, M.B., and Sorokin, L. (2005). The conduit system transports soluble antigens from the afferent lymph to resident dendritic cells in the T cell area of the lymph node. Immunity 22, 19–29.

36. Drumea-Mirancea, M., Wessels, J.T., Muller, C.A., Essl, M., Eble, J.A., Tolosa, E., Koch, M., Reinhardt, D.P., Sixt, M., Sorokin, L., et al. (2006). Characterization of a conduit system containing laminin-5 in the human thymus: a potential transport system for small molecules. J. Cell Sci. 119, 1396–1405.

37. Lammermann, T., Renkawitz, J., Wu, X., Hirsch, K., Brakebusch, C., and Sixt, M. (2009). Cdc42-dependent leading edge coordination is essential for interstitial dendritic cell migration. Blood 113, 5703–5710.

38. Ricart, B.G., Yang, M.T., Hunter, C.A., Chen, C.S., and Hammer, D.A. (2011). Measuring traction forces of motile dendritic cells on micropost arrays. Biophys. J. 101, 2620–2628.

39. Baba, K., Yoshida, W., Toriyama, M., Shimada, T., Manning, C.F., Saito, M., Kohno, K., Trimmer, J.S., Watanabe, R., and Inagaki, N. (2018). Gradient-reading and mechano-effector machinery for netrin-1-induced axon guidance. Elife 7, e34593.

40. Renkawitz, J., Schumann, K., Weber, M., Lammermann, T., Pflicke, H., Piel, M., Polleux, J., Spatz, J.P., and Sixt, M. (2009). Adaptive force transmission in amoeboid cell migration. Nat. Cell Biol. 11, 1438–1443.

41. Sanchez-Sanchez, N., Riol-Blanco, L., and Rodriguez-Fernandez, J.L. (2006). The multiple personalities of the chemokine receptor CCR7 in dendritic cells. J. Immunol. 176, 5153–5159.

42. Lu, M., Xu, C., Zhang, Q., Wu, X., Tang, L., Wang, X., Wu, J., and Wu, X. (2018). Inhibition of p21-activated kinase 1 attenuates the cardinal features of asthma through suppressing the lymph node homing of dendritic cells. Biochem. Pharmacol. 154, 464–473.

43. Toriyama, M., Kozawa, S., Sakumura, Y., and Inagaki, N. (2013). Conversion of a signal into forces for axon outgrowth through Pak1-mediated shootin1 phosphorylation. Curr. Biol. 23, 529–534.

44. Alvarez, D., Vollmann, E.H., and von Andrian, U.H. (2008). Mechanisms and consequences of dendritic cell migration. Immunity 29, 325–342.

45. Pfeiffer, F., Kumar, V., Butz, S., Vestweber, D., Imhof, B.A., Stein, J.V., and Engelhardt, B. (2008). Distinct molecular composition of blood and lymphatic vascular endothelial cell junctions establishes specific functional barriers within the peripheral lymph node. Eur. J. Immunol. 38, 2142–2155.

46. Sobocinski, G.P., Toy, K., Bobrowski, W.F., Shaw, S., Anderson, A.O., and Kaldjian, E.P. (2010). Ultrastructural localization of extracellular matrix proteins of the lymph node cortex: evidence supporting the reticular network as a pathway for lymphocyte migration. BMC Immunol. 11, 42.

47. Haessler, U., Pisano, M., Wu, M., and Swartz, M.A. (2011). Dendritic cell chemotaxis in 3D under defined chemokine gradients reveals differential response to ligands CCL21 and CCL19. Proc. Natl. Acad. Sci. U. S. A. 108, 5614–5619.

48. Giannone, G., Mege, R.M., and Thoumine, O. (2009). Multi-level molecular clutches in motile cell processes. Trends Cell Biol. 19, 475–486.

49. Reversat, A., Gaertner, F., Merrin, J., Stopp, J., Tasciyan, S., Aguilera, J., de Vries, I., Hauschild, R., Hons, M., Piel, M., et al. (2020). Cellular locomotion using environmental topography. Nature 582, 582–585.

50. Osswald, M., Jung, E., Sahm, F., Solecki, G., Venkataramani, V., Blaes, J., Weil, S., Horstmann, H., Wiestler, B., Syed, M., et al. (2015). Brain tumour cells interconnect to a functional and resistant network. Nature 528, 93–98.

51. Venkataramani, V., Yang, Y., Schubert, M.C., Reyhan, E., Tetzlaff, S.K., Wissmann, N., Botz, M., Soyka, S.J., Beretta, C.A., Pramatarov, R.L., et al. (2022). Glioblastoma hijacks neuronal mechanisms for brain invasion. Cell 185, 2899–2917 e2831.

52. Maaser, K., Wolf, K., Klein, C.E., Niggemann, B., Zanker, K.S., Brocker, E.B., and Friedl, P. (1999). Functional hierarchy of simultaneously expressed adhesion receptors: integrin alpha2beta1 but not CD44 mediates MV3 melanoma cell migration and matrix reorganization within three-dimensional hyaluronan-containing collagen matrices. Mol. Biol. Cell 10, 3067–3079.

53. Kaltenbach, L., Martzloff, P., Bambach, S.K., Aizarani, N., Mihlan, M., Gavrilov, A., Glaser, K.M., Stecher, M., Thunauer, R., Thiriot, A., et al. (2023). Slow integrin-dependent migration organizes networks of tissue-resident mast cells. Nat. Immunol. 24, 915–924.

54. Aratyn-Schaus, Y., and Gardel, M.L. (2010). Transient frictional slip between integrin and the ECM in focal adhesions under myosin II tension. Curr. Biol. 20, 1145–1153.

55. del Rio, A., Perez-Jimenez, R., Liu, R., Roca-Cusachs, P., Fernandez, J.M., and Sheetz, M.P. (2009). Stretching single talin rod molecules activates vinculin binding. Science 323, 638–641.

56. Changede, R., Xu, X., Margadant, F., and Sheetz, M.P. (2015). Nascent integrin adhesions form on all matrix rigidities after integrin activation. Dev. Cell 35, 614–621.

57. Sun, Z., Guo, S.S., and Fassler, R. (2016). Integrin-mediated mechanotransduction. J. Cell Biol. 215, 445–456.

58. Elosegui-Artola, A., Trepat, X., and Roca-Cusachs, P. (2018). Control of mechanotransduction by molecular clutch dynamics. Trends Cell Biol. 28, 356–367.

59. Palecek, S.P., Loftus, J.C., Ginsberg, M.H., Lauffenburger, D.A., and Horwitz, A.F. (1997). Integrin-ligand binding properties govern cell migration speed through cell-substratum adhesiveness. Nature 385, 537–540.

60. Kuboki, T., Ebata, H., Matsuda, T., Arai, Y., Nagai, T., and Kidoaki, S. (2020). Hierarchical development of motile polarity in durotactic cells just crossing an elasticity boundary. Cell Struct. Funct. 45, 33–43.

61. Petrie, R.J., Koo, H., and Yamada, K.M. (2014). Generation of compartmentalized pressure by a nuclear piston governs cell motility in a 3D matrix. Science 345, 1062–1065.

62. Stroka, K.M., Jiang, H., Chen, S.H., Tong, Z., Wirtz, D., Sun, S.X., and Konstantopoulos, K. (2014). Water permeation drives tumor cell migration in confined microenvironments. Cell 157, 611–623.

63. Qiu, Z., Minegishi, T., Aoki, D., Abe, K., Baba, K., and Inagaki, N. (2024). Adhesion-clutch between DCC and netrin-1 mediates netrin-1-induced axonal haptotaxis. Front. Mol. Neurosci. 17, 1307755.

64. Wu, D. (2005). Signaling mechanisms for regulation of chemotaxis. Cell Res. 15, 52–56.

65. Lawson, C.D., and Ridley, A.J. (2018). Rho GTPase signaling complexes in cell migration and invasion. J. Cell Biol. 217, 447–457.

66. Weiss, F., Lauffenburger, D., and Friedl, P. (2022). Towards targeting of shared mechanisms of cancer metastasis and therapy resistance. Nat. Rev. Cancer 22, 157–173.

67. Demuth, T., and Berens, M.E. (2004). Molecular mechanisms of glioma cell migration and invasion. J. Neurooncol. 70, 217–228.

68. Watkins, S., Robel, S., Kimbrough, I.F., Robert, S.M., Ellis-Davies, G., and Sontheimer, H. (2014). Disruption of astrocyte-vascular coupling and the blood-brain barrier by invading glioma cells. Nat, Commun, 5, 4196.

69. Drumm, M.R., Dixit, K.S., Grimm, S., Kumthekar, P., Lukas, R.V., Raizer, J.J., Stupp, R., Chheda, M.G., Kam, K.L., McCord, M., et al. (2020). Extensive brainstem infiltration, not mass effect, is a common feature of end-stage cerebral glioblastomas. Neuro. Oncol. 22, 470–479.

70. Fields, R.D. (2015). A new mechanism of nervous system plasticity: activity-dependent myelination. Nat. Rev. Neurosci. 16, 756–767.

71. Lau, L.W., Cua, R., Keough, M.B., Haylock-Jacobs, S., and Yong, V.W. (2013). Pathophysiology of the brain extracellular matrix: a new target for remyelination. Nat. Rev. Neurosci. 14, 722–729.

72. Neftel, C., Laffy, J., Filbin, M.G., Hara, T., Shore, M.E., Rahme, G.J., Richman, A.R., Silverbush, D., Shaw, M.L., Hebert, C.M., et al. (2019). An integrative model of cellular states, plasticity, and genetics for glioblastoma. Cell 178, 835–849 e821.

73. Winkler, F., Venkatesh, H.S., Amit, M., Batchelor, T., Demir, I.E., Deneen, B., Gutmann, D.H., Hervey-Jumper, S., Kuner, T., Mabbott, D., et al. (2023). Cancer neuroscience:: state of the field, emerging directions. Cell 186, 1689–1707.

74. Ostrom, Q.T., Bauchet, L., Davis, F.G., Deltour, I., Fisher, J.L., Langer, C.E., Pekmezci, M., Schwartzbaum, J.A., Turner, M.C., Walsh, K.M., et al. (2014). The epidemiology of glioma in adults: a “state of the science” review. Neuro. Oncol. 16, 896–913.

75. Weller, M., Wick, W., Aldape, K., Brada, M., Berger, M., Pfister, S.M., Nishikawa, R., Rosenthal, M., Wen, P.Y., Stupp, R., and Reifenberger, G. (2015). Glioma. Nat. Rev. Dis. Primers 1, 15017.

76. Lutz, M.B., Kukutsch, N., Ogilvie, A.L., Rossner, S., Koch, F., Romani, N., and Schuler, G. (1999). An advanced culture method for generating large quantities of highly pure dendritic cells from mouse bone marrow. J. Immunol. Methods 223, 77–92.

77. Wang, W., Li, J., Wu, K., Azhati, B., and Rexiati, M. (2016). Culture and Identification of Mouse Bone Marrow-Derived Dendritic Cells and Their Capability to Induce T Lymphocyte Proliferation. Med. Sci. Monit. 22, 244–250.

78. Bamba, Y., Shofuda, T., Kanematsu, D., Nonaka, M., Yamasaki, M., Okano, H., and Kanemura, Y. (2014). Differentiation, polarization, and migration of human induced pluripotent stem cell-derived neural progenitor cells co-cultured with a human glial cell line with radial glial-like characteristics. Biochem. Biophys. Res. Commun. 447, 683–688.

79. Eino, D., Tsukada, Y., Naito, H., Kanemura, Y., Iba, T., Wakabayashi, T., Muramatsu, F., Kidoya, H., Arita, H., Kagawa, N., et al. (2018). LPA4-Mediated Vascular Network Formation Increases the Efficacy of Anti-PD-1 Therapy against Brain Tumors. Cancer Res. 78, 6607–6620.

80. Kim, H., Sa, J.K., Kim, J., Cho, H.J., Oh, H.J., Choi, D.H., Kang, S.H., Jeong, D.E., Nam, D.H., Lee, H., et al. (2022). Recapitulated crosstalk between cerebral metastatic lung cancer cells and brain perivascular tumor microenvironment in a microfluidic co-culture chip. Adv Sci (Weinh) 9, e2201785.

81. Inagaki, N., Fukui, H., Taguchi, Y., Wang, N.P., Yamatodani, A., and Wada, H. (1989). Characterization of histamine H1-receptors on astrocytes in primary culture: [3H]mepyramine binding studies. Eur. J. Pharmacol. 173, 43–51.

82. Io, S., Iemura, Y., and Takashima, Y. (2021). Optimized protocol for naive human pluripotent stem cell-derived trophoblast induction. STAR Protoc. 2, 100921.

83. Watanabe, M., Buth, J.E., Vishlaghi, N., de la Torre-Ubieta, L., Taxidis, J., Khakh, B.S., Coppola, G., Pearson, C.A., Yamauchi, K., Gong, D., et al. (2017). Self-organized cerebral organoids with human-specific features predict effective drugs to combat Zika virus infection. Cell Rep. 21, 517–532.

84. Lancaster, M.A., and Knoblich, J.A. (2014). Generation of cerebral organoids from human pluripotent stem cells. Nat. Protoc. 9, 2329–2340.

85. Minegishi, T., Uesugi, Y., Kaneko, N., Yoshida, W., Sawamoto, K., and Inagaki, N. (2018). Shootin1b mediates a mechanical clutch to produce force for neuronal migration. Cell Rep. 25, 624–639.e626.

86. Azizi, L., Cowell, A.R., Mykuliak, V.V., Goult, B.T., Turkki, P., and Hytonen, V.P. (2021). Cancer associated talin point mutations disorganise cell adhesion and migration. Sci. Rep. 11, 347.

87. Abe, K., Katsuno, H., Toriyama, M., Baba, K., Mori, T., Hakoshima, T., Kanemura, Y., Watanabe, R., and Inagaki, N. (2018). Grip and slip of L1-CAM on adhesive substrates direct growth cone haptotaxis. Proc. Natl. Acad. Sci. U. S. A. 115, 2764–2769.

88. Salmon, H., Rivas-Caicedo, A., Asperti-Boursin, F., Lebugle, C., Bourdoncle, P., and Donnadieu, E. (2011). Ex vivo imaging of T cells in murine lymph node slices with widefield and confocal microscopes. J. Vis. Exp., e3054.

89. Katakai, T., Habiro, K., and Kinashi, T. (2013). Dendritic cells regulate high-speed interstitial T cell migration in the lymph node via LFA-1/ICAM-1. J. Immunol. 191, 1188–1199.

90. Kastian, R.F., Baba, K., Kaewkascholkul, N., Sasaki, H., Watanabe, R., Toriyama, M., and Inagaki, N. (2023). Dephosphorylation of neural wiring protein shootin1 by PP1 phosphatase regulates netrin-1-induced axon guidance. J. Biol. Chem. 299, 104687.

