## Supplemental information for "Weak and tunable adhesion-clutch drives rapid cell migration and glioblastoma motility"

**This file includes:**

Supplementary Figures S1-S9, Supplementary Videos S1-S13

### Supplementary Figure legends

**Figure S1. Shootin1b promotes dendritic cell migration under CCL19 signaling**

(A, D) Dendritic cells were cultured in a mixture of collagen gel and Matrigel. One hour after the bath application of 200 ng/mL CCL19, time-lapse phase-contrast/fluorescence images of WT (A) and shootin1b KO (D) dendritic cells were obtained. Nuclei were visualized by Hoechst to accurately trace the trajectories of cell migrations (see Video S2). The pictures show representative images from the time-lapse series taken every 1 min for 180 min. Scale bars: 50 μm.

(B, C, E, F) Trajectories of the migrations of WT (B, C) and shootin1b KO (E, F) dendritic cells in the presence of 20 ng/mL (B, E) and 200 ng/mL (C, F) CCL19. The initial cell positions are normalized at x = 0 μm and y = 0 μm.

**Figure S2. Expression and localization of shootin1b in dendritic cells**

(A) Immunoblot analysis of dendritic cells with anti-shootin1a and anti-shootin1b antibodies. Embryonic day 16.5 mouse brain lysate was used as positive control for the detection of shootin1a and shootin1b. Actin was used as a loading control.

(B) Fluorescence images of dendritic cells co-stained with anti-CD11c (dendritic cell marker) and anti-shootin1b or anti-shootin1a antibody. Cell nucleus was visualized by DAPI. Scale bar: 10 μm (in the inset, 2 μm).

(C, D) Fluorescence images of dendritic cells co-stained with anti-shootin1b antibody and anti-cortactin (C) or anti-L1 antibody (D). Enlarged views of the rectangular region are shown to the right. Arrowheads indicate shootin1b localization with cortactin (C) or L1 (D) in filopodia. The images were obtained by STED microscopy. Scale bar: 10 μm (in the inset, 2 μm).

(E, F) Shootin1b KO does not affect the expression of the CCL19 receptor CCR7 in dendritic cells. Immunoblot analysis of WT and shootin1b KO dendritic cells with anti-CCR7 antibody (E). Actin was used as a loading control. Quantitative data for the CCR7 expression level in (F). Two-tailed unpaired Welch's *t*-test was performed (n = 3 independent experiments). Data represent means ± SEM; ns, not significant.

**Figure S3. Shootin1b-L1 interaction mediates generation of weak forces by dendritic cells**

(A, B) Overlayed DIC and fluorescence images showing dendritic cells overexpressing EGFP (control) (A) or EGFP-shootin1-DN (B) migrating under the semi-3D condition in (Figure 1A) in the presence of 200 ng/mL CCL19. See Video S4. The pictures show representative images from the time-lapse series taken every 3 sec for 270 sec. The original and displaced positions of the beads in polyacrylamide gels are indicated by green and red colors, respectively. The cells were visualized by EGFP (blue color); dashed lines indicate the boundaries of the cells. The kymographs (panel below) along the axis of bead displacement (white dashed arrows) at indicated areas 1 and 2 show movement of beads recorded by every 3 sec. The bead in area 2 is a reference bead. Scale bar: 5 μm (in the inset, 1 μm).

(C) Analyses of magnitude of the traction force under the dendritic cells overexpressing EGFP or EGFP-shootin1-DN in (A, B). Two-tailed unpaired Student's *t*-test (EGFP, n = 10 cells; EGFP-shootin1-DN, n = 10 cells).

**Figure S4. Pak1-mediated shootin1b phosphorylation under CCL19 signaling**

(A, B) Dendritic cells were treated with 200 ng/mL CCL19 for 0, 10, 30 or 60 min. Cell lysates were then analyzed by immunoblot with anti-pSer101 shootin1, anti-pSer249 shootin1 and anti-shootin1b antibodies (A). Quantitative data for phospho-shootin1b levels at 0, 10, 30 or 60 min are shown in (B). For multiple comparison, one-way ANOVA with Turkey's post hoc test was performed (n = 3 independent experiments).

(C, D) Dendritic cells were treated with 200 ng/mL CCL19 or 0.1 % BSA (for control) for 30 min (C). To inhibit Pak1 function, 0.25 μM Pak1 inhibitor (NVS-PAK-1) was applied for 30 min before the CCL19 treatment. Cell lysates were then analyzed by immunoblot with anti-pSer101 shootin1, anti-pSer249 shootin1 and anti-shootin1b antibodies. Quantitative data for phospho-shootin1b levels are shown in (D). For multiple comparison, one-way ANOVA with Turkey's post hoc test was performed (n = 3 independent experiments).

Data represent means ± SEM; *, $p < 0.05$; ***, $p < 0.01$.

**Figure S5. Shootin1b-L1 interaction mediates CCL19-induced dendritic cell chemotaxis**

(A, B) A gradient of CCL19 was applied to dendritic cells overexpressing EGFP (control) (A) or EGFP-shootin1-DN (B) cultured in a mixture of collagen gel and Matrigel (left panel, Figure 4A). The graphs depict the migration trajectories of individual dendritic cells overexpressing EGFP (A) or EGFP-shootin1-DN (B). The initial cell positions are normalized at x = 0 μm and y = 0 μm. See also Video S8.

(C, D) Analyses of migration speed (C) and chemotaxis index (D) for dendritic cells overexpressing EGFP (control) (A) and EGFP-shootin1-DN (B). Two-tailed Mann-Whitney *U*-test was performed (EGFP, n = 71 cells; EGFP-shootin1-DN, n = 54 cells).

Data represent means ± SEM; ***, $p < 0.01$.

**Figure S6. Shootin1b does not interact with β1- or β2-integrin**

(A) Co-immunoprecipitation assay of AcGFP (control), AcGFP-integrin β1-ICD (intracellular domain), AcGFP-integrin β2-ICD and AcGFP-L1-ICD with flag-shootin1b. Cell lysates from HEK293T cells, expressing AcGFP-tagged proteins and flag-shootin1b, were incubated with anti-flag antibody. The immunoprecipitates and cell lysates (1 %) were immunoblotted with anti-GFP or anti-flag antibody. AcGFP-L1-ICD, but not AcGFP-integrin β1-ICD or AcGFP-integrin β2-ICD, was co-precipitated with flag-shootin1b.

(B) HEK293T cells were co-transfected with mouse talin1-EGFP vector and candidate mouse talin1 RNAi vectors (#1 ~ #5). Cell lysates were then analyzed by immunoblot with anti-GFP. Actin was used as a loading control. Talin1 RNAi vector #3 was used in the knockdown analysis.

**Figure S7. Shootin1b KO inhibits dendritic cell migration in lymph node slice**

(A) Fluorescence images of WT (green) or shootin1b KO (magenta) cells migrating in a lymph node slice. Enlarged images in the rectangle area are shown to the right; the pictures show representative time-lapse images from the time-lapse series taken every 30 sec for 60 min. See Video S10. Scale bar: 50 μm (in the inset, 20 μm).

(B) Analyses of the migration speeds of WT and shootin1b KO dendritic cells in lymph node slices in (A). Two-tailed unpaired Student's *t*-test was performed (WT, $n = 56$ cells; KO, $n = 55$ cells).

Data represent means ± SEM; **, $p < 0.02$.

**Figure S8. Expression of shootin1b, cortactin and L1 in human glioblastoma cells**

(A) Immunoblot analysis of human glioblastoma cells with anti-shootin1b, anti-shootin1a, anti-L1 and anti-cortactin antibodies. Actin was used as a loading control. Embryonic day 16.5 mouse brain lysate was used as positive control for the detection of endogenous shootin1a, shootin1b, L1 and cortactin.

(B) HEK293T cells was transfected with human shootin1b RNAi candidate vectors (#1 ~ #6). Cell lysates were then analyzed by immunoblot with anti-shootin1b. Actin was used as a loading control. We used shootin1b RNAi vector #3 to knockdown shootin1b in glioblastoma cells in Figures 6, 7B-C.

(C) Human glioblastoma cells were infected with lentivirus carrying shScramble (control RNAi) or candidates of shShootin1b (shootin1b RNAi #7, #8). Glyceraldehyde-3-

phosphate dehydrogenase (GAPDH) was used as a loading control. We used shootin1b RNAi vector #7 to knockdown shootin1b in glioblastoma cells in Figure 7H-I.

**Figure S9. Shootin1b KO does not affect dendritic cell maturation**

(A-D) Flow cytometry analysis of the expression of dendritic cell surface markers on WT and shootin1b KO dendritic cells after maturation. Dendritic cell surface markers against CD11c (B), MHC-II (C), CD80 (D) were detected with FITC-conjugated antibodies. The corresponding isotype control antibody did not bind to dendritic cell surface (A). Dendritic cell population was detected as dot plot. The x and y axis of graphs indicate FITC intensity and backscatter coefficient, respectively. Magenta rectangles in the graphs indicate FITC-negative cell population (A). Green rectangles indicate FITC-positive cell population (B-D). Numbers within magenta and green rectangles represent the percentage of FITC-negative and FITC-positive cell population to the total cell number, respectively.

### Supplementary Video legends

**Video S1. A time-lapse fluorescence movie of traction forces exerted by a dendritic cell migrating in the presence of 200 ng/mL CCL19.** Blue lines indicate the movement of the beads from their original positions. Yellow arrows indicate the direction and relative amplitude of traction forces. See the legend for Figure 1B.

**Video S2. Time-lapse fluorescence movies of dendritic cells migrating in collagen gel + Matrigel in the presence of 20 and 200 ng/mL CCL19.** Nuclei were visualized by Hoechst to accurately trace the trajectories of cell migrations. See the legend for Figure S1A-C.

**Video S3. A time-lapse fluorescence movies of traction forces exerted by WT and shootin1b KO dendritic cells migrating in the presence of 200 ng/mL CCL19.** See the legends for Figure 1B, G.

**Video S4. A time-lapse fluorescence movies of traction forces exerted by dendritic cells expressing EGFP (control) or EGFP-shootin1-DN and migrating in the presence of 200 ng/mL CCL19.** See the legends for Figure S3A-B.

**Video S5. Time-lapse fluorescence movies of WT and shootin1b KO dendritic cells migrating in collagen gel + Matrigel in the presence of 200 ng/mL CCL19.** Nuclei were visualized by Hoechst to trace the trajectories of cell migrations. See the legend for Figure S1A, D.

**Video S6. Movement of fluorescent speckles of HaloTag-actin, HaloTag-shootin1b and L1-halotag at the leading edge of dendritic cells on laminin (LN)- or PDL-coated glass.** See the legend for Figure 2A-C, F-H.

**Video S7. Time-lapse fluorescence movies of HaloTag-actin and Lifeact-EGFP in the leading edge of WT and shootin1b KO dendritic cells.** See the legend for Figure 3E.

**Video S8. Time-lapse fluorescence movies of WT and shootin1b KO dendritic cells and WT dendritic cells overexpressing EGFP (control) or EGFP-shootin1-DN migrating under CCL19 gradient.** See the legend for Figures 4A-B, S5. Nuclei were visualized by Hoechst to trace the trajectories of cell migrations.

**Video S9. Time-lapse fluorescence movies of WT and shootin1b KO dendritic cells expressing control RNAi or talin1 RNAi migrating under CCL19 gradient.** Nuclei were visualized by Hoechst to trace the trajectories of cell migrations. See the legend for Figure 5D.

**Video S10. A time-lapse fluorescence movie of WT and shootin1b KO dendritic cells migrating in a lymph node slice.** See the legend for Figure S7A.

**Video S11. Time-lapse movies of migration of a human astrocyte and a glioblastoma cell, glioblastoma cells expressing control RNAi or shootin1b RNAi, and glioblastoma cells expressing EGFP (control) or EGFP-shootin1.** See the legend for Figures 6C, 7A-B, D-E.

**Video S12. Movement of fluorescent speckles of HaloTag-actin at the tip of the tumor microtube of glioblastoma cells expressing control RNAi and shootin1b RNAi.** See the legend for Figure 6E.

**Video S13. A time-lapse fluorescence movie of traction force at the tip of the tumor microtube of glioblastoma cells expressing control RNAi and shootin1b RNAi.** See the legend for Figure 6G-H.
