## Supplementary figures and images for "Weak and tunable adhesion-clutch drives rapid cell migration and glioblastoma motility"

### Supplementary Figures S1-S9

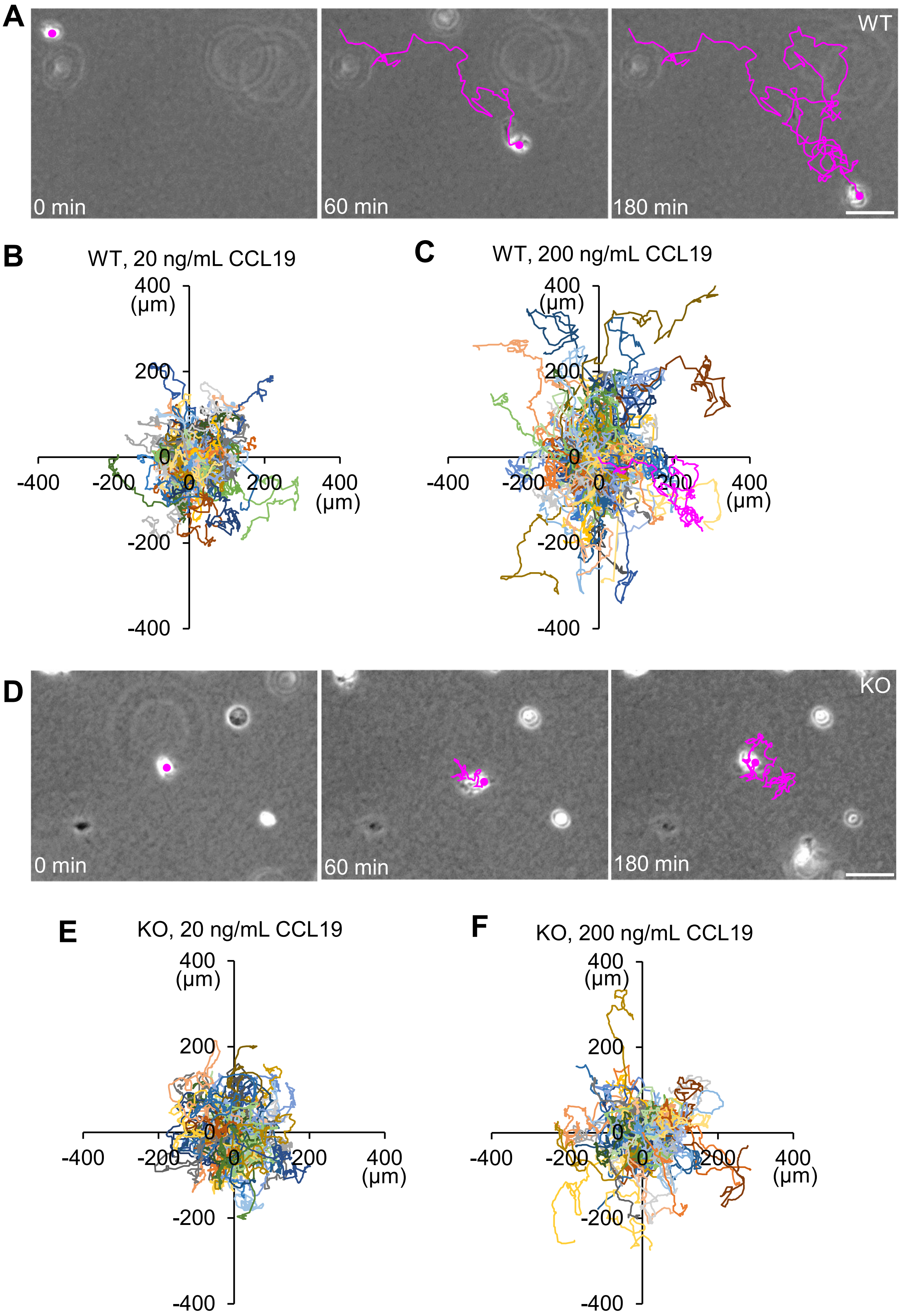

Figure S2 Baba et al.

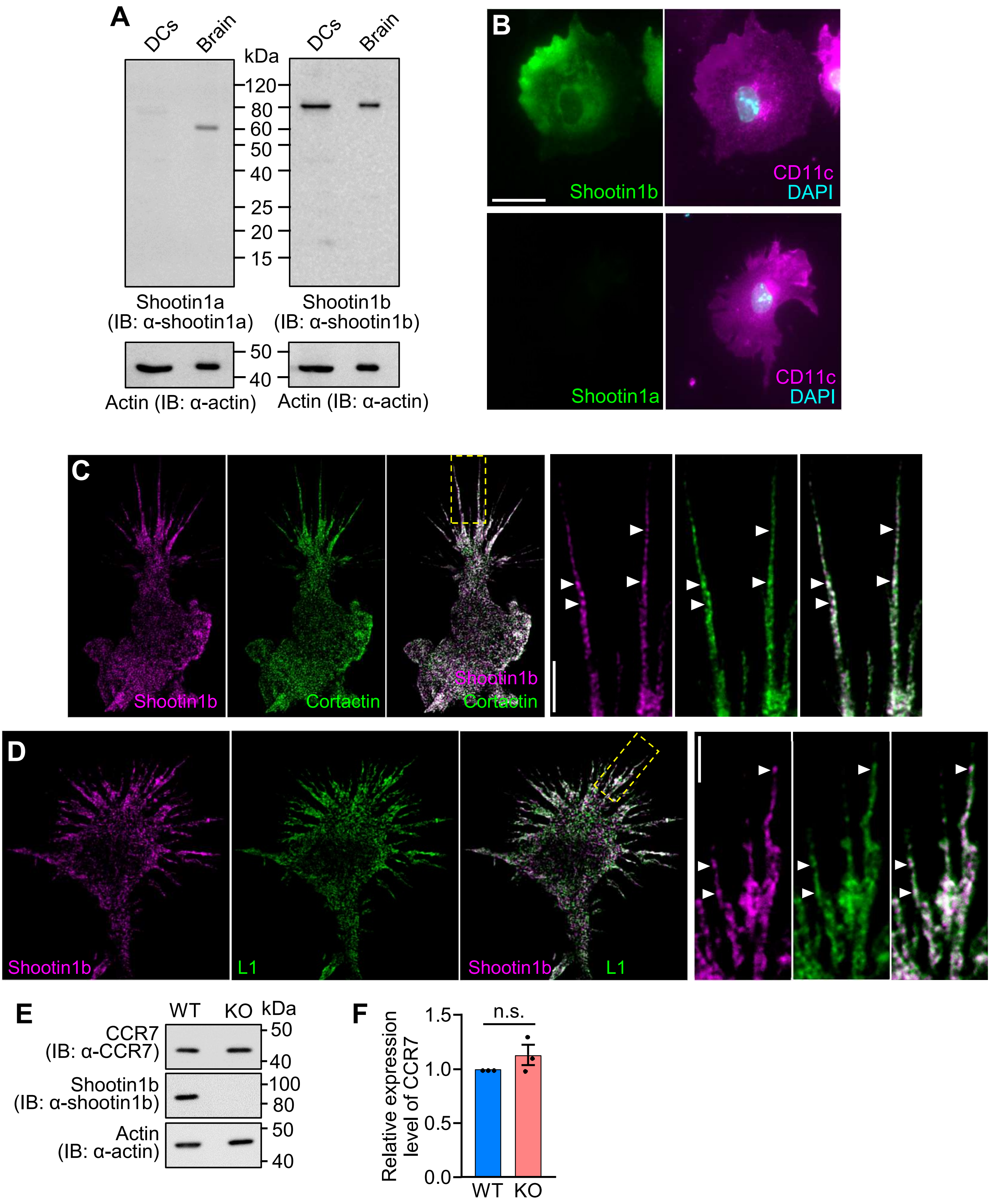

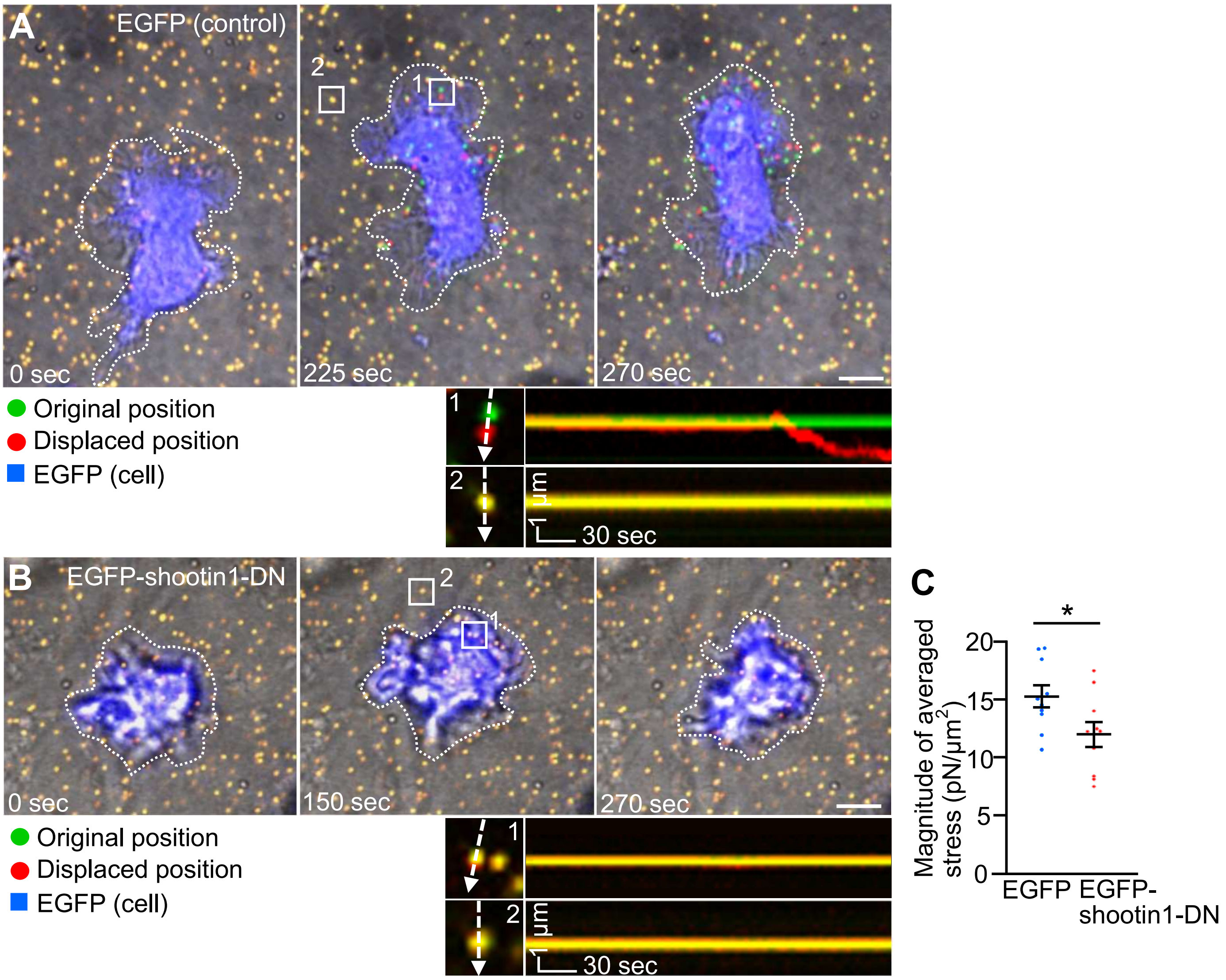

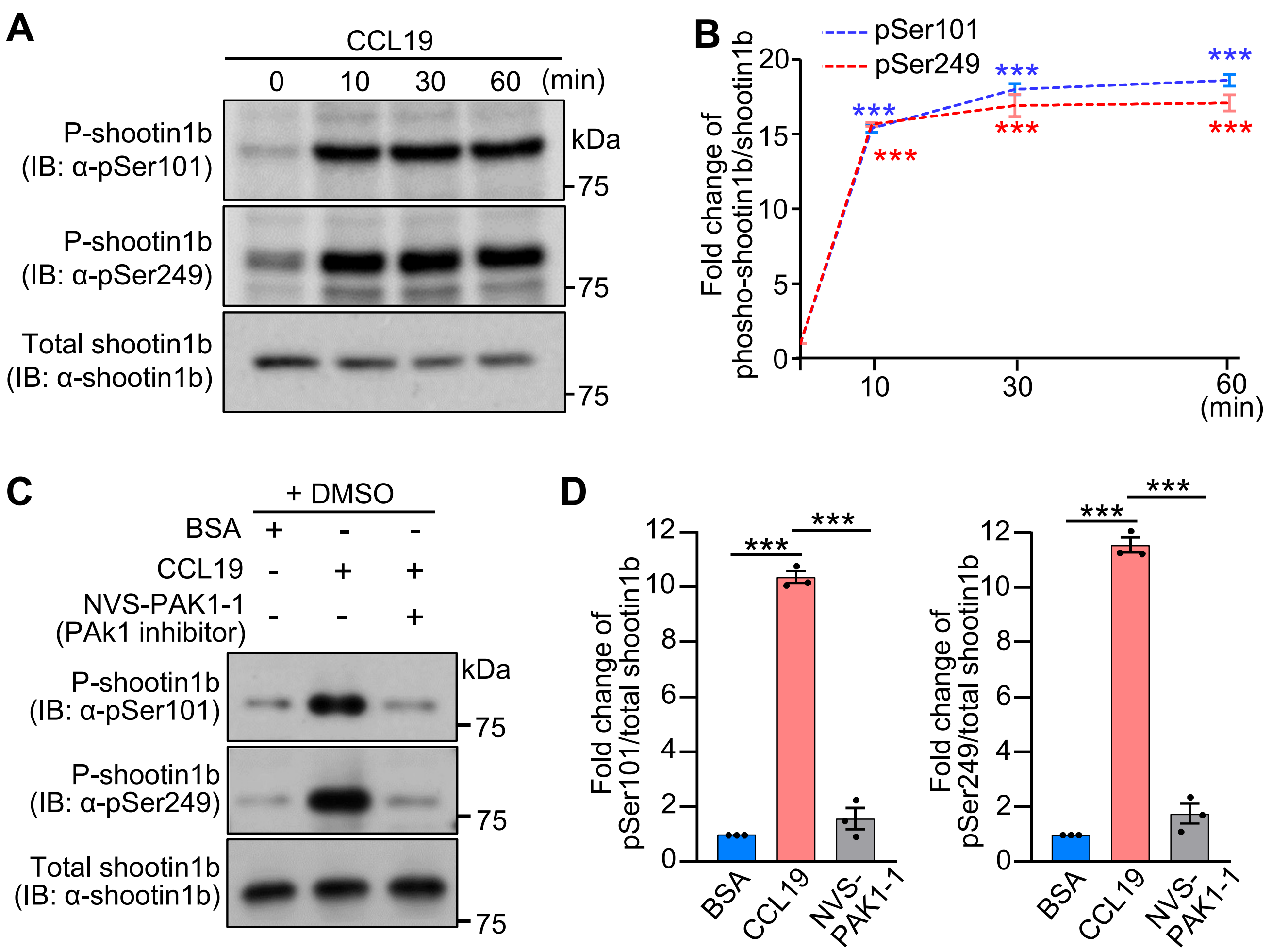

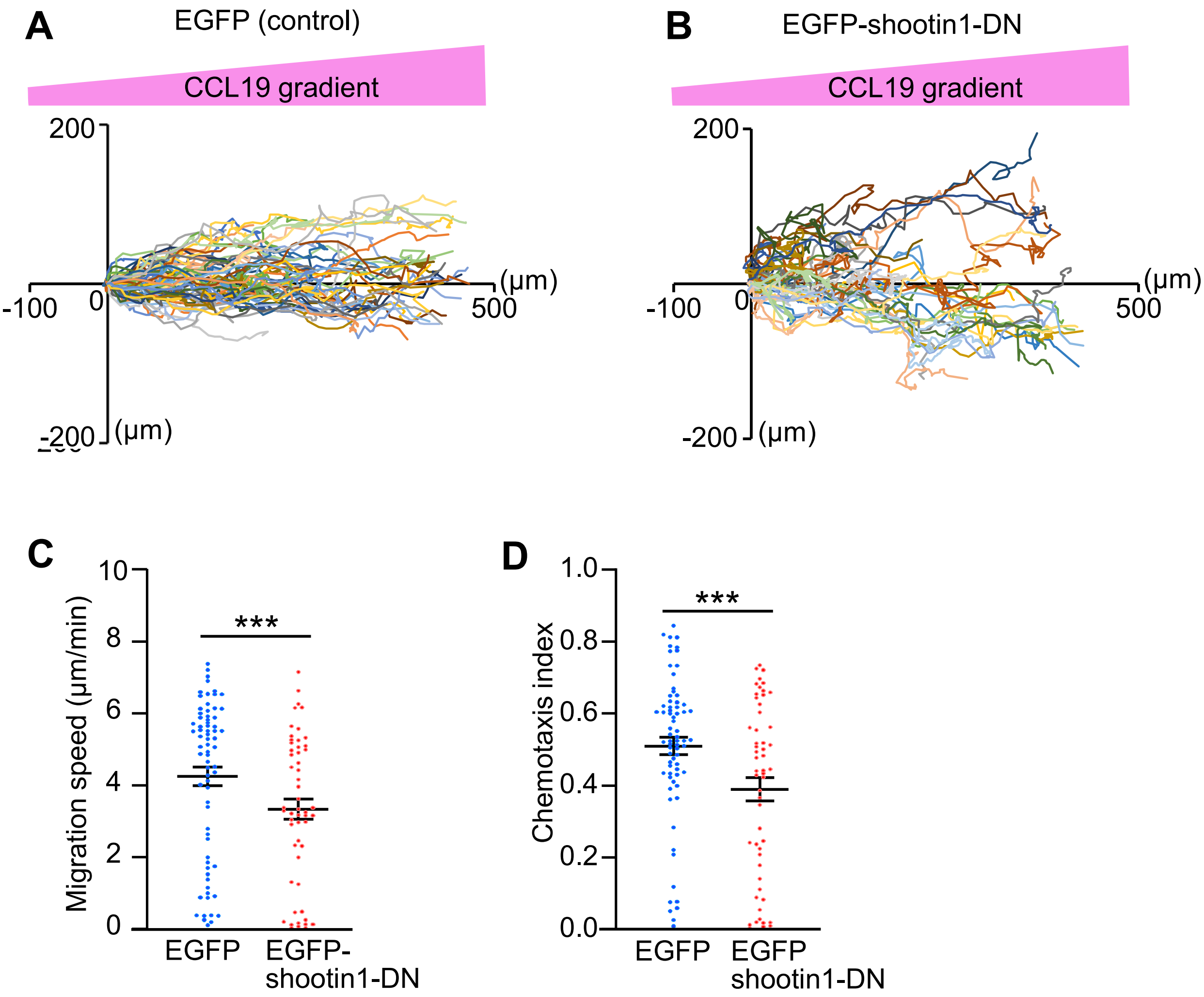

#### Figure S6 Baba et al.

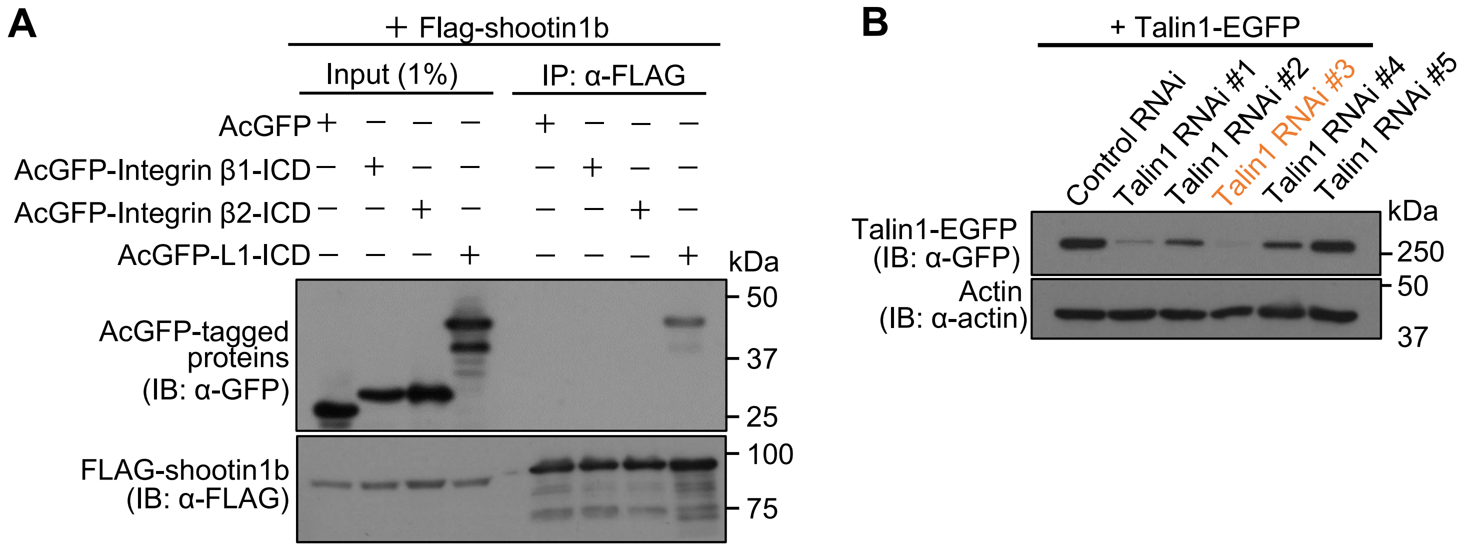

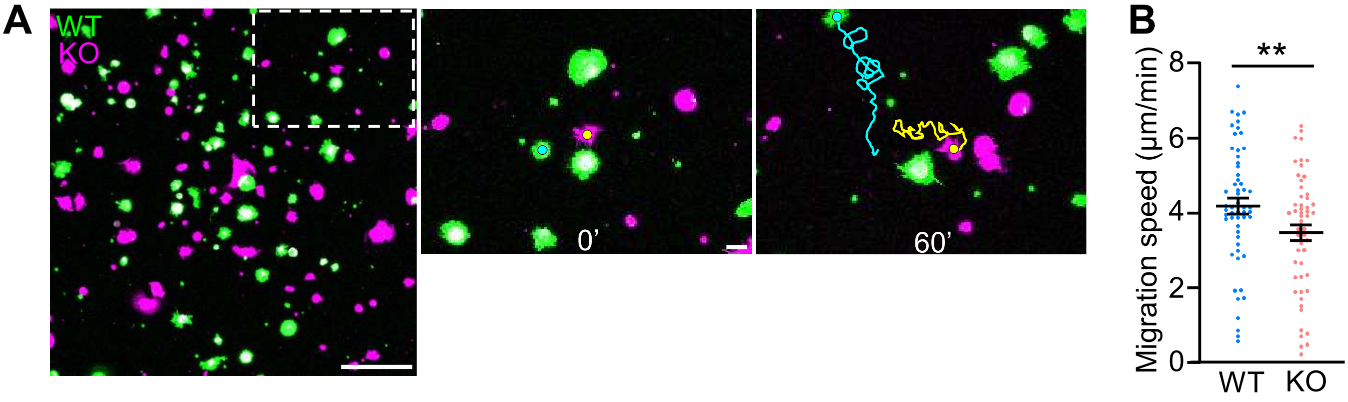

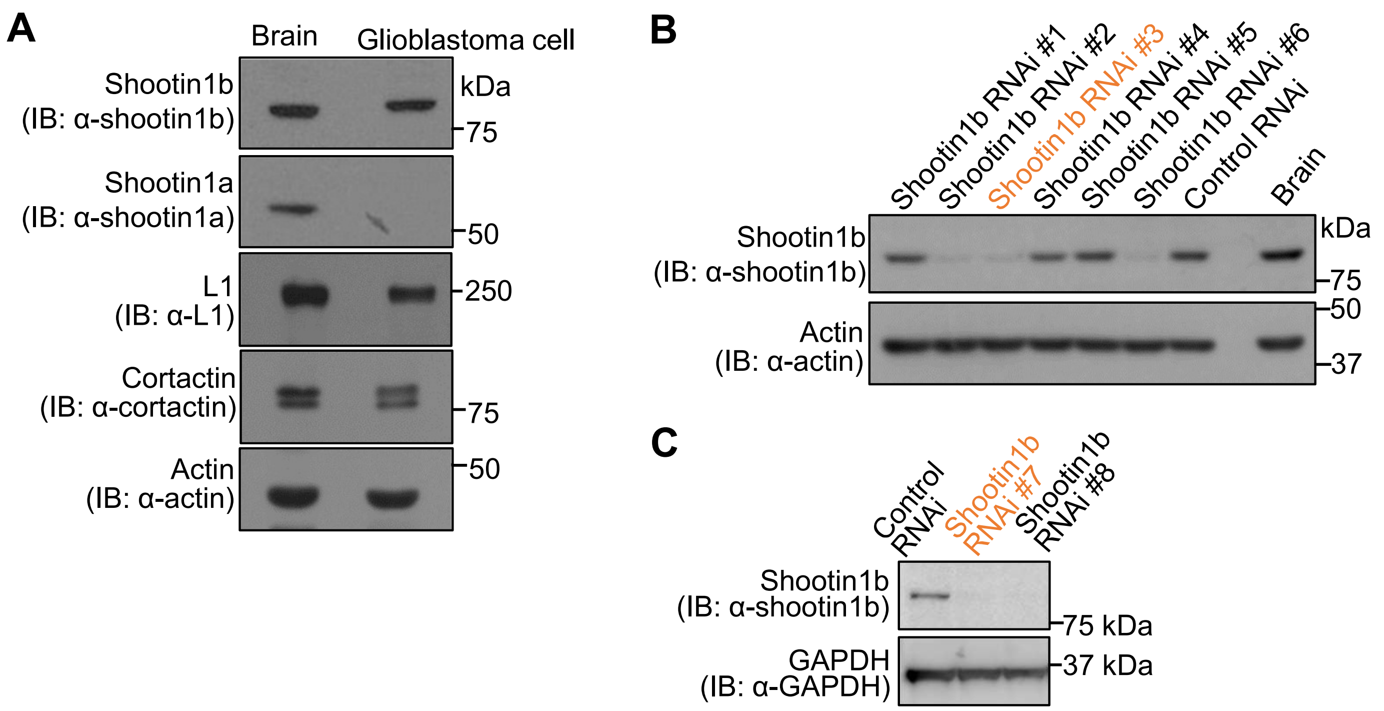

**Figure S9 Baba et al.**

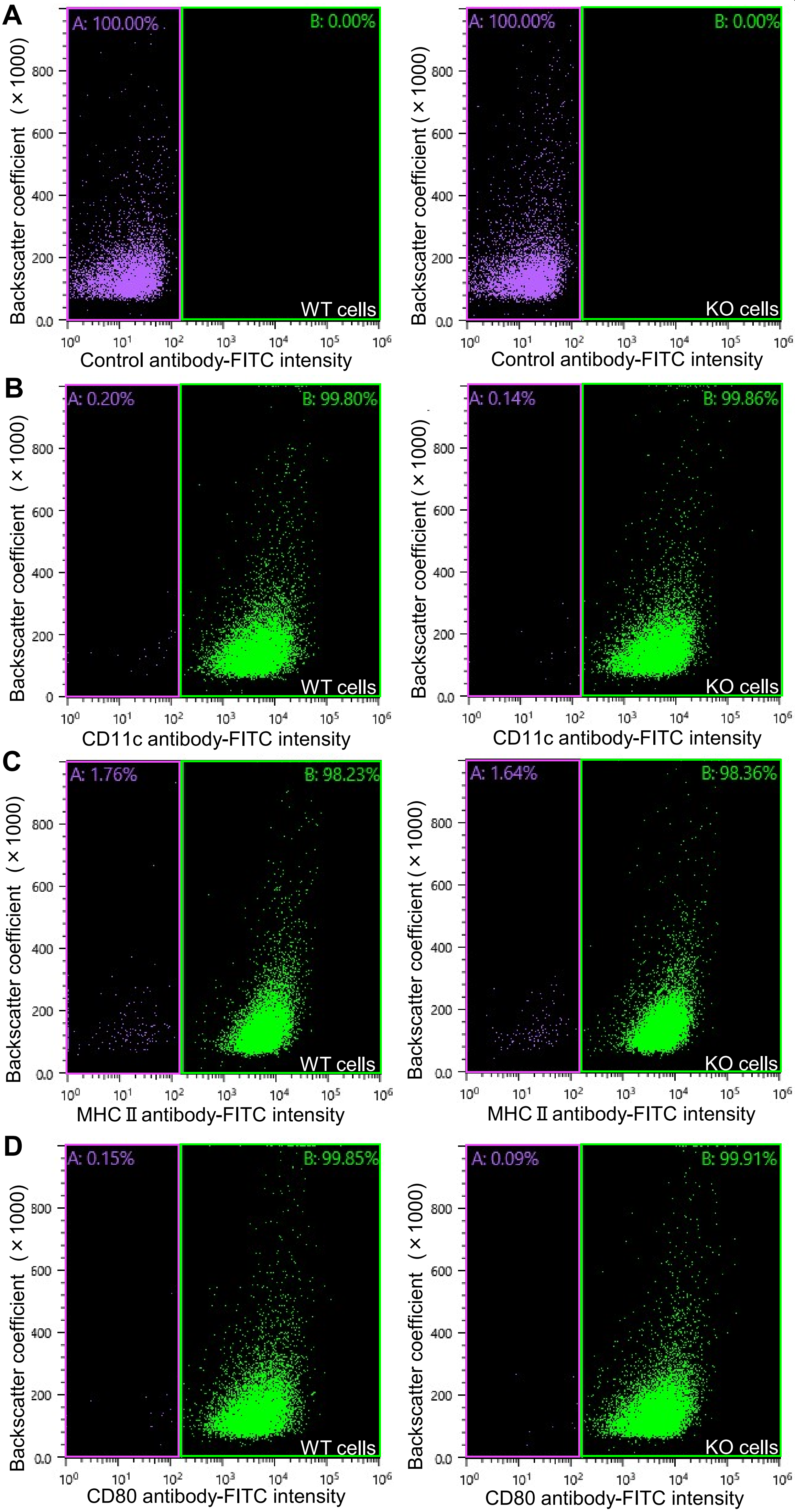
