## Supplementary Table S1 for "Weak and tunable adhesion-clutch drives rapid cell migration and glioblastoma motility"

Preparation of 1.5 mg /mL collagen gel + 10 % matrigel for chemotaxis assay

| Stock conc. | Volume (μL) | Final conc. |
|---|---|---|
| 10 ×MEM | 100 μL | 1×MEM |
| 1 M HEPES pH 7.4 | 10 μL | 10 mM HEPES |
| 1 M NaHCO$_3$ | 10 μL | 10 mM NaHCO3 |
| H$_2$O | 480 μL | |
| 100 % Matrigel | 100 μL | 10 % Matrigel |
| 5 mg /mL Collagen | 300 μL | 1.5 mg/mL collagen |
| Total volume | 1000 μL | |

Preparation of 1.5 mg /mL collagen gel + 10 % matrigel + CCL19 (20 or 200 ng/mL) for random migration assay in the bath application of CCL19

<table>
<tr><th>Stock conc.</th><th colspan="2">Volume (μL)</th><th colspan="2">Final conc.</th></tr>
<tr><td>10 ×MEM</td><td colspan="2">100 μL</td><td colspan="2">1×MEM</td></tr>
<tr><td>1 M HEPES pH 7.4</td><td colspan="2">10 μL</td><td colspan="2">10 mM HEPES</td></tr>
<tr><td>1 M NaHCO$_3$</td><td colspan="2">10 μL</td><td colspan="2">10 mM NaHCO3</td></tr>
<tr><td>H$_2$O</td><td>460 μL</td><td>478 μL</td><td colspan="2"></td></tr>
<tr><td>10 μg/ mL CCL19</td><td>20 μL</td><td>2 μL</td><td>200 ng/mL CCL19</td><td>20 ng/mL CCL19</td></tr>
<tr><td>100 % Matrigel</td><td colspan="2">100 μL</td><td colspan="2">10 % Matrigel</td></tr>
<tr><td>5mg /mL Collagen</td><td colspan="2">300 μL</td><td colspan="2">1.5 mg/mL Collagen</td></tr>
<tr><td>Total volume</td><td colspan="2">1000 μL</td><td colspan="2"></td></tr>
</table>

・10 ×MEM (Sigma, catalog number: M0275)
・1M HEPES, pH 7.4 (Nacalai, catalog number: 17557-94)
・Matrigel (Corning, 356231)
・5 mg/mL Collagen (IAC-50, catalog number: IAC-50)
・CCL19 (R&D system, catalog number: 440-M3-025)
