## Supplementary Table S2 for "Weak and tunable adhesion-clutch drives rapid cell migration and glioblastoma motility"

| Sample name | Oligonucleotide | Source |
|---|---|---|
| Genotyping Fw1 | 5’-CAGACTGCTACCCACTACCCCCTAC-3’ | (Baba et al., e Life. 2018) |
| Genotyping Rv1 | 5’-CCTAGAGCTGGACAGCGGATCTGAG-3’ | (Baba et al., e Life. 2018) |
| Genotyping Fw2 | 5’-CCCAGAAAGCGAAGGAACAAAGCTG-3’ | (Baba et al., e Life. 2018) |
| Genotyping Rv2 | 5’-ACCTTGCTCCTTCAAGCTGGTGATG-3’ | (Baba et al., e Life. 2018) |
| S101A Fw | 5’-AAAAGAATCGCCATGCTATACATG-3’ | This study |
| S101A Rv | 5’-CATGTATAGCATGGCGATTCTTTT-3’ | This study |
| S101D Fw | 5’- AAAAGAATCGACATGCTATACATG-3’ | This study |
| S101D Rv | 5’- CATGTATAGCATGTCGATTCTTTT-3’ | This study |
| S249A Fw | 5’- AAGAGACAAGCCCACCTTCTGCTG-3’ | This study |
| S249A Rv | 5’- CAGCAGAAGGTGGGCTTGTCTCTT-3’ | This study |
| S249D Fw | 5’- AAGAGACAAGACCACCTTCTGCTG-3’ | This study |
| S249D Rv | 5’- CAGCAGAAGGTGGTCTTGTCTCTT-3’ | This study |
| Integrin β1-ICD Fw | 5’-AAAGGATCCAAACTTTTAATGATAATTCAT-3’ | This study |
| Integrin β1-ICD Rv | 5’-AAAGTCGACTCATTTTCCCTCATACTTCG-3’ | This study |
| Integrin β2-ICD Fw | 5’-AAAGGATCCAAGGCCCTGACCCACCT-3’ | This study |
| Integrin β2-ICD Rv | 5’-AAAGTCGACCTAGCTTTCAGCAAACTTGGG-3’ | This study |
| Lifeact top | 5’- TCGAGATGGGTGTCGCAGATTTGATCAAGAAATTCGAAAGCATCTCAAAGGAAGAAGGG-3’ | This study |
| Lifeact bottom | 5’-GATCCCCTTCTTCCTTTGAGATGCTTTCGAATTTCTTGATCAAATCTGCGACACCCATC-3’ | This study |
| Shootin1b RNAi #1 Top | 5'-TGCTGCTTCATATTCGCCTATTGCTTGTTTGGCCACTGACTGACAAGCAATACGAATATGAAG-3' | This study |
| Shootin1b RNAi #1 Bottom | 5'-CCTGCTTCATATTCGTATTGCTTGTCTTGTCAGTCAGTGGCCAAAACAAGCAATAGGCGAATATGAAGC-3' | This study |
| Shootin1b RNAi #2 Top | 5'-TGCTGACACTTTCTCGACAAGTCTTGTTTTGGCCACTGAATGACAAGACTTGGAGAAAGTGCT-3' | This study |
| Shootin1b RNAi #2 Bottom | 5'-CCTGAGCACTTTCTCCAAGTCTCTTGTCAGTCAGTTGGCCAAAACAAGACTTGTCGAGAAAGTGCTC-3' | This study |
| Shootin1b RNAi #3 Top | 5'-TGCTGTATTACATCTGGTCCCAGCTTGTTTTGGCCACTGACTGACAAGCTGGGCAGATGTAATA-3' | This study |
| Shootin1b RNAi #3 Bottom | 5'-CCTGTATTACATCTGCCCAGCTTGTCAGTCAGTGCCCAAACAAGCTGGGACCAGATGTAATAC-3' | This study |
| Shootin1b RNAi #4 Top | 5'-TGCTGTCTTCTACAGCTAACATGGACGTTTTGGCCACTGACTGACGTCCATGTGCTGTAGAAGA-3' | This study |
| Shootin1b RNAi #4 Bottom | 5'-CCTGGTCTTCTACAGCACATGGACGTCAGTCAGTGGCCAAACGTCCATGTTAGCTGTATAATAC-3' | This study |
| Shootin1b RNAi #5 Top | 5'-TGCTGTTTCGAAGGTCCTTCTCCAGCGTTTGGCCACTGACTGACGCTGGAGAGACCTTCGAAA-3' | This study |
| Shootin1b RNAi #5 Bottom | 5'-CCTGTTTCGAAGGTCTCTCCAGCGTCAGTCAGTGGCCAAACGCTGGAGAAGGACCTTCGAAAC-3' | This study |
| Shootin1b RNAi #6 Top | 5'-TGCTGTGAAACGTAACCTTGGAACTTGTTTTGGCCACTGACTGACAAGTTCCTTCCAGTTACGTTTCA-3' | This study |
| Shootin1b RNAi #6 Bottom | 5'-CCTGTGAAACGTAACTGGAACTTGTCAGTCAGTGGCCAAAACAAGTTCCAAGGTTACGTTTCAC -3' | This study |
| Shootin1b RNAi #7 Top | 5’-CCGGTGGTCATAGAGGAAGTTAATTCTCGAGAATTAACTTCCTCTATGACCATTTTTG-3’ | This study |
| Shootin1b RNAi #7 Bottom | 5’-AATTCAAAAATGGTCATAGAGGAAGTTAATTCTCGAGAATTAACTTCCTCTATGACCA-3’ | This study |
| Shootin1b RNAi #8 Top | 5’-CCGGAAGACTTGTCGAGAAAGTGCTCTCGAGAGCACTTTCTCGACAAGTCTTTTTTTG-3’ | This study |
| Shootin1b RNAi #8 Bottom | 5’-AATTCAAAAAAAGACTTGTCGAGAAAGTGCTCTCGAGAGCACTTTCTCGACAAGTCTT-3’ | This study |
| Control scramble shRNA top | 5’-CCGGCCTAAGGTTAAGTCGCCCTCGCTCGAGCGAGGGCGACTTAACCTTAGGTTTTTG-3’ | This study |
| Control scramble shRNA bottom | 5’-AATTCAAAAACCTAAGGTTAAGTCGCCCTCGCTCGAGCGAGGGCGACTTAACCTTAGG-3’ | This study |
| Talin1 RNAi #1 Top | 5'-TGCTGAACCAGTGAATACTCATCATGGTTTTGGCCACTGACTGACCATGATGAATTCACTGGTT-3' | This study |
| Talin1 RNAi #1 Bottom | 5'-CCTGAACCAGTGAATTCATCATGGTCAGTCAGTGGCCAAAACCATGATGAGTATTCACTGGTTC-3' | This study |
| Talin1 RNAi #2 Top | 5'-TGCTGTTCTGTGGCGGCCTGATTTAAGTTTTGGCCACTGACTGACTTAAATCACCGCCACAGAA-3' | This study |
| Talin1 RNAi #2 Bottom | 5'-CCTGTTCTGTGGCGGTGATTTAAGTCAGTCAGTGGCCAAAACTTAAATCAGGCCGCCACAGAAC-3' | This study |
| Talin1 RNAi #3 Top | 5'-TGCTGTGTACTCGGTGTTTGATGTGAGTTTTGGCCACTGACTGACTCACATCACACCGAGTACA-3' | This study |
| Talin1 RNAi #3 Bottom | 5'-CCTGTGTACTCGGTGTGATGTGAGTCAGTCAGTGGCCAAAACTCACATCAAACACCGAGTACAC-3' | This study |
| Talin1 RNAi #4 Top | 5'-TGCTGTGACCTTGCAAGCTACCAGGAGTTTTGGCCACTGACTGACTCCTGGTATTGCAAGGTCA-3' | This study |
| Talin1 RNAi #4 Bottom | 5'-CCTGTGACCTTGCAATACCAGGAGTCAGTCAGTGGCCAAAACTCCTGGTAGCTTGCAAGGTCAC-3' | This study |
| Talin1 RNAi #5 Top | 5'-TGCTGTTACCAGGTTATCTGAGGCCCGTTTTGGCCACTGACTGACGGGCCTCATAACCTGGTAA-3' | This study |
| Talin1 RNAi #5 Bottom | 5'-CCTGTTACCAGGTTATGAGGCCCGTCAGTCAGTGGCCAAAACGGGCCTCAGATAACCTGGTAAC-3' | This study |
